# Circuit-selective cognitive vulnerability to environmental stress: multi-domain assessment of space radiation in both sexes reveals countermeasure trade-offs

**DOI:** 10.64898/2026.03.26.714605

**Authors:** Sheridan A. O’Connor, Pragatee Narain, Amishi Mahajan, Grace L. Bancroft, Harley A. Haas, Elise Wallen-Friedman, Shubha Vasisht, Hajime Takano, Frederico C. Kiffer, Amelia J. Eisch, Sanghee Yun

**Affiliations:** Department of Anesthesiology and Critical Care Medicine, The Children’s Hospital of Philadelphia (CHOP) Research Institute, Philadelphia, PA, USA; Department of Pathobiology, University of Pennsylvania School of Veterinary Medicine, Philadelphia, PA, USA; Department of University Laboratory Animal Resources,University of Pennsylvania, Philadelphia, PA, USA; University of Pennsylvania, School of Arts and Sciences, Philadelphia, PA, USA; Division of Neurology, The Children’s Hospital of Philadelphia (CHOP) Research Institute, Philadelphia, PA, USA; Department of Neuroscience, University of Pennsylvania Perelman School of Medicine, Philadelphia, PA, USA; Neuroscience Graduate Group, University of Pennsylvania Perelman School of Medicine, Philadelphia, PA, USA; Department of Anesthesiology and Critical Care Medicine, University of Pennsylvania Perelman School of Medicine, Philadelphia, PA, USA

## Abstract

Environmental stressors rarely affect just one brain circuit. Most studies assess single cognitive endpoints, obscuring whether vulnerabilities are global or circuit-selective and how effects distribute across interconnected systems. To address this, we used galactic cosmic radiation (GCR), a Mars mission-relevant stressor that disrupts the hippocampal-nucleus accumbens-prefrontal circuit. C57BL/6J mice received 33-ion GCR simulation (33-GCR, 0.75 Gy) or sham radiation with the Nrf2-activating compound CDDO-EA or vehicle, followed by multi-domain behavioral testing in both sexes. Under very high memory load, male Veh/33-GCR mice showed enhanced pattern separation compared to Veh/Sham males, an effect normalized by CDDO-EA. Female mice showed no radiation-induced changes in pattern separation but weighed 9-18% more than Veh/Sham females and had reduced locomotor activity. Reward-based learning differed by sex: males showed no changes, while female Veh/33-GCR mice displayed enhanced reward anticipation that was further increased by CDDO-EA alone, with both treatments contributing to elevated goal-tracking. For behavioral flexibility, CDDO-EA impaired reversal learning in males regardless of radiation, while 33-GCR impaired reversal learning in females regardless of CDDO-EA. Principal component analysis revealed that treatments disrupted specific circuit relationships while leaving others intact, consistent with selective rather than global cognitive effects. Fiber photometry showed enhanced dentate gyrus encoding activity in irradiated males under high memory load. Combined CDDO-EA/33-GCR selectively reduced dentate gyrus progenitors in females. Males and females showed distinct, circuit-selective vulnerability patterns, demonstrating that multi-domain, both-sex assessment is necessary to capture how stressors and interventions affect integrated brain function. CDDO-EA proved to be a double-edged sword: protecting one cognitive domain while impairing another, a trade-off invisible to single-endpoint assessment. This framework has immediate relevance for astronaut risk assessment and extends to any context where neuroprotective interventions are evaluated against environmental stressors.

## Introduction

Understanding how environmental stressors affect the brain requires assessing multiple cognitive domains simultaneously [1–4]. However, most studies examine single behavioral endpoints, obscuring how effects distribute across interconnected neural circuits [5,6]. Cognitive functions central to real-world performance, including spatial navigation, reward processing, and behavioral flexibility, rely on coordinated activity across the hippocampus, nucleus accumbens (NAc), and prefrontal cortex (PFC) [7–10]. These regions function as an integrated circuit; disruption at any node can cascade system-wide [11–17]. It remains poorly understood if environmental stressors produce global or circuit-selective vulnerabilities and if these patterns differ by sex [18,19].

Galactic cosmic radiation (GCR) is a tractable and timely model for investigating these questions. Astronauts on deep space missions beyond Earth’s magnetosphere face chronic GCR exposure, a complex mixture of high-energy particles with documented risks to cognitive performance and mission success [20–22]. GCR consists predominantly of protons (85%), helium nuclei (14%), and HZE ions (≤1%); particles such as ¹⁶O, ¹²C, ⁵⁶Fe, and ²⁸Si are notable for their high linear energy transfer which causes DNA damage, oxidative stress, and inflammation [20,23–25]. The CNS is highly vulnerable to these effects, with space radiation inducing persistent structural and functional alterations across multiple brain regions [26–30]. However, most ground-based studies have used single-ion exposures, limited behavioral endpoints, and male-only designs, a limitation shared broadly across neuroscience [24,31,32]. These approaches neither replicate deep space conditions nor capture how radiation affects integrated cognitive function in both sexes.

The hippocampal-NAc-PFC circuit supports the cognitive functions most relevant to both astronaut performance and everyday human cognition. The hippocampus provides spatial and contextual information, with the dentate gyrus (DG) specialized for pattern separation, the ability to form distinct representations of similar experiences [33,34]. Through projections to the NAc and PFC, the hippocampus integrates contextual information with reward value and executive control [35,36]. he NAc serves as a hub where hippocampal information converges with reward signals to guide motivated behavior [37–39]; Pavlovian autoshaping paradigms provide a well-validated measure of this function [40,41]. The PFC provides top-down executive control, receiving hippocampal input for context-guided decision-making while modulating NAc activity to regulate behavioral flexibility [35,42–46].

Within this circuit, the DG is a particularly vulnerable node. Adult-born granule neurons contribute to pattern separation and memory formation [47–49], but their high metabolic activity and low antioxidant capacity render them susceptible to oxidative damage [50,51]. DG dysfunction can disrupt downstream signaling to the NAc and PFC, with consequences for circuit-wide function [35]. All three regions show vulnerability to radiation-induced changes, including altered dopaminergic signaling, reduced dendritic complexity, and disrupted synaptic plasticity [26,52–57]. Prior work demonstrates that GCR impairs hippocampal-dependent learning and neurogenesis and induces neuroinflammation [58–64]. The relative vulnerability of the hippocampus, NAc, and PFC to multi-ion GCR tested within a single study remains unknown.

Despite this progress, critical knowledge gaps remain. Prior GCR studies examining hippocampal-dependent tasks have shown inconsistent effects (deficits and enhancements alike) depending on dose and paradigm [22,55,65]. Recent multi-ion studies demonstrate impairments in hippocampal long-term memory [58,62] and PFC-based attention [66], while single-ion work has shown effects on NAc and reward circuitry [56,67–73]. No study has assessed how multi-ion GCR affects the integrated hippocampal-NAc-PFC circuit across multiple cognitive domains, or whether effects are global versus domain-selective. Males and females may also show different vulnerability patterns [74]. Sex differences in space radiation effects remain poorly characterized [22,65,75,76], and most studies have tested only males. Given that disruption at any circuit node can cascade system-wide [11–17], understanding multi-domain cognitive outcomes in both sexes is essential for astronaut risk assessment [20,77] and for the broader question of how environmental stressors affect integrated brain function.

No adequate shielding currently exists for deep space radiation, making pharmacological countermeasures essential [76,78,79]. Because GCR-induced oxidative damage and inflammation can compromise multiple nodes within the hippocampal-NAc-PFC circuit [66], effective countermeasures must provide broad neuroprotection. CDDO-EA (2-cyano-3,12-dioxooleana-1,9(11)-dien-28-oic acid ethyl amide) is a candidate countermeasure: a synthetic triterpenoid that activates the nuclear factor erythroid 2-related factor 2 (Nrf2) pathway [80,81]. Nrf2 is a master regulator of cellular antioxidant responses, inducing cytoprotective gene expression [82,83]. CDDO-EA and related triterpenoids have demonstrated neuroprotective effects in injury and neurological disorder models, with Nrf2 activators mitigating radiation-induced damage across multiple organ systems [83–89]. The structurally related Nrf2-activating compound omaveloxolone (Skyclarys) recently received FDA approval for treating Friedreich’s ataxia [90], supporting the translational potential of this compound class. Our laboratory demonstrated that CDDO-EA provides cognitive protection in female mice exposed to 33-GCR, including enhanced pattern separation and improved reversal learning [91]. However, that study used a single behavioral endpoint in males [92], leaving open whether domain-specific effects were missed. This raises the question: does CDDO-EA protect uniformly, or does protection in one domain come at the cost of impairment in another?

Here we investigated how 33-ion GCR exposure affects cognition across multiple domains in male and female mice, and whether CDDO-EA provides broad neuroprotection or domain-selective effects. Based on prior work [22,65,75], we hypothesized that 33-GCR would produce domain-specific rather than global cognitive effects that differ between sexes, with hippocampal-dependent functions showing particular vulnerability in males [93,94] and metabolic or locomotor effects predominating in females [22,65,75]. We further hypothesized that CDDO-EA would show domain-dependent rather than uniform protection, with potential trade-offs across circuit nodes [91,92]. To test these hypotheses, we assessed four cognitive domains (hippocampal-dependent pattern separation, reward-based learning, behavioral flexibility, and anxiety-related behavior), alongside body weight, locomotor activity, hippocampal neurogenesis markers, and neural activity over 7.25 months post-irradiation (post-IRR). Neural activity and cellular markers were assessed at later timepoints to determine whether radiation-induced alterations persist beyond the behavioral testing window. Our findings reveal domain-selective rather than global effects, with distinct vulnerability patterns in males and females and countermeasure trade-offs invisible to single-endpoint assessment. This framework extends beyond space radiation to any context where neuroprotective interventions are evaluated against environmental stressors.

## Results

Male and female C57BL/6J mice received 33-ion GCR simulation (0.75 Gy) or sham irradiation with concurrent CDDO-EA or vehicle treatment, followed by multi-domain behavioral testing over 7.25 months post-IRR (**Fig. 1**).

**Fig. 1.**
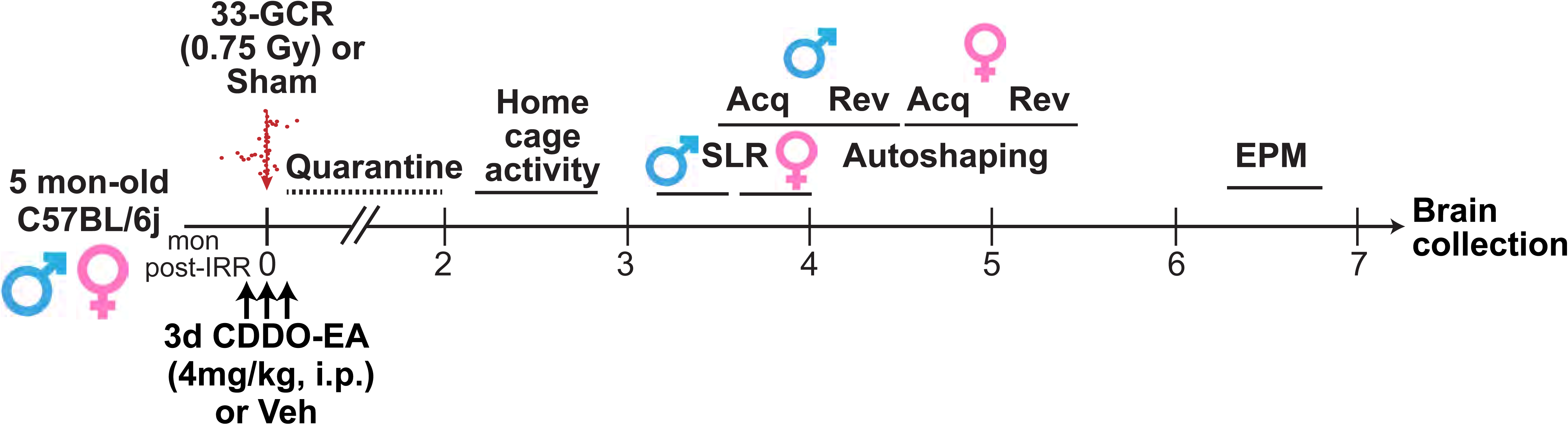
Experimental timeline showing treatment and behavioral testing schedule over 7.25 months. Male and female C57BL/6J mice (n=47/sex) received 2-cyano-3,12-dioxooleana-1,9-dien-28-oic acid-ethylamide (CDDO-EA; 4 mg/kg, i.p.) or vehicle (Veh) for 3 days surrounding administration of whole-body 33-ion galactic cosmic radiation (33-GCR, 0.75 Gy) or sham irradiation at 4.5–5 months of age. Behavioral assessments included home cage activity, spontaneous location recognition (SLR), touchscreen-based Pavlovian learning, and elevated plus maze testing over 7.25 months post-irradiation. Abbreviations: 33-GCR, 33-ion galactic cosmic radiation; CDDO-EA, 2-cyano-3,12-dioxooleana-1,9-dien-28-oic acid-ethylamide; EPM, elevated plus maze; IRR, irradiation; SLR, spontaneous location recognition; Veh, vehicle.

### 33-GCR enhances pattern separation in males under very high memory load; CDDO-EA normalizes this effect

Hippocampal-dependent cognition was assessed using the spontaneous location recognition (SLR) task with varying memory loads (**Fig. 2A**). Task difficulty increased as objects were placed closer together: dissimilar (d-SLR, low load), similar (s-SLR, high load), and extra-similar (xs-SLR, very high load). Males began testing at 3.25 months post-IRR.

**Fig. 2.**
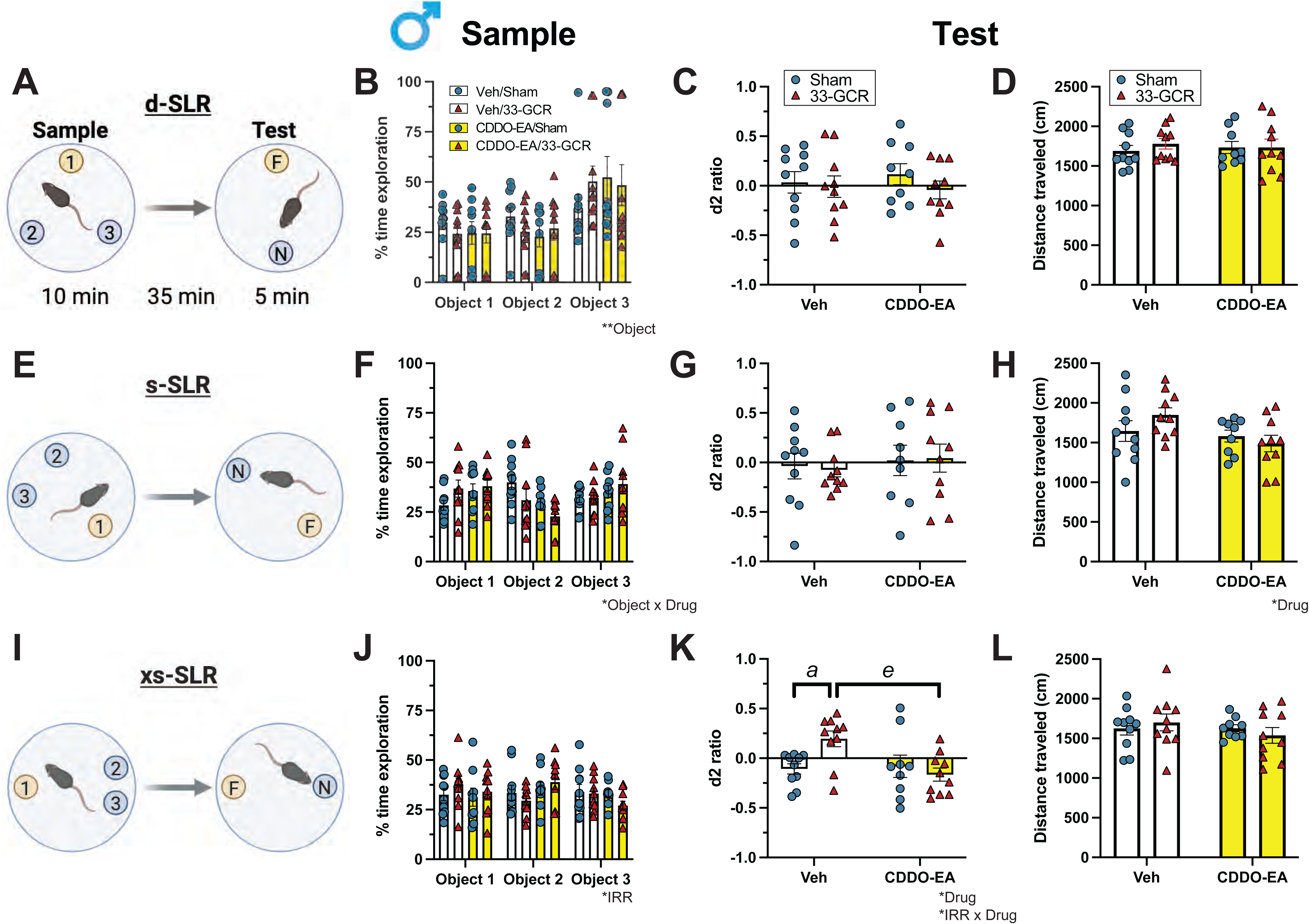
33-GCR enhances male pattern separation under very high memory load; CDDO-EA normalizes this. **(A)** Schematic of the d-SLR (dissimilar: low memory load) task. Mice explored three identical objects (objects 2 and 3, 108° apart) during the Sample phase (10 min), then after a 35-min delay explored two objects—one at a familiar location [F] and one at a novel location [N]—during the Test phase (5 min). **(B)** d-SLR Sample: percent time exploring each object location. **(C)** d-SLR Test: discrimination index (d2 ratio). **(D)** d-SLR Test: total distance traveled (cm). **(E)** Schematic of the s-SLR (similar: high memory load) task (objects 2 and 3, 72° apart). **(F)** s-SLR Sample: percent time exploring each location. **(G)** s-SLR Test: d2 ratio. (H) s-SLR Test: total distance traveled. **(I)** Schematic of the xs-SLR (extra-similar: very high memory load) task (objects 2 and 3, 36° apart). **(J)** xs-SLR Sample: percent time exploring each location. **(K)** xs-SLR Test: d2 ratio. **(L)** xs-SLR Test: total distance traveled. Data are presented as mean ± SEM (n=9–10/group). Statistical analysis: 3-way RM ANOVA (IRR × Drug × Object) in **B, F**, J; 2-way ANOVA (IRR × Drug) in **C–D**, **G–H**, **K–L**. *p<0.05, **p<0.01. Post-hoc: Veh/Sham vs Veh/33-GCR, ᵃ p<0.05; Veh/33-GCR vs CDDO-EA/33-GCR, ᵉ p<0.05. Complete subject numbers and detailed statistical analyses are provided in **S1 Table**.

Under low and high memory loads (d-SLR, s-SLR), all groups showed similar object exploration during the Sample phase and comparable discrimination and locomotion during the Test phase (**Fig. 2B-H**, **S1 Table**; all post-hoc p>0.05).

Group differences emerged under very high memory load (xs-SLR). During the Sample phase, males in all groups explored objects similarly (**Fig. 2J**; 3-way RM ANOVA, IRR p<0.05; all post-hoc p>0.05). During the Test phase, Veh/33-GCR males showed a positive d2 ratio (0.20 ± 0.09) while Veh/Sham males showed a negative d2 ratio (−0.11 ± 0.10), indicating successful discrimination of the novel location in irradiated but not control males (**Fig. 2K**; 2-way ANOVA, Drug p<0.05, IRR×Drug p<0.05; post-hoc Veh/Sham vs Veh/33-GCR, p<0.05). CDDO-EA prevented this radiation-induced enhancement: CDDO-EA/33-GCR males showed a negative d2 ratio (−0.17 ± 0.08), similar to Veh/Sham controls and less than Veh/33-GCR males (post-hoc p<0.05). Locomotion did not differ among groups (**Fig. 2L**; all p>0.05).

In sum, 33-GCR enhanced pattern separation in males specifically under very high cognitive demand, an effect normalized by CDDO-EA.

### Female mice show no radiation-induced changes in pattern separation despite reduced locomotion during testing

In contrast to males, female mice showed no group differences in pattern separation at any memory load. Females began testing at 3.75 months post-IRR.

Across all three memory loads, female groups showed similar object exploration during the Sample phase and comparable discrimination during the Test phase (**Fig. 3B-C, F-G, J-K, S1 Table**; all post-hoc p>0.05).

**Fig. 3.**
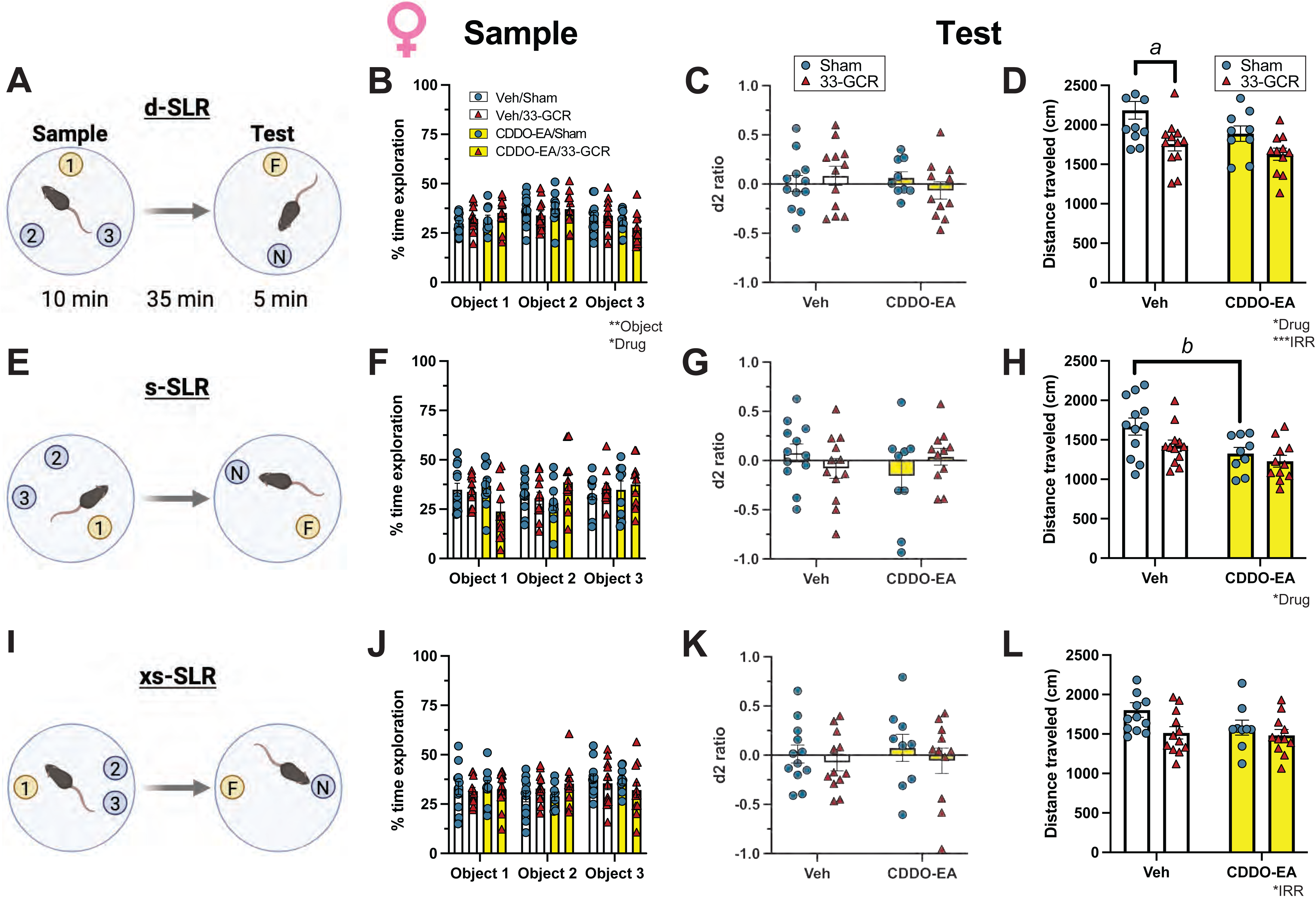
Female mice show no radiation-induced pattern separation changes despite reduced locomotion. **(A)** Schematic of the d-SLR task. **(B)** d-SLR Sample: percent time exploring each object location. **(C)** d-SLR Test: discrimination index (d2 ratio). **(D)** d-SLR Test: total distance traveled (cm). **(E)** Schematic of the s-SLR task. **(F)** s-SLR Sample: percent time exploring each location. **(G)** s-SLR Test: d2 ratio. **(H)** s-SLR Test: total distance traveled. **(I)** Schematic of the xs-SLR task. **(J)** xs-SLR Sample: percent time exploring each location. **(K)** xs-SLR Test: d2 ratio. **(L)** xs-SLR Test: total distance traveled. Data are presented as mean ± SEM (n=9–12/group). Statistical analysis: 3-way RM ANOVA (IRR × Drug × Object) in **B, F, J**; 2-way ANOVA (IRR × Drug) in **C–D, G–H, K–L**. *p<0.05, **p<0.01, ***p<0.001. Post-hoc: Veh/Sham vs Veh/33-GCR, ᵃ p<0.05; Veh/Sham vs CDDO-EA/Sham, ᵇ p<0.05. Complete subject numbers and detailed statistical analyses are provided in **S1 Table**.

However, locomotion during SLR testing differed among female groups. Under d-SLR, Veh/33-GCR females traveled 19.6% shorter distances during the Test phase than Veh/Sham females (**Fig. 3D**; 2-way ANOVA, IRR p<0.01, Drug p<0.05; post-hoc p<0.05). Under s-SLR, CDDO-EA/Sham females traveled 20.6% shorter distances than Veh/Sham females (**Fig. 3H**; Drug p<0.01; post-hoc p<0.05). Under xs-SLR, IRR affected locomotion overall but no post-hoc differences reached significance (**Fig. 3L**; IRR p<0.05; all post-hoc p>0.05).

Thus, neither 33-GCR nor CDDO-EA affected pattern separation in females, despite both treatments independently reducing locomotion during testing.

### 33-GCR produces physiological effects in females that dissociate from cognitive outcomes

Since females showed reduced locomotion during SLR testing without cognitive impairment (**Fig. 3**), we assessed body weight in both sexes throughout the 7.25-month experiment and measured home cage locomotor activity at 2.25 months post-IRR to determine whether 33-GCR and CDDO-EA produced systemic physiological effects.

All male mice gained weight progressively throughout the experiment, with no weight differences among the four groups (**Fig. 4A**, **S1 Table**; male: 3-way mixed effect RM ANOVA, Time F(6,258)=353.618, p<0.001; all other main effects and interactions p>0.05). Female mice also gained weight but, unlike males, showed group differences in weight trajectory (**Fig. 4B**, **S1 Table**; female: 3-way RM ANOVA, Time F(6,258)=250.06, p<0.001, IRR F(1,43)=6.28, p<0.05; Drug × Time F(6,258)=3.25, p<0.01, IRR × Time F(6,258)=12.13, p<0.001). At 1 month post-IRR, CDDO-EA/Sham females transiently weighed 8.6% more than Veh/Sham females (post-hoc p<0.01). From 2 months post-IRR through study completion, Veh/33-GCR females weighed 9.3-18.3% more than Veh/Sham females (post-hoc p<0.05 at months 2-4; p<0.01 at months 6-7). This weight difference was present before the onset of appetitive touchscreen testing (months 2-3, 9.3-11.2% increase) and became more pronounced during the food-restricted touchscreen period (months 4-7, 13.4-18.3% increase), suggesting that caloric restriction may have amplified rather than driven the effect.

**Fig. 4.**
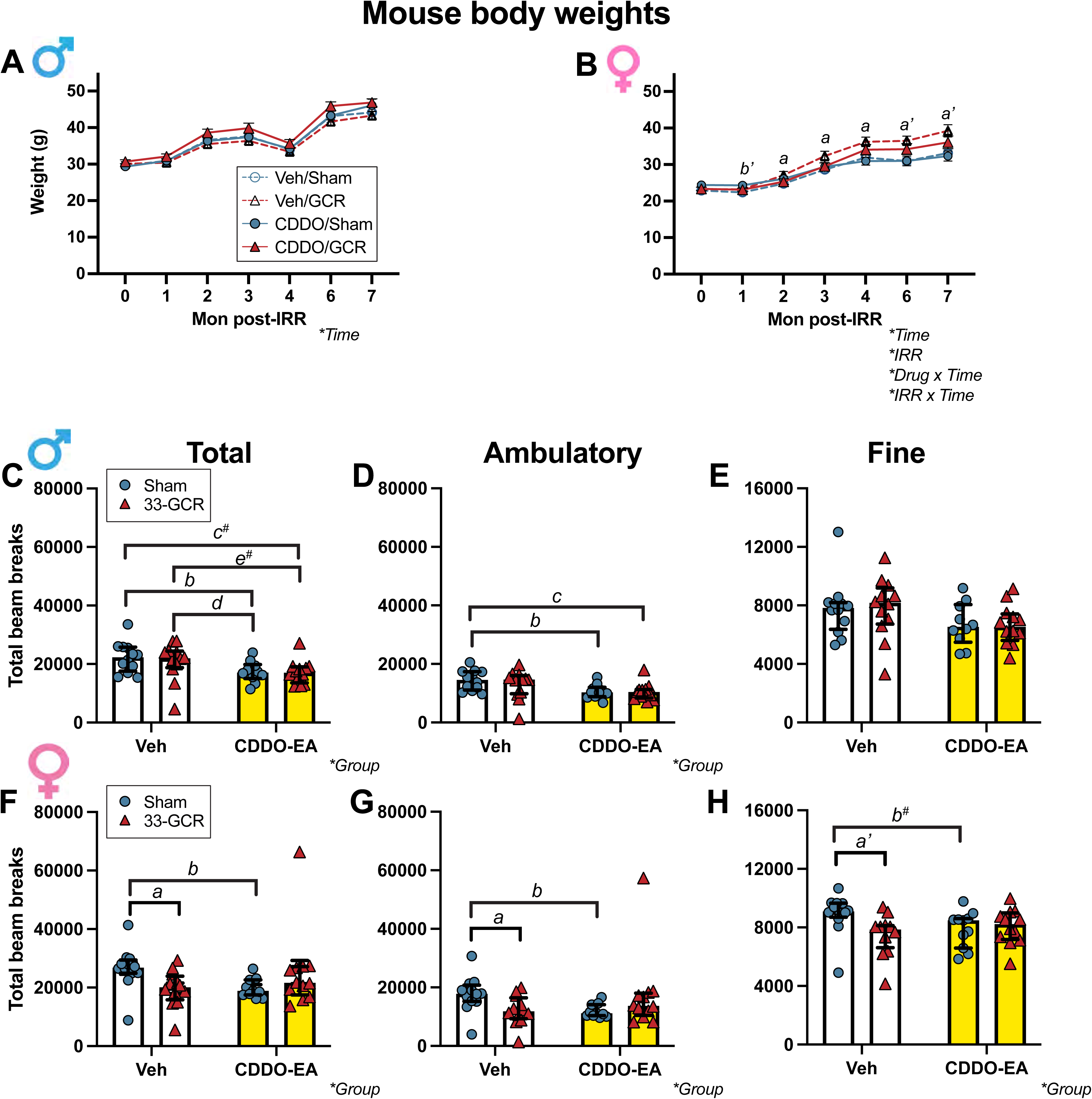
33-GCR produces physiological effects in females that dissociate from cognitive outcomes. **(A–B)** Body weights recorded monthly throughout the experiment in male **(A)** and female **(B)** mice (n=11–12/group). **(C–E)** Male home cage locomotor activity measured over 18 hours (4pm–10am) via 4×8 photobeam breaks in the xy-plane at 2.25 months post-IRR: total **(C)**, ambulatory **(D)**, and fine **(E)** beam breaks. **(F–H)** Female home cage locomotor activity: total **(F)**, ambulatory **(G)**, and fine **(H)** beam breaks (n=10–12/group). Body weight data are presented as mean ± SEM; locomotor data are presented as median ± IQR. Statistical analysis: mixed-effects 3-way RM ANOVA in **A**; 3-way RM ANOVA in **B**; Kruskal-Wallis test with Dunn’s post-hoc comparisons in **C–H**. Main effect and/or interaction denoted by *p<0.05. Post-hoc comparisons for body weight: Veh/Sham vs Veh/33-GCR, ᵃ p<0.05, ᵃ’ p<0.01; Veh/Sham vs CDDO-EA/Sham, ᵇ’ p<0.01 **(B)**. Post-hoc comparisons for locomotor activity: Veh/Sham vs Veh/33-GCR, ᵃ p<0.05, ᵃ’ p<0.01; Veh/Sham vs CDDO-EA/Sham, ᵇ p<0.05; Veh/33-GCR vs CDDO-EA/Sham, ᵈ p<0.05; # 0.05<p<0.08. Complete subject numbers and detailed statistical analyses are provided in **S1 Table**.

Home cage locomotor activity, measured over 18 hours including 12 hours of the dark cycle, was consistent with the locomotion differences observed during SLR testing. In males, Veh/Sham and Veh/33-GCR mice made a similar number of total beam breaks, indicating no effect of radiation alone on home cage activity. However, CDDO-EA/Sham males made 21.9% fewer total beam breaks compared to Veh/Sham males (**Fig. 4C, S1 Fig., S1 Table**; Kruskal-Wallis p<0.05; post-hoc Veh/Sham vs. CDDO-EA/Sham, p<0.05). This reduction was driven by 26.2% less ambulatory movement in CDDO-EA/Sham males compared to Veh/Sham males (**Fig. 4D**; Kruskal-Wallis p<0.05; post-hoc p<0.05), while fine movement did not differ among groups (**Fig. 4E**; Kruskal-Wallis p>0.05). Veh/33-GCR males also made fewer total and ambulatory beam breaks compared to CDDO-EA/Sham males (**Fig. 4C-D**; post-hoc p<0.05), consistent with CDDO-EA reducing activity regardless of radiation status.

In females, both 33-GCR and CDDO-EA independently reduced home cage locomotor activity. Veh/33-GCR females made 26.0% fewer total beam breaks than Veh/Sham females (**Fig. 4F**, **S1 Table**; Kruskal-Wallis p<0.05; post-hoc p<0.05), with 31.1% fewer ambulatory beam breaks (**Fig. 4G**; post-hoc p<0.05) and 15.9% fewer fine movement beam breaks (**Fig. 4H**; post-hoc p<0.01). Similarly, CDDO-EA/Sham females made 24.7% fewer total beam breaks than Veh/Sham females (**Fig. 4F**; post-hoc p<0.05), with 31.5% fewer ambulatory beam breaks (**Fig. 4G**; post-hoc p<0.05) and a trend toward 12.9% fewer fine movement beam breaks (**Fig. 4H**; post-hoc p=0.06).

These data show that 33-GCR exposure produced sustained weight gain and reduced home cage activity in female mice, while CDDO-EA reduced home cage activity in both sexes. These physiological effects dissociated from hippocampal-dependent cognition: females showed no radiation-induced pattern separation deficits despite pronounced weight and activity changes, while males showed radiation-enhanced pattern separation without corresponding weight or home cage activity differences.

### 33-GCR and CDDO-EA independently enhance goal tracking in females; males show no treatment effects on reward-based learning

We next examined whether 33-GCR affected reward-based learning, given its distinct effects on hippocampal cognition in males versus physiological outcomes in females. Mice were tested on touchscreen-based Pavlovian autoshaping over 11 days (**Fig. 5A**). This paradigm measures goal tracking (approaches to the reward magazine during CS+ presentation) and sign tracking (preferential approach to CS+ over CS−).

**Fig. 5.**
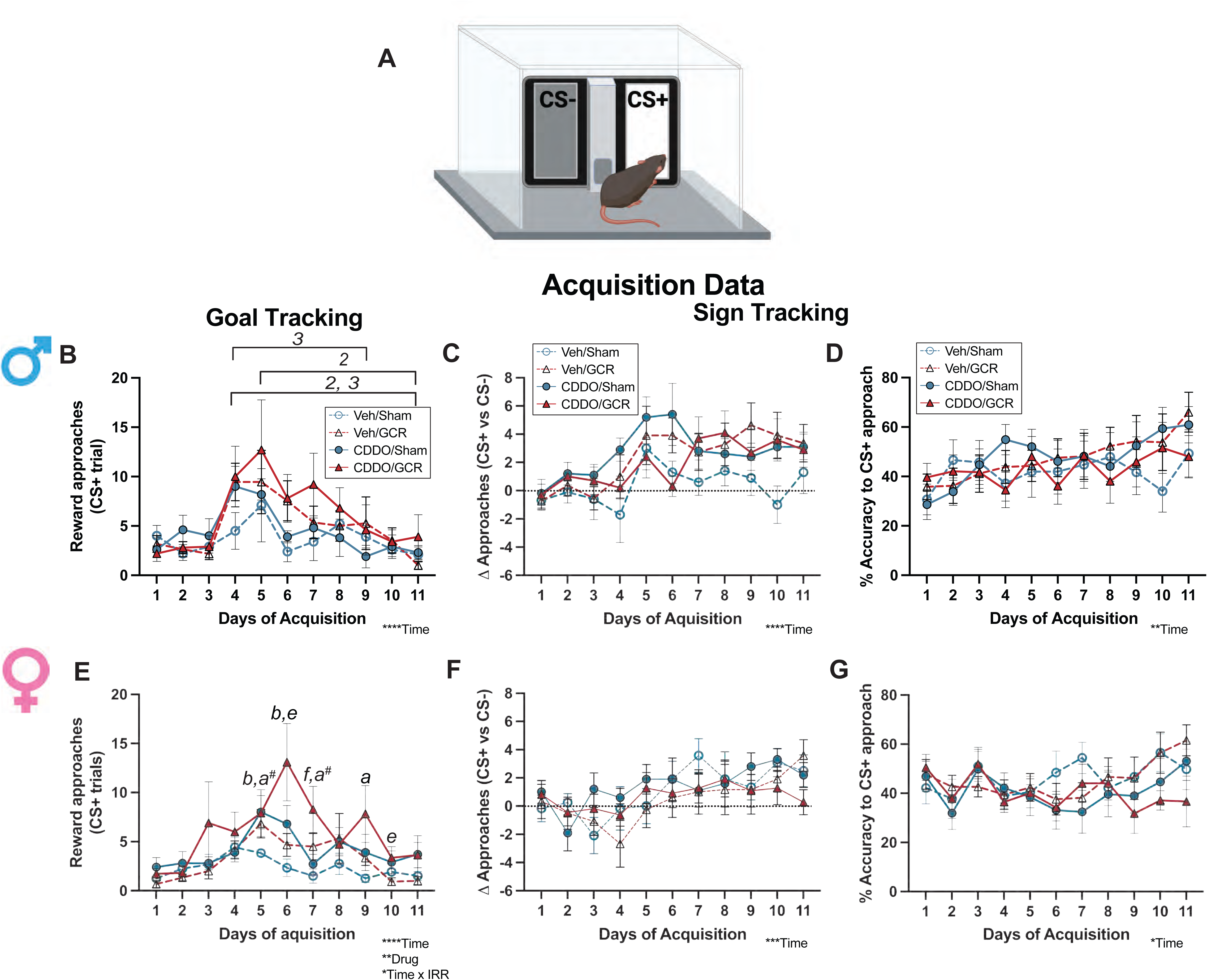
33-GCR and CDDO-EA independently enhance female goal-tracking; males are unaffected. **(A)** Schematic of the autoshaping task. During acquisition, a conditioned stimulus (CS+, illuminated panel) is paired with reward delivery (strawberry milkshake). Mice learn the CS-reward association over 11 days. **(B, E)** Goal-tracking: approaches to the reward magazine during CS+ presentation in males **(B)** and females **(E)**. **(C, F)** Sign-tracking: difference in approaches to CS+ vs CS− in males **(C)** and females **(F)**. **(D, G)** Accuracy: percent CS+ approaches out of total CS+ trials in males **(D)** and females **(G)**. Data are presented as mean ± SEM (n=10–12/group). Statistical analysis: 3-way RM ANOVA (IRR × Drug × Time). *p<0.05, **p<0.01, ***p<0.001, ****p<0.0001. Post-hoc: timepoints in Veh/GCR, ² p<0.05; timepoints in CDDO-EA/Sham, ³ p<0.05; Veh/Sham vs Veh/33-GCR, ᵃ p<0.05; Veh/Sham vs CDDO-EA/Sham, ᵇ p<0.05; Veh/33-GCR vs CDDO-EA/33-GCR, ᵉ p<0.05; CDDO-EA/Sham vs CDDO-EA/33-GCR, ᶠ p<0.05; # 0.05<p<0.08. Complete subject numbers and detailed statistical analyses are provided in **S1 Table**.

In males, all groups increased reward magazine approaches over acquisition days (**Fig. 5B**, **S1 Table**; 3-way RM ANOVA, Time p<0.0001). Within-group increases from early to late acquisition were confirmed in Veh/33-GCR and CDDO-EA/Sham groups (Days 4-5 vs. Days 9 or 11; post-hoc: Veh/33-GCR Day 4 vs Day 11 and Day 5 vs. Day 11, p<0.05; CDDO-EA/Sham Day 4 vs Day 9 and Day 4 vs Day 11, p<0.05), with similar numerical trends in Veh/Sham and CDDO-EA/33-GCR groups that did not reach significance. All male groups also progressively increased their preferential approach to CS+ over CS− (**Fig. 5C**; Time p<0.0001) and showed similar accuracy in approaching CS+ (**Fig. 5D**; Time p<0.001). Thus, neither 33-GCR nor CDDO-EA affected Pavlovian learning in males.

In contrast, female mice showed treatment-dependent differences in goal tracking (**Fig. 5E**, **S1 Table**; 3-way RM ANOVA, Time p<0.0001, Drug p<0.01, Time×IRR p<0.05). Veh/33-GCR females made more reward magazine approaches than Veh/Sham females on Day 9 (167% more; post-hoc p<0.05), with similar trends on Day 5 (78% more; p=0.056) and Day 7 (200% more; p=0.062). CDDO-EA also independently increased goal tracking: CDDO-EA/Sham females made 109% (Day 5) and 191% (Day 6) more approaches than Veh/Sham females (post-hoc p<0.05). CDDO-EA/33-GCR females also made 181% (Day 6) and 267% (Day 10) more approaches than Veh/33-GCR females (post-hoc p<0.05). These combined effects suggest additive influences on reward anticipation, with each treatment independently contributing to enhanced goal-tracking. Despite these goal tracking differences, all female groups showed similar sign tracking (**Fig. 5F**; Time p<0.0001; no group differences) and CS+ accuracy (**Fig. 5G**; Time p<0.01; no group differences).

Thus, both 33-GCR and CDDO-EA enhanced goal tracking in females, with effects accumulating across treatments, while sign tracking remained unaffected in both sexes. This pattern of results — altered hippocampal function in males but altered reward processing in females — indicates that 33-GCR does not produce uniform effects across interconnected circuit nodes.

### CDDO-EA impairs reversal learning in males; 33-GCR impairs reversal learning in females

Following acquisition, mice underwent reversal learning in which CS+ and CS− locations were switched (**Fig. 6A**). Males were tested for 5 days; females for 6 days.

**Fig. 6.**
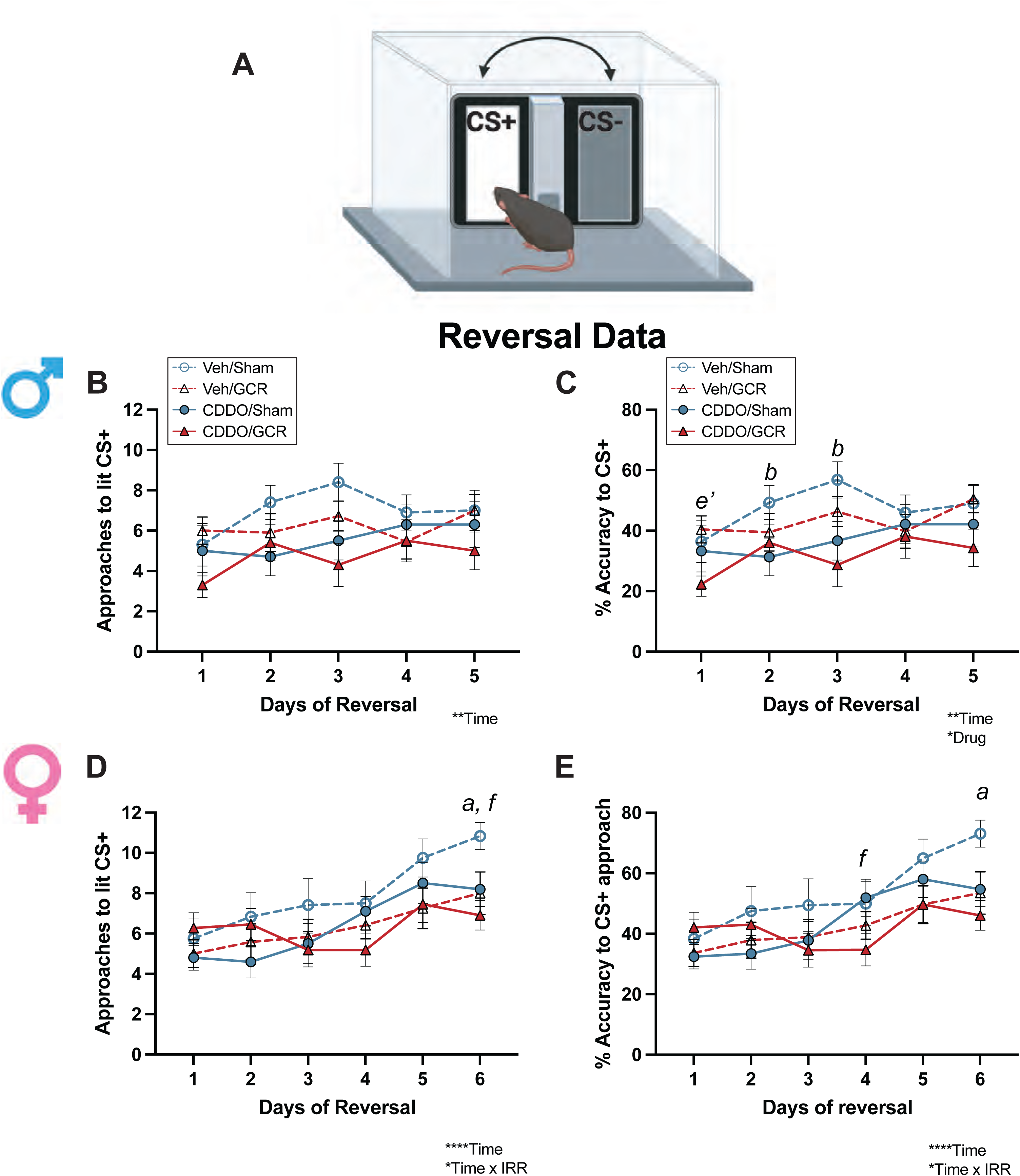
CDDO-EA impairs male reversal learning; 33-GCR impairs female reversal learning. **(A)** Schematic of the reversal learning task. Following 11 days of acquisition, the CS+ and CS− locations were switched. Males underwent 5 days of reversal; females underwent 6 days. **(B–C)** Male reversal learning: approaches to the new CS+ **(B)** and accuracy **(C)**. **(D–E)** Female reversal learning: approaches to the new CS+ **(D)** and accuracy **(E)**. Data are presented as mean ± SEM (n=10–12/group). Statistical analysis: 3-way RM ANOVA (IRR × Drug × Time). *p<0.05, **p<0.01, ***p<0.001, ****p<0.0001. Post-hoc: Veh/Sham vs Veh/33-GCR, ᵃ p<0.05; Veh/Sham vs CDDO-EA/Sham, ᵇ p<0.05; Veh/33-GCR vs CDDO-EA/33-GCR, ᵉ’ p<0.01; CDDO-EA/Sham vs CDDO-EA/33-GCR, ᶠ p<0.05. Complete subject numbers and detailed statistical analyses are provided in **S1 Table**.

In males, all groups showed similar numbers of approaches to the new CS+ location across reversal days (**Fig. 6B**, **S1 Table**; 3-way RM ANOVA, Time p<0.01; Drug, IRR, and interactions all p>0.05). However, CDDO-EA impaired reversal accuracy regardless of irradiation status (**Fig. 6C**; Time p<0.01, Drug p<0.05). On Day 1, CDDO-EA/33-GCR males showed 45% lower accuracy than Veh/33-GCR males (post-hoc p<0.01). On Days 2 and 3, CDDO-EA/Sham males showed 36-37% lower accuracy than Veh/Sham males (post-hoc p<0.05). Thus, CDDO-EA impaired behavioral flexibility in males independent of radiation exposure.

In females, 33-GCR impaired reversal learning regardless of CDDO-EA treatment. On Day 6, both irradiated groups made fewer approaches to the new CS+ than their respective sham controls: Veh/33-GCR females made 26% fewer approaches than Veh/Sham females, and CDDO-EA/33-GCR females made 16% fewer approaches than CDDO-EA/Sham females (**Fig. 6D**, **S1 Table**; 3-way RM ANOVA, Time p<0.0001, Time×IRR p<0.05; post-hoc p<0.05 for both comparisons). Similarly, 33-GCR reduced reversal accuracy: CDDO-EA/33-GCR females showed 33% lower accuracy than CDDO-EA/Sham females on Day 4 (post-hoc p<0.05), and Veh/33-GCR females showed 27% lower accuracy than Veh/Sham females on Day 6 (**Fig. 6E**; Time p<0.0001, Time×IRR p<0.05; post-hoc p<0.05).

The source of behavioral flexibility impairment thus differed between sexes: CDDO-EA impaired reversal learning in males regardless of radiation status, while 33-GCR impaired reversal learning in females regardless of CDDO-EA treatment. This is a direct countermeasure trade-off: CDDO-EA normalized radiation-enhanced pattern separation in males (**Fig. 2**) while independently impairing reversal learning in the same animals. This effect would be missed entirely by any study assessing only a single cognitive endpoint.

### Neither 33-GCR nor CDDO-EA affects anxiety-like behavior; combined treatment reduces locomotion in females

To determine whether the locomotor reductions observed across testing contexts (**Figs. 3, 4**) reflected anxiety, mice were tested on the elevated plus maze (EPM) at 6.5 months (males) and 6.75 months (females) post-IRR. Males showed no group differences in time spent in open arms, open arm entries, open-to-closed arm ratio, or total distance traveled (**Fig. 7A-D**, **S1 Table**; 2-way ANOVA, all p>0.05 or post-hoc p>0.05). In females, Drug×IRR interactions emerged for open arm time and entries (**Fig. 7E-F**; Drug×IRR p<0.05 and p<0.01, respectively). However, the open-to-closed arm ratio, a locomotion-independent measure of anxiety, did not differ among female groups (**Fig. 7G**; all post-hoc p>0.05), indicating no true anxiety-like phenotype.

**Fig. 7.**
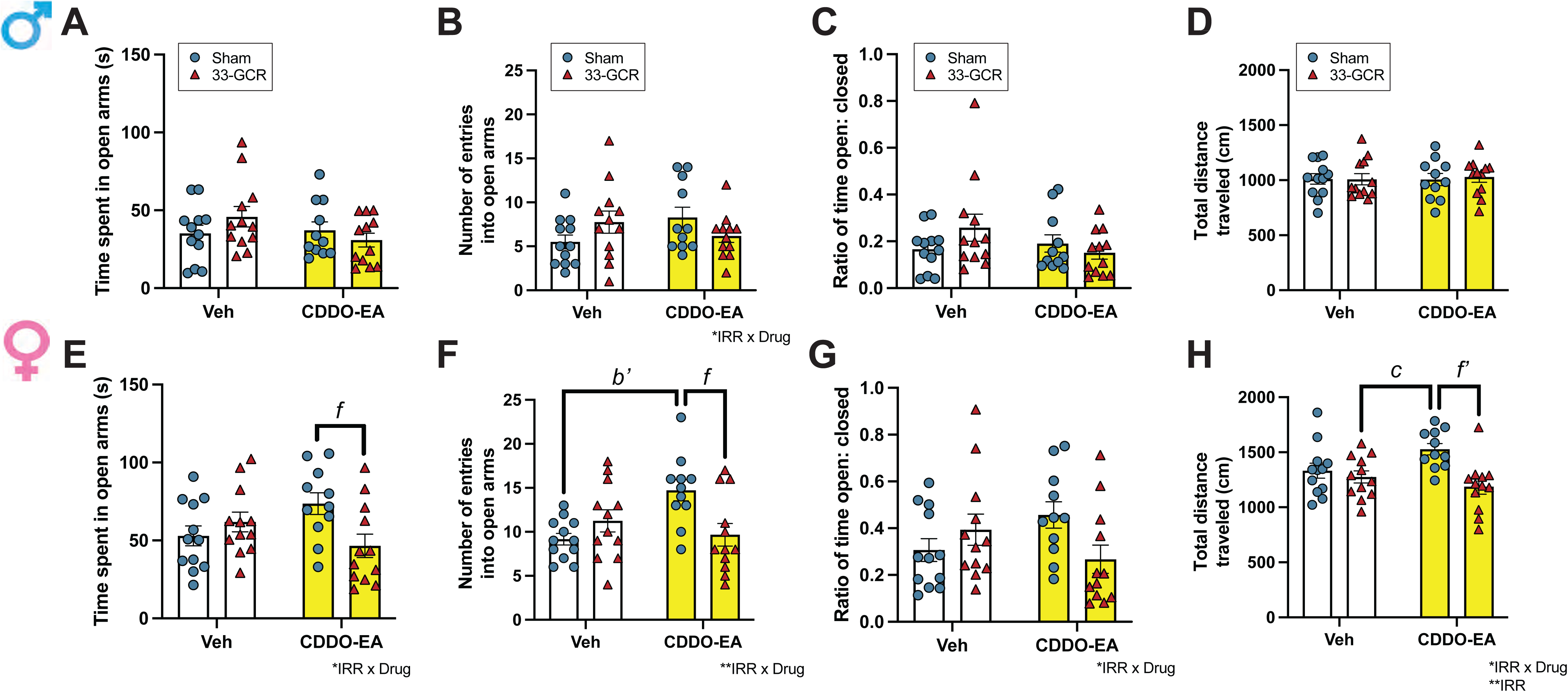
Neither 33-GCR nor CDDO-EA affects anxiety-like behavior; combined treatment reduces locomotion in females. **(A–H)** Mice were recorded during 5 minutes of free exploration under 200 lux in an elevated plus maze (EPM) with male mice at 6.5 months post-IRR **(A–D)** and female mice at 6.75 months post-IRR **(E–H)**. Anxiety-like behaviors were measured by time spent **(A, E)** and number of entries **(B, F)** in open arms, and the ratio of open to closed arm time **(C, G)**. Locomotor activity was measured by total movement distance (cm) in the entire arena **(D, H)**. Data are presented as mean ± SEM. Statistical analysis: 2-way ANOVA (IRR × Drug) in **A–H**. Main effect and/or interaction denoted by *p<0.05, **p<0.01; post-hoc: Veh/Sham vs CDDO-EA/Sham, ᵇ’ p<0.01; Veh/Sham vs CDDO-EA/33-GCR, ᶜ p<0.05; CDDO-EA/Sham vs CDDO-EA/33-GCR, ᶠ p<0.05, ᶠ’ p<0.01. Complete subject numbers and detailed statistical analyses are provided in **S1 Table**.

Rather, these interactions reflected locomotor differences: CDDO-EA/33-GCR females traveled 22% shorter distances than CDDO-EA/Sham females (**Fig. 7H**; IRR p<0.01, Drug×IRR p<0.05; post-hoc p<0.01), consistent with the home cage activity and SLR locomotion patterns described above. These data indicate that neither 33-GCR nor CDDO-EA induced anxiety-like behavior in either sex. The locomotor reductions observed throughout the study therefore reflect a separable physiological effect rather than anxiety-driven changes in exploration.

### Principal component analysis reveals circuit-specific treatment effects across cognitive domains

To examine how 33-GCR and CDDO-EA affected relationships among interconnected cognitive domains, we performed principal component analysis (PCA) on six behavioral measures: pattern separation at three difficulty levels (d-SLR, s-SLR, xs-SLR), goal tracking, sign tracking, and reversal learning (**Fig. 8**). The first three principal components explained 57.6% of total variance (PC1: 20.8%, PC2: 19.7%, PC3: 17.2%; **Fig 8A**).

**Fig. 8.**
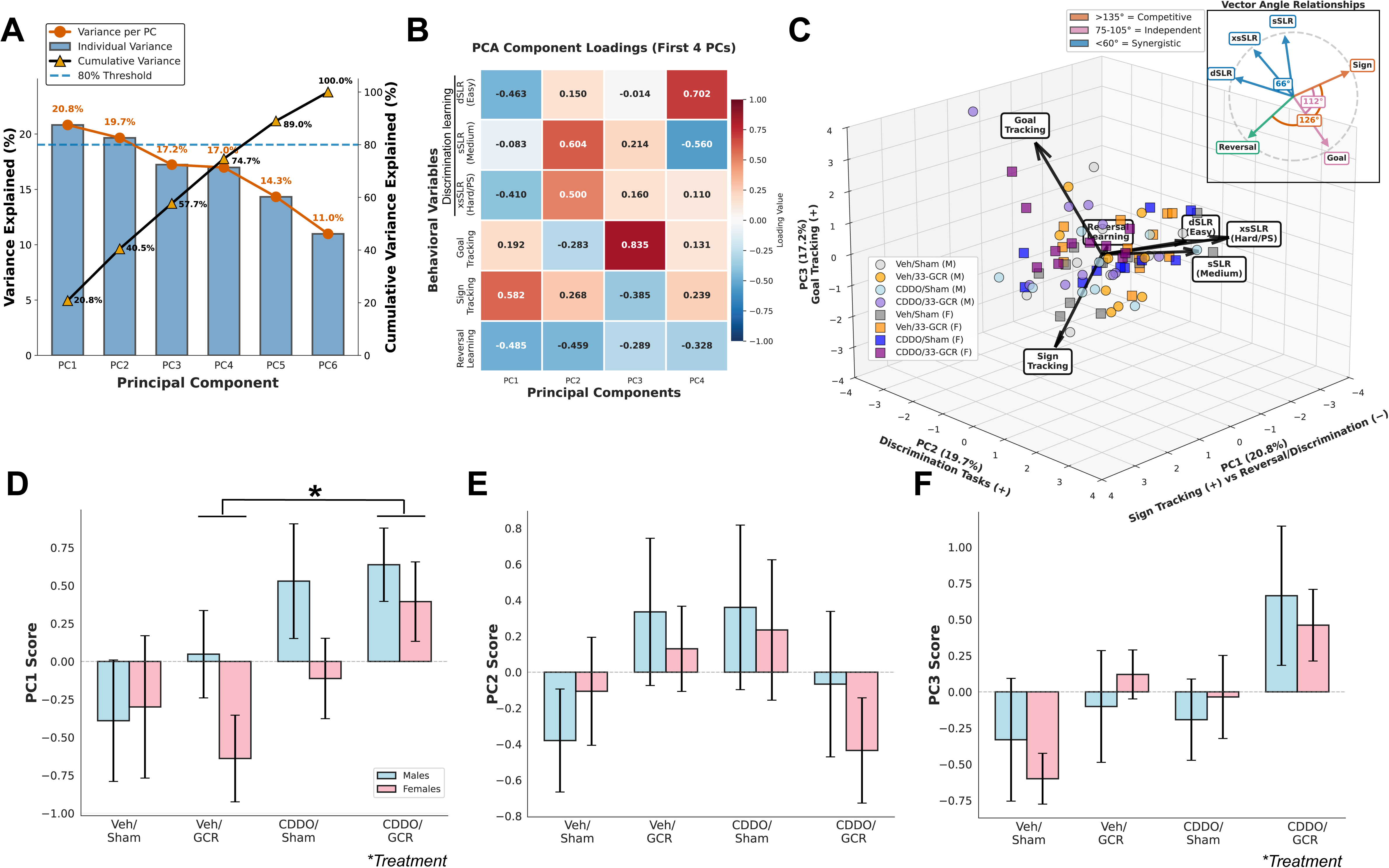
Principal component analysis reveals circuit-specific treatment effects across cognitive domains. **(A)** Scree plot showing variance explained by each PC. Blue bars: individual variance; orange circles/line: cumulative variance; dashed line: 80% threshold. **(B)** Component loading heatmap for the first four PCs across six behavioral variables: d2 ratio (d-SLR, s-SLR, xs-SLR), sign-tracking (CS+ vs CS− approach difference, Days 4–7), goal-tracking (reward chamber approaches, Days 4–7), and reversal learning (CS+ approaches, Days 1–4). Red: positive loading; blue: negative loading. **(C)** Three-dimensional biplot showing individual animals across PC1 × PC2 × PC3 (57.6% total variance). Black arrows: loading vectors. Points color-coded by group (gray: Veh/Sham; orange: Veh/33-GCR; light blue: CDDO-EA/Sham; purple: CDDO-EA/33-GCR) and shape (circles: males; squares: females). Inset: vector angle relationships indicating competitive (>135°, orange), independent (∼90°, pink), and synergistic (<60°, blue) domain interactions. **(D–F)** Treatment effects on PC scores shown separately for males (blue) and females (pink). Statistical analysis: 2-way ANOVA (Treatment × Sex). *p<0.05. Post-hoc: Veh/33-GCR vs CDDO-EA/33-GCR, * p<0.05. Complete subject numbers and detailed statistical analyses are provided in **S1 Table**.

PC1 captured an axis contrasting stimulus-driven responding with executive control. Sign tracking loaded positively (+0.582) while reversal learning (−0.485) and discrimination tasks (d-SLR: −0.463; s-SLR: −0.410) loaded negatively (**Fig. 8B**). Vector angles confirmed competitive interactions (>135°) between sign tracking and reversal/discrimination tasks (**Fig. 8C**, inset, orange arcs). CDDO-EA/33-GCR animals showed higher PC1 scores than Veh/33-GCR animals (**Fig. 8D**; 2-way ANOVA, Treatment p<0.05; post-hoc p<0.05), indicating a shift toward stimulus-driven responding in the combined treatment group.

PC2 captured an axis contrasting hippocampal-dependent discrimination with reversal learning. The discrimination tasks loaded positively (xs-SLR: +0.835; s-SLR: +0.604) while reversal learning loaded negatively (−0.459). Vector angles between discrimination tasks were <60° (**Fig. 8C**, inset, blue arcs), indicating coordinated, rather than competing, operations. PC2 scores did not differ across treatment groups or sex (**Fig. 8E**; 2-way ANOVA, all p>0.05), indicating that hippocampal discrimination capacity was preserved despite radiation and drug treatment.

PC3 captured goal-directed behavior, with goal tracking loading strongly positive (+0.702) and sign tracking (−0.385) and reversal (−0.328) loading negatively. Vector angles showed goal tracking oriented ∼90° from sign tracking (**Fig. 8C**, inset, purple arcs), indicating functional independence. PC3 scores differed across treatment groups (**Fig. 8F**; 2-way ANOVA, Treatment p<0.05), consistent with the selective goal-tracking changes observed in females.

These data demonstrate that 33-GCR and CDDO-EA selectively altered specific circuit relationships, particularly the balance between stimulus-driven and goal-directed behavior (PC1, PC3), while still preserving hippocampal discrimination capacity (PC2). Preserved PC2 scores indicate that the hippocampus itself was not globally impaired. Instead, the behavioral effects observed in males reflect altered circuit communication involving NAc and PFC nodes rather than hippocampal damage *per se*. This selective vulnerability with preserved function is consistent with the domain-specific patterns observed in individual behavioral measures (**Figs. 2-6**).

### 33-GCR produces persistent alterations in dentate gyrus activity during memory encoding

To assess whether the domain-specific behavioral effects reflected stable changes in hippocampal circuit function, we performed fiber photometry in a subset of male Veh/Sham and Veh/33-GCR mice at 7.25 months post-IRR (**Fig. 9A-B**). Ca²⁺ transients were recorded from DG glutamatergic neurons during SLR testing. Behaviorally, in this small surgical cohort (n=3–4/group), both groups showed positive d2 ratios under d-SLR. Under xs-SLR, the directional pattern mirrored the main cohort: Veh/Sham mice showed a negative d2 ratio (−0.15±0.17) while Veh/33-GCR mice showed a positive d2 ratio (+0.21±0.06; **Fig. 9C**). This did not reach statistical significance, consistent with the reduced power of this subset (all p>0.05).

**Fig. 9.**
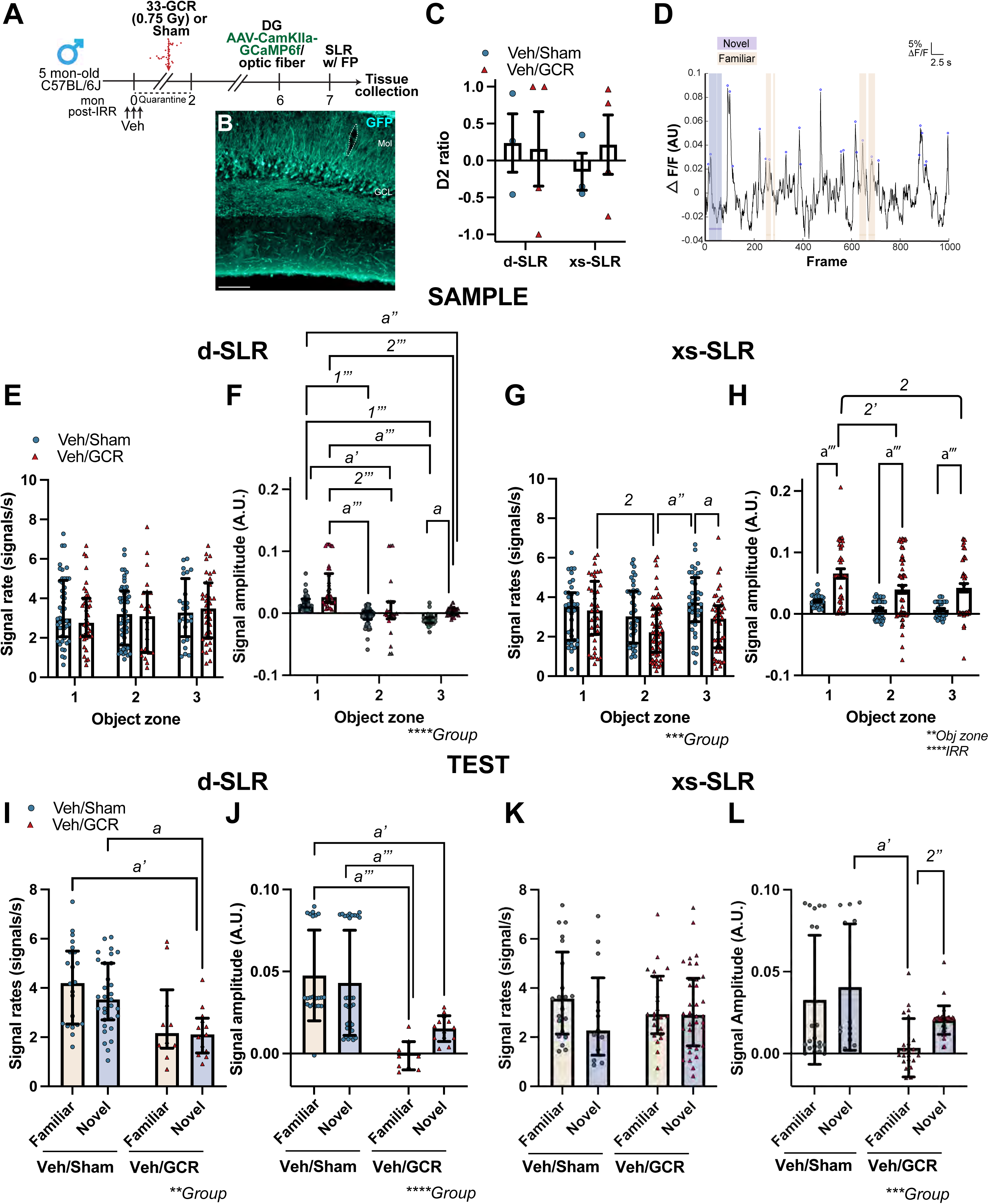
33-GCR produces persistent alterations in dentate gyrus activity during memory encoding. **(A)** Experimental timeline. A subset of male Veh/Sham and Veh/33-GCR mice (n=3–4/group) underwent AAV-CaMKIIα-GCaMP6f viral infusion in the DG hilus and optic fiber implantation in the molecular layer at 6 months post-IRR. Mice performed SLR with simultaneous fiber photometry recording at 7.25 months post-IRR. **(B)** GFP expression in the DG granule cell layer; dotted line indicates fiber track. Scale bar: 100 μm. **(C)** Discrimination index (d2 ratio) in d-SLR and xs-SLR. **(D)** Representative Ca²⁺ signal (ΔF/F) during the Test phase. Purple: novel location; beige: familiar location. Scale bar: 5% ΔF/F, 2.5 s. **(E–H)** Sample phase Ca²⁺ signals: transient rate **(E)** and amplitude **(F)** in d-SLR; transient rate (G) and amplitude **(H)** in xs-SLR. **(I–L)** Test phase Ca²⁺ signals: transient rate **(I)** and amplitude **(J)** in d-SLR; transient rate **(K)** and amplitude **(L)** in xs-SLR. Data are presented as mean ± SEM **(C, H)** or median ± IQR **(E–G, I–L)**. Each symbol indicates an animal **(C)** or detected event **(E–L)**. Statistical analysis: 2-way RM ANOVA **(C, H)**; Kruskal-Wallis with Dunn’s post-hoc **(E–G, I–L)**. **p<0.01, ***p<0.001, ****p<0.0001. Post-hoc: Veh/Sham vs Veh/33-GCR, ᵃ p<0.05, ᵃ’ p<0.01, ᵃ’’ p<0.001, ᵃ’’’ p<0.0001. Complete subject numbers and detailed statistical analyses are provided in **S1 Table**.

During the Sample phase (encoding), Veh/33-GCR mice showed enhanced DG activity compared to Veh/Sham mice. Under d-SLR, Ca²⁺ transient rates were similar between groups (p>0.05; **Fig. 9E**), but Veh/33-GCR mice showed 127% higher signal amplitudes at Object zone 3 than Veh/Sham mice (Kruskal-Wallis p<0.0001; post-hoc p<0.0001 at all zones; **Fig. 9F**). Under xs-SLR, Veh/33-GCR mice showed 29% lower transient rates at Object 3 than Veh/Sham mice (Kruskal-Wallis p<0.001; post-hoc p<0.05; **Fig. 9G**) but 210-516% higher amplitudes across all object zones (Object p<0.01, IRR p<0.0001; post-hoc p<0.0001 at all zones; **Fig. 9H**).

During the Test phase (retrieval), Veh/33-GCR mice showed 40% lower transient rates at the Novel location than Veh/Sham mice under d-SLR (Kruskal-Wallis p<0.01; post-hoc p<0.01; **Fig. 9I**) and 103% lower amplitudes at the Familiar location (p<0.0001; post-hoc p<0.001; **Fig. 9J**)."Under xs-SLR, transient rates did not differ across groups (p>0.05; **Fig. 9K**). Signal amplitudes showed a significant group effect (Kruskal-Wallis p<0.001; **Fig. 9L**): Veh/33-GCR mice showed higher amplitude at the Novel versus Familiar location (p<0.001), whereas Veh/Sham mice showed no location-specific difference.

Given the small cohort size (n=3–4/group), these findings should be interpreted with caution and treated as preliminary; the patterns reported here are intended to motivate future work with adequately powered samples rather than to support mechanistic conclusions. These findings are consistent with 33-GCR producing persistent alterations in DG circuit activity detectable 7 months post-IRR, well beyond the behavioral testing window. The enhanced encoding amplitudes and location-selective retrieval patterns suggest that hippocampal circuit dynamics may be persistently shifted rather than transiently perturbed.

### Combined CDDO-EA and 33-GCR reduces dentate gyrus progenitor cells in females despite intact pattern separation

To assess whether treatments affected cellular indices relevant to hippocampal function, we quantified doublecortin-immunoreactive (DCX+) cells in the dentate gyrus at 7.25 months post-IRR (**Fig. 10A**).

**Fig. 10.**
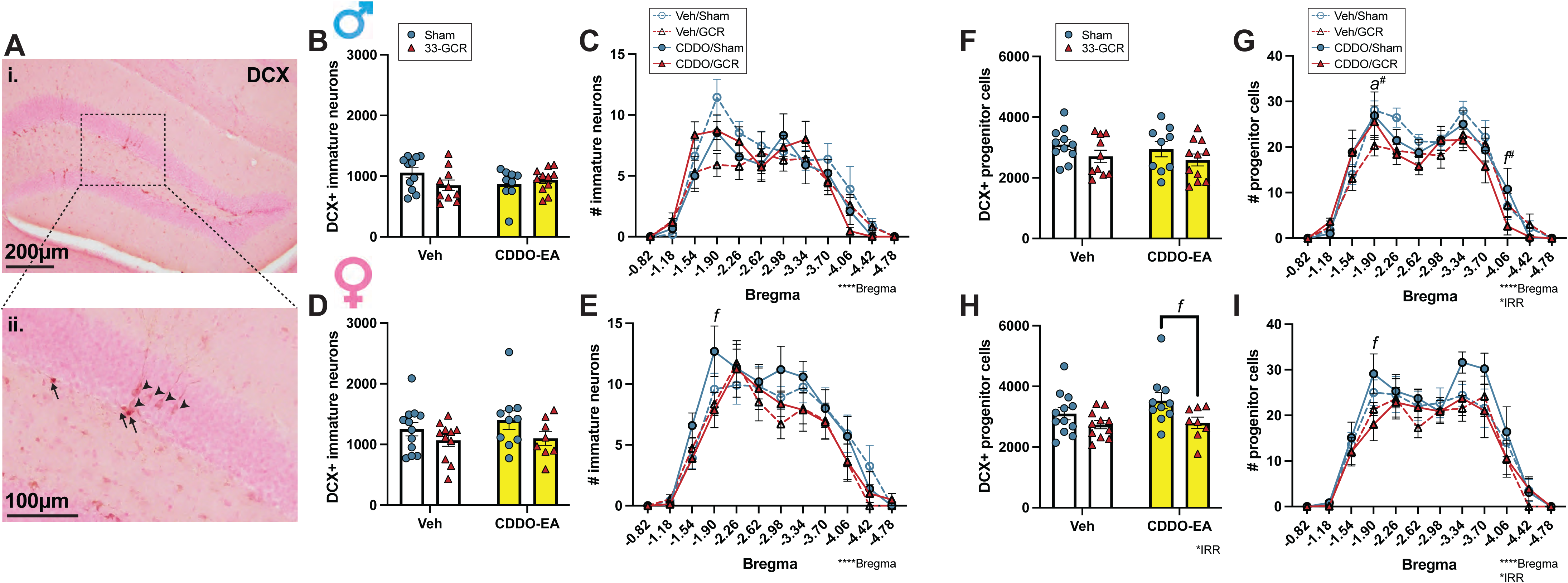
Combined CDDO-EA and 33-GCR reduces DCX+ progenitor cells in females despite intact pattern separation. **(A)** Representative photomicrographs of DCX+ cells in the DG at 100× (i) and higher magnification (ii). Immature neurons were identified by oval cell bodies with extensive dendritic processes containing at least one dendritic node (arrowheads). Progenitor cells were identified by irregular cell bodies with short processes lacking dendritic nodes (arrows). **(B–I)** Quantification of DCX+ cells using unbiased stereology in males **(B, C, F, G)** and females **(D, E, H, I)**. Total DCX+ immature neurons **(B, D)** and distribution across the rostro-caudal axis **(C, E)**. Total DCX+ progenitor cells **(F, H)** and distribution across the rostro-caudal axis **(G, I)**. Data are presented as mean ± SEM (n=8–12/group). Statistical analysis: 2-way ANOVA (IRR × Drug) in **B, D, F, H**; 3-way RM ANOVA (Bregma × IRR × Drug) in **C, E, G, I**. *p<0.05, ****p<0.0001. Post-hoc: CDDO-EA/Sham vs CDDO-EA/33-GCR, ᶠ p<0.05. Scale bars: 200 μm (i), 100 μm (ii). Complete subject numbers and detailed statistical analyses are provided in **S1 Table**.

In males, all groups showed similar numbers of DCX+ immature neurons (**Fig. 10B**, **S1 Table**; 2-way ANOVA, all p>0.05), with comparable distribution across the rostro-caudal axis (**Fig. 10C**; 3-way RM ANOVA, Bregma p<0.0001; no group differences). DCX+ progenitor cell numbers were also similar across groups (**Fig. 10F**; all p>0.05), with no differences along the rostro-caudal axis (**Fig. 10G**; Bregma p<0.0001; all post-hoc p>0.05).

In females, DCX+ immature neuron numbers were similar across all groups (**Fig. 10D**; 2-way ANOVA, all p>0.05), with comparable rostro-caudal distribution (**Fig. 10E**; 3-way RM ANOVA, Bregma p<0.0001). However, 33-GCR reduced DCX+ progenitor cells in CDDO-EA-treated females. CDDO-EA/33-GCR females had 21% fewer total DCX+ progenitor cells than CDDO-EA/Sham females (**Fig. 10H**; 2-way ANOVA, IRR p<0.05; post-hoc p<0.05). This reduction was particularly evident in the dorsal DG: at Bregma −1.90, CDDO-EA/33-GCR females had significantly fewer progenitors than CDDO-EA/Sham females (**Fig. 10I**; 3-way RM ANOVA, Bregma p<0.0001, IRR p<0.05; post-hoc p<0.05). Veh/33-GCR females did not differ from Veh/Sham females (all p>0.05), indicating that this progenitor reduction required the combination of CDDO-EA and 33-GCR. This suggests that Nrf2 activation may alter the cellular context in which radiation affects progenitor populations.

These data demonstrate that combined CDDO-EA and 33-GCR exposure reduced DCX+ progenitor cells in females, particularly in the dorsal hippocampus, despite intact pattern separation performance in these animals (**Fig. 3**). Pattern separation was assessed at 3.75 months post-IRR while DCX+ cells were quantified at 7.25 months post-IRR; these measurements are therefore temporally decoupled and cannot be interpreted as a direct cellular-behavioral dissociation.

## Discussion

Prior GCR studies have assessed single cognitive endpoints in single sexes, leaving open whether radiation effects are global or circuit-selective and whether countermeasures protect uniformly or trade off across domains. The present data resolve both questions, and the answers are more complex than either framing anticipated. Here we show that 33-GCR produced domain-specific rather than global cognitive effects, with distinct vulnerability patterns in males and females. CDDO-EA acted as a double-edged sword: normalizing some radiation effects while impairing others, a trade-off invisible to single-endpoint assessment.

In males, 33-GCR enhanced pattern separation specifically under very high memory load, consistent with prior single-ion studies showing improved pattern separation following ⁵⁶Fe or ²⁸Si exposure in males [93,94]. The convergence across radiation types and testing paradigms suggests this reflects a genuine radiation effect rather than a paradigm artifact. CDDO-EA normalized this enhancement, to our knowledge the first demonstration that this compound can reverse radiation-induced enhancements rather than solely prevent impairments. Enhanced pattern separation does not necessarily indicate improved cognitive function: in clinical populations, overly distinct memory representations can interfere with generalization and flexible cognition [95].

In females, pattern separation did not differ among groups at any memory load, contrasting with prior work reporting impairments at earlier timepoints using a different paradigm [58]. That task required spatial memory retention over ≥24 hours, whereas the SLR task tests discrimination within ∼35 minutes, a fundamental difference in memory demand. Furthermore, females in the prior work showed hippocampal-dependent deficits during training before pattern separation testing [58], raising the possibility that pre-existing impairments confounded performance. Consistent with the present findings, a prior 33-GCR study in females using touchscreen-based tasks also found no radiation effect on pattern separation [91].

Fiber photometry revealed enhanced DG signal amplitudes during encoding in irradiated males, particularly under very high memory load, with location-selective retrieval patterns at 7 months post-IRR. To our knowledge, this is the first use of in vivo Ca²⁺ imaging to assess hippocampal activity during cognitive testing in the context of space radiation. The stage-specific nature of these alterations, enhanced encoding but shifted retrieval, points to changes in circuit dynamics rather than uniform suppression or excitation [96,97]. Increased encoding amplitude may strengthen memory formation and contribute to more distinct representations [33,34,98,99], though causal relationships cannot be established from these data. These alterations were detectable 7 months post-IRR, suggesting stable rather than transient changes in circuit function. The small cohort size (n=3-4/group) limits interpretation; these findings are best treated as preliminary.

DCX+ progenitor cells were reduced in CDDO-EA/33-GCR females at 7.25 months post-IRR, but not in Veh/33-GCR females, indicating this reduction required combined treatment. This contrasts with our prior work showing that 33-GCR alone reduces DCX+ immature neurons but not progenitors at 14.25 months post-IRR [91], suggesting CDDO-EA may alter the temporal trajectory of radiation effects on neurogenesis-related populations: combined treatment affects earlier-stage progenitor cells at intermediate timepoints, whereas radiation alone affects later-stage immature neurons at extended timepoints. Since pattern separation was assessed at 3.75 months post-IRR and DCX+ cells at 7.25 months post-IRR, these measurements are temporally decoupled and cannot be interpreted as a direct cellular-behavioral dissociation; rather, they reveal a late-emerging cellular consequence of combined treatment not captured by behavioral assessment alone.

Female mice often show resilience to space radiation-induced cognitive deficits relative to males [22,65,75,76,102–106], and the present findings are consistent with this pattern. In males, 33-GCR altered hippocampal-dependent cognition without affecting body weight or home cage activity, suggesting effects concentrated in cognitive circuits rather than distributed across metabolic and motor systems [93,100,101]. The mechanisms underlying female resilience are not established; proposed factors include differences in neuroimmune responses, antioxidant capacity, and gonadal hormone signaling [22,107–109], though which, if any, contribute here cannot be determined from the present data. Preserved cognition in females may nonetheless come at a metabolic cost: persistent weight gain in Veh/33-GCR females suggests radiation-induced disruption of energy homeostasis [55,110,111], and reduced locomotor activity may reflect metabolic changes or altered motivation rather than cognitive impairment [91]. These distinct patterns between males and females underscore why single-sex studies - still common in radiation neuroscience [22,65,75,102] and in neuroscience more broadly [32] - can produce incomplete or misleading conclusions about stressor effects on cognition.

Locomotor reductions were observed across multiple contexts in both sexes but followed sex-specific patterns: in males, CDDO-EA alone reduced home cage activity while radiation had no additional effect, whereas in females both 33-GCR and CDDO-EA independently reduced activity across home cage, SLR, and EPM contexts. These reductions did not confound cognitive outcomes: pattern separation was enhanced in irradiated males despite no locomotor changes, and intact in females despite pronounced locomotor reductions. The EPM confirmed these reductions do not reflect anxiety, supporting the interpretation that locomotor and cognitive effects represent separable consequences of treatment.

The autoshaping paradigm dissociates sign-tracking (dorsolateral striatum/amygdala-dependent), goal-tracking (NAc/PFC-dependent), and reversal learning (OFC-dependent) [43,112–117]. The selective enhancement of goal-tracking in females, with no autoshaping effects in males, points to regional rather than uniform circuit vulnerability. Goal-tracking changes without corresponding hippocampal alterations may reflect the NAc’s integration of multiple input streams beyond the hippocampus, including direct cortical and amygdala inputs [118–120].

CDDO-EA enhanced goal-tracking in females but had no reward-related effects in males, indicating sex-specific modulation of NAc-PFC circuitry by Nrf2 activation [121].This sex specificity extended to the combined treatment: CDDO-EA and 33-GCR produced additive goal-tracking enhancement in females, with no parallel effect in males. This raises the possibility that GCR and Nrf2 activation converge on shared circuitry in a sex-dependent manner, consistent with evidence that opponent striatal monoamine signaling modulates reinforcement behavior [120]. The specificity of these effects to goal-tracking rather than sign-tracking suggests CDDO-EA preferentially modulates NAc-PFC circuits [43,112–117], possibly through sex-dependent differences in dopaminergic signaling or Nrf2 expression [83].

In males, CDDO-EA impaired reversal accuracy without affecting initial acquisition, pointing to selective disruption of behavioral updating — the ability to revise a learned response when contingencies change — rather than reward learning itself. In females the picture was different; the drug was not the culprit, radiation was, and the impairment likely reflects the learning context established during acquisition. Enhanced goal-tracking in irradiated females may have created strong reward-location associations that made updating cue-reward contingencies particularly difficult, given that sign-tracking and goal-tracking behaviors resist contingency changes [122–125]. Reversal impairment looked the same on the surface in both sexes but arose from distinct circuit-level processes: behavioral updating in males, acquisition strategy in females.

The autoshaping reversal impairment in females contrasts with our previous finding that CDDO-EA/33-GCR females showed enhanced cognitive flexibility in touchscreen-based spatial discrimination reversals [91]. This dissociation is reconcilable: the touchscreen task requires spatial discrimination without competing goal-tracking strategies, so reversal does not require overcoming a strongly established reward-location habit. Irradiated females thus retain capacity for flexible behavioral updating; the impairment is context-dependent, not a signature of broad OFC dysfunction.

PCA quantified what the individual behavioral results suggested qualitatively. The PC2 null result is particularly informative: pattern separation tasks and reversal learning maintained their relationship across all treatment groups, indicating the hippocampus itself was not globally disrupted. The behavioral effects seen in males therefore reflect altered signaling at NAc and PFC nodes rather than hippocampal damage per se. PC1 and PC3, by contrast, shifted with treatment, capturing the rebalancing of stimulus-driven versus goal-directed behavior. Some circuit relationships were disrupted; others held. That selective pattern, rather than uniform impairment, is the defining feature of circuit-selective vulnerability.

Domain-specific drug-induced cognitive effects are well recognized in clinical pharmacology [130]; preclinical countermeasure evaluation has rarely followed suit, largely because single cognitive endpoints remain the norm [127–129]. Region-specific neural effects of environmental stressors are equally well established [1,18,126], but the corollary - that countermeasures may show domain-dependent trade-offs - has not been systematically tested. The practical consequences are stark: a study assessing only pattern separation would conclude CDDO-EA is protective; a study assessing only reversal learning would conclude it is harmful. Neither conclusion captures the reality that CDDO-EA’s effects are circuit-dependent, sex-dependent, and only fully visible through multi-domain assessment.

Three limitations warrant mention. Fiber photometry and DCX+ quantification were conducted at 7–7.25 months post-IRR while behavioral testing occurred at 3.25–4.5 months; concurrent measurements at matched timepoints are needed to establish whether cellular changes relate directly to behavioral performance. Fiber photometry was conducted only in males; extending these recordings to females would clarify whether distinct circuit activity patterns underlie preserved pattern separation in that sex. The separate statistical analyses of males and females, while appropriate given divergent response profiles, preclude direct statistical comparison of sex differences, a limitation shared broadly in the field [32].

Male and female mice showed distinct, circuit-selective vulnerability patterns that would have been invisible to single-task or single-sex designs. The core finding is not simply that radiation affects cognition, but that its effects are circuit-specific and sex-dependent. A candidate countermeasure can simultaneously protect one function while impairing another, a trade-off invisible to single-endpoint assessment. As space agencies plan missions beyond Earth orbit, these findings have immediate relevance for astronaut risk assessment. Any stressor or neuroprotective intervention evaluated with limited behavioral endpoints risks missing exactly these trade-offs; the present framework provides a template for doing better.

## Materials and methods

### Animals

Male (M, n=124) and female (F, n=48) C57BL/6J mice (*Mus musculus* C57BL/6J, 4.5-5 months old; RRID:IMSR_JAX:000664; Jackson Laboratory, Bar Harbor, ME) were shipped to Brookhaven Laboratory Animal Facility (BLAF) at Brookhaven National Laboratory (BNL, Upton, NY). Males arrived in variable group sizes (2 cages of 5 mice, 23 cages of 4 mice, 1 cage of 3 mice, 8 cages of 2 mice, and 3 single-housed mice), while females arrived in uniform groups (12 cages of 4 mice). Upon arrival, mice were maintained with their original cagemates throughout the study. After three days of acclimation, mice received ear punch followed by first weight measurement. Two days later, mice were transported to the NASA Space Radiation Laboratory (NSRL) within BLAF for treatment with either 33-GCR IRR or Sham IRR and returned to BLAF the following day. At both facilities, mice were housed 4/cage in HEPA-filtered, closed airflow vivarium systems under a 12:12 h light/dark cycle (06:00 light on) at 22°C, 30-70% humidity with standard rodent chow (5015; Lab Diet, cat# 0001328) and water ad libitum. Two days post-IRR, mice were transported by ground to Children’s Hospital of Philadelphia (CHOP) and held in quarantine for 6 weeks with ad libitum access to medicated chow (13 PPM ivermectin and 150 PPM fenbendazole; Test Diet, custom cat# 1813527[5SKU]). Behavioral testing began at 9 weeks post-IRR when mice were released from quarantine and returned to standard chow (5015). At CHOP, mice were housed in HEPA-filtered, closed airflow vivarium systems (Enviro-Gard™ A; Lab Products Inc.) under a 12:12 h light/dark cycle (06:15 light on) at 20-23°C, 30-40% humidity. Each cage received a nestlet square at cage changes; no other enrichment was provided. All procedures were approved by IACUCs at BNL and CHOP in accordance with AAALAC and NIH guidelines (CHOP: AAALAC #000427, PHS D16-00280 [OLAW A3442-01]; BNL: AAALAC #000048, PHS D16-00067 [OLAW A3106-01]). Our reporting adheres to ARRIVE 2.0 guidelines [131].

### Drug administration

Cages were sequentially preassigned to treatment groups in rotating order (Veh/Sham, Veh/33-GCR, CDDO-EA/Sham, CDDO-EA/33-GCR), balanced for mean cage body weight and cage size (number of mice per cage). Drug treatment groups consisted of mice receiving either CDDO-EA (2-cyano-3,12-dioxooleana-1,9-dien-28-oic acid ethylamide, 4 mg/kg IP; MedChemExpress, cat# HY-12213; n=83, M:59, F:24) or vehicle control (matching volume IP; n=89, M:65, F:24).

The vehicle solution was prepared according to the manufacturer’s instructions and consisted of 5% DMSO (Sigma-Aldrich, cat# D2650) and 20% Sulfobutylether-β-Cyclodextrin (SBE; MedChemExpress, cat# HY-17031) in 0.9% saline solution (Grainger, cat# 3PWK4). Both CDDO-EA and vehicle were administered intraperitoneally once daily between 8:00-10:00 AM for three consecutive days. IRR was performed on Day 2 of the treatment regimen (one day after the first injection, concurrent with the second injection, and one day before the final injection).

### Irradiation (IRR)

On the second day of drug treatment (Day 0 of IRR), mice underwent IRR during the BNL 22A campaign as previously described [91]. Mice were placed with cagemates in well-ventilated polycarbonate containers (10 × 10 × 4.5 cm), with 2 mice per container when possible or individually housed if no cagemate was available. Irradiated mice were exposed to 75 cGy of NASA’s whole-body 33-beam GCR simulation delivered over a 60 × 60 cm field for approximately 1.25 hours beginning at 11:45 AM. Beam uniformity and dosimetry were monitored by NSRL staff. Sham-irradiated mice were placed in containers with cagemates (or individually) for the same duration but were not exposed to the beam. This resulted in four experimental groups: Veh/Sham (M:32, F:12), Veh/33-GCR (M:33, F:12), CDDO-EA/Sham (M:30, F:12), and CDDO-EA/33-GCR (M:29, F:12).

### Surgery

C57BL/6J male mice (3-4 mice/group) underwent viral infusion and fiber implantation at 6 months post-IRR. Before surgery, each mouse was weighed and the head was shaved using an electric clipper (Wahl, cat# 41590-0438). Anesthesia was induced in an induction chamber with 4-5% isoflurane (Piramal Pharma Limited, cat# 66794-013-25) in 100% oxygen and maintained at 1-3% isoflurane during surgery. Ophthalmic lubricant (Dechra, cat# B00HGMZ7RQ) was applied to both eyes, and buprenorphine ER (1 mg/kg, s.c.) was administered before incision. Following aseptic preparation of the surgical site with betadine (Avrio Health, cat# 67618-155-16) and 70% isopropyl alcohol, a midline incision was made. The skull was scored with a scalpel blade and treated with 30% hydrogen peroxide solution (Sigma Aldrich, cat# MKCJ1024). A burr hole was drilled above the target site using a surgical electric drill with a ⅛ inch engraving bit (Dremel, cat# 7350).

AAV9-CaMKII-GCaMP6f (Penn Vector Core, 100834) was unilaterally infused into the hilus of the dorsal dentate gyrus (DG) hilus (A/P -2.0 mm, M/L -1.4 mm, D/V -2.2 mm from bregma) using a 33-gauge Hamilton syringe (Hamilton, cat# 2141205) at 0.1 µl/min [132]. An optic fiber (Thorlabs, #CFML22L05, Ø1.25 mm SS ferrule, Ø200 µm core, 0.22 NA, L=2 mm) was implanted in the molecular layer of the DG middle/outer molecular layer (A/P -2.0 mm, M/L -1.4 mm, D/V -1.8 mm from bregma). The fiber was secured to the skull with light-cured resin (Ivoclar Vivadent AG, cat# 595979US) followed by light-cured adhesive (Pearson, cat# 595979). Following surgery, the incision was sutured, triple-antibiotic ointment was applied topically, and meloxicam (5 mg/kg, s.c.; Norbrook, cat# 5552904010) was administered. Mice were monitored daily for 48 hours post-operatively.

### Body weight monitoring and health observations

Mouse body weight was measured monthly from arrival at BNL (4.5 months of age) through tissue collection (12.25 months of age). Health status and cage conditions were monitored during weighing sessions and biweekly cage changes by the Children’s Hospital of Philadelphia (CHOP) Department of Veterinary Resources, with documentation of fighting, illness, or loss of cagemates. When aggressive behavior was identified, aggressor mice were isolated into separate cages to prevent injury. Seven male cages required splitting due to aggression (3 Veh/Sham, 3 CDDO-EA/Sham, 1 Veh/33-GCR). All mice from these disrupted cages were excluded from behavioral testing to maintain consistent social housing conditions across experimental groups.

### Behavioral testing

#### Overview of behavioral testing

Behavioral testing groups were selected in rotating order across treatment groups (Veh/Sham, Veh/33-GCR, CDDO-EA/Sham, CDDO-EA/33-GCR) and balanced for mean cage body weight. Most cages contained 4 mice per cage when behavioral testing began, except one female cage with 3 mice. Two deaths occurred: female #28 (CDDO-EA/Sham) died at 1 month post-IRR before behavioral testing began, and male #9 (CDDO-EA/Sham) died at 2.5 months post-IRR during locomotor testing. Both deaths reduced their respective cages to 3 mice. Despite these losses, both affected cages were retained for behavioral testing due to limited female availability and to avoid excluding the male cagemates who had already begun testing. Both deceased mice were excluded from analyses. This resulted in 47 mice per sex (total n=94) for behavioral testing. Following release from quarantine at 2 months post-IRR, behavioral testing began at 2.25 months post-IRR with locomotor activity recording (**Fig 1A**). Gentle handling (2 min/day) began at 2.5 months post-IRR and continued for 5 days. Male mice underwent arena habituation for 5 days beginning at 3 months post-IRR to acclimate to the Spontaneous Location Recognition (SLR) arena, followed by SLR testing at 3.25 months post-IRR. Males then completed the Autoshaping task (Pavlovian learning) at 3.5 months post-IRR and anxiety behavioral testing at 6.5 months post-IRR. Female mice underwent arena habituation for one week beginning at 3.5 months post-IRR, followed by SLR testing at 3.75 months post-IRR. Females then completed the Autoshaping task at 4.5 months post-IRR and anxiety behavioral testing at 6.75 months post-IRR.

#### Home cage activity monitoring

Each mouse was individually placed in a clean mouse conventional cage containing fresh bedding to record 18 hours of locomotion activity from 4pm to 10am. This cage was positioned between 4×8 photocells, with identical lighting parameters to home housing room dim/red lighting during the light cycle and red lighting during the dark cycle. Their movement across the XY plane was monitored by a computer-controlled photobeam activity system (San Diego Instruments), which recorded photocell beam breaks in 15-minute intervals over a period of 18 hours [133]. One male mouse from (mouse ID #10, group: CDDO/Sham) was excluded due to equipment failure.

#### Spontaneous location recognition (SLR)

The spontaneous location recognition (SLR) task was performed following established protocols [96]. *<u>Arena</u> <u>Setup and Objects.</u>* The testing arena consisted of a circular open field with 20 marked segments radiating from the center, etched by laser cutting at 18° intervals on the base and covered with corncob bedding during testing. Objects consisted of 50 ml conical centrifuge tubes containing three blue latex gloves, secured to the arena base with screws and nuts. Each arena was surrounded by three-sided black cardboard barriers displaying three distinct visual cues, with two additional cues mounted on the wall. These spatial cues remained consistent throughout testing to provide reliable landmark references. *<u>Habituation</u>*. Prior to SLR testing, mice underwent gentle handling (2 min/day) for 5 days to reduce stress, followed by arena habituation beginning at 3.0 months post-IRR for males and 3.5 months post-IRR for females. Subjects were placed in the arena with spatial cues for 10 minutes daily over five consecutive days. *<u>Testing Procedures</u>*. On test days, mice were transported to the testing room and acclimated in their home cages for 30 minutes. Testing time was kept consistent across subjects. Between subjects, one scoop of bedding was removed and replaced with clean bedding, and arena floors and walls were wiped with 10% ethanol solution to eliminate olfactory cues. The experimenter exited the room during testing sessions. *<u>Experimental Design and Memory Loads.</u>* The SLR experiment evaluated three memory loads based on spacing between objects 2 and 3: Dissimilar (d-; low memory load, 108° apart), Similar (s-; high memory load, 72° apart), and Extra similar (xs-; very high memory load, 36° apart). During the sample phase (10 min), three identical objects were placed at predetermined distances corresponding to one of the three memory load conditions, and mice freely explored the arena. Following a 35-minute retention interval in home cages, mice were returned for the test phase (5 min) with two objects: one at a familiar location (matching object 1 from the sample phase) and one at a novel location (positioned midway between the original locations of objects 2 and 3) [96]. *<u>Data Analysis</u>*. Time spent in object zones was calculated based on nose position, and movement distance was calculated based on center body position using EthoVision XT 12 (Noldus Information Technology). The discrimination index (d2 ratio) was calculated as

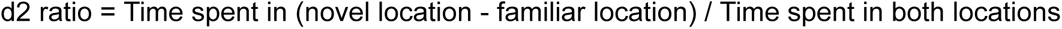

#### Autoshaping

Autoshaping was performed between 8:00 AM and 2:00 PM daily (Monday-Friday) at CHOP using the Bussey-Saksida operant touchscreen platform (Lafayette Life Sciences, cat# 80614A) equipped with the ABET Core Intelli-Interface (Lafayette Life Sciences, cat# 81430) and ABET II Software (Lafayette Life Sciences, cat# 89509)[134,135]. The operant chamber contained a light, auditory cue speaker, food dispenser, and two white vertical rectangular response windows (6.5 × 14 cm) positioned left and right of the reward dispenser. Two infrared photobeams detected approaches to the touchscreen and entries/exits to the food magazine. Behavioral tasks and data collection were controlled by ABET II Autoshaping Software (cat# 89544). The reward stimulus was Strawberry Ensure® Nutrition Shake (Abbott Laboratories) delivered without food deprivation. *<u>Habituation.</u>* Mice were handled for one minute daily for three days prior to habituation. During Hab 1 (one day), mice were placed in the chamber for 10 minutes with all lights off while strawberry milkshake was delivered into the food tray for 2800 ms (70 μl). The number of broken photobeams was recorded to assess locomotor activity. During Hab 2, mice remained in the chamber for 30 minutes while milkshake (280 ms, 7 μl) was delivered after variable intervals (0-30 seconds), accompanied by tray light illumination and a tone. Once reward was delivered, the program waited for the mouse to enter the food tray before restarting the variable interval. Upon tray entry, the tray light was turned off and the procedure repeated. The criterion for Hab 2 was 30/40 trials within the 30-minute session, which most mice achieved within 2 days. *<u>Acquisition Training.</u>* Following habituation, mice underwent 11 days of autoshaping acquisition to associate one side of the screen as a positive conditioned stimulus (CS+) and the other side as a negative conditioned stimulus (CS-), counterbalanced between animals. Trials were conducted in pairs presenting both CS+ and CS- stimuli in random order, ensuring no more than two consecutive presentations of the same stimulus type and preventing the same side from being illuminated first in trial pairs more than 3 times consecutively. After a variable interval (10-40 seconds), the chosen stimulus was presented for 10 seconds when the animal was breaking the rear beam. For CS- trials, another variable interval (10-40 seconds) followed before the other stimulus was presented. For CS+ trials, reward was delivered, tray entry was awaited, then a 10-40 second variable interval preceded the other stimulus presentation. Sessions lasted 30 minutes. The criterion was completing at least 25 trials within the 30-minute session for two out of three consecutive days. *<u>Reversal Learning.</u>* After 11 days of acquisition training, mice underwent reversal learning for 5 (males) or 6 (females) days using identical procedures except CS+ and CS- assignments were switched. The difference in reversal duration between sexes was due to equipment availability. Data (the number of reward chamber approaches, approach difference between CS+ and CS- approach, accuracy to CS+ approaches) were collected by ABET II Software (cat# 89509).

#### Elevated-plus maze

Anxiety behavior in the EPM (Harvard Apparatus, cat# 760075) was assessed 6 mon post-IRR. The EPM apparatus (99cm elevation; 2 open and 2 closed arms each measuring L 67cm x W 6cm, closed arm walls H 17cm) was constructed as described previously [91,93]. Mice were placed in the center of the apparatus pseudorandomly facing one of two open arms, and allowed 5 minutes of free exploration under white lighting conditions (200 lux). Ethovision ver 12 software (Noldus Information Technology) was used to record the time spent in the open arms, closed arms, and center zone, as well as the frequency of entries into each area. From these measurements, an exploration index was calculated:

The ratio of the time spent in open arms vs closed arms:

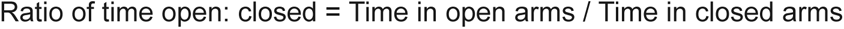

### Fiber photometry recording

A separate *in vivo* imaging cohort (n=7 males, 3 Veh/Sham and 4 Veh/33-GCR mice) received AAV9-CaMKIIa-GCaMP6f viral infusion in the DG hilus and optic fiber implantation in the molecular layer of the DG at 6 months post-IRR to monitor DG granule cell activity as described in Surgery section. This group performed the SLR with fiber photometry (FP) recording at 7.25 months post-IRR. *<u>Recording System and</u> <u>Setup.</u>* Three weeks post-surgery, fiber photometry recordings were conducted using the Neurophotometrics FP3002 system (Neurophotometrics LTD, CA) with customized Bonsai software during the SLR paradigm as previously described [136–138]. A patch cord (Doric Lenses, D204-80052, BBP(2)_200/220/900-0.37_2m_SMA-2xMF1.25) was connected unilaterally to the implanted optic fiber. Calcium-dependent fluorescence changes were recorded at 470 nm excitation, while calcium-independent fluorescence was captured at 415 nm excitation to control for motion artifacts and photobleaching. Data acquisition occurred continuously during both the Sample phase (10 min) and Test phase (5 min) of the SLR paradigm, with simultaneous behavioral video recording. *<u>Behavioral Video Analysis</u>*. Behavioral videos were analyzed using Social LEAP Estimates Animal Poses (SLEAP v1.4.1) for pose estimation [139], followed by Simple Behavioral Analysis (SimBA version 3.2.8 with Python 3.10) for behavioral quantification [140]. For SLEAP model training, 461 frames from 19 videos were annotated, achieving mean Average Precision of 0.854 and mean Average Recall of 0.881. All experimental videos were analyzed using this validated model. SLEAP output files (.csv format) were imported into SimBA for exploratory behavior quantification. Regions of interest were defined as: (1) a 5-cm radius circle centered at the base of each object, and (2) a 29-cm radius circle encompassing the entire arena. Behavioral metrics extracted included time spent in each zone, zone entry counts (calculated relative to nose position), total movement distance, and average velocity (calculated relative to center body position). Frame-by-frame Boolean values indicating zone occupancy were extracted for temporal alignment with photometry data using custom MATLAB code (MathWorks).

Fluorescence changes (ΔF/F) were calculated as previously described [141]:

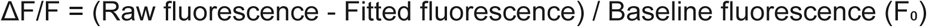

The ΔF/F signals were normalized using Z-score transformation:

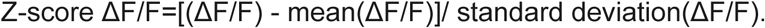

Calcium transient peaks were identified using a noise threshold defined by the Median Absolute Deviation (MAD) method. Peaks were defined as local maxima with: (1) amplitude greater than 0.1×MAD above baseline, and (2) minimum separation of 0 data points between events. Peak frequency and amplitude were quantified from these identified events. For event-locked analysis, ΔF/F traces were aligned to specific behavioral events, including entry into the Familiar Object zone and Novel Object zone during the Test phase of the SLR paradigm.

### Tissue collection and processing

Brain tissue was collected at 7.25 months post-IRR for the behavioral cohort and at 8.75 months post-IRR for the in vivo imaging cohort. Following decapitation, brains were immersed in 4% paraformaldehyde (PFA; Sigma-Aldrich, cat# P6148) in PBS for 3 days [142,143], then cryoprotected by immersion in 30% sucrose (Fisher Scientific, cat# S5-3) containing 0.01% sodium azide (NaN₃; Sigma-Aldrich, cat# S8032) at 4°C for 24 hours until complete equilibration. Brains were coronally sectioned at 40 μm thickness using a freezing microtome (Leica SM2010R). The left hemisphere was marked with a 26-gauge needle on the dorsal cortical area for consistent orientation. Serial sections were collected systematically throughout the hippocampus for stereological assessment as previously reported [94,142,144]. Sectioned tissue was stored in 1× PBS containing 0.01% NaN₃ at 4°C until further processing.

### Immunohistochemistry (IHC)

IHC was performed on slide-mounted coronal brain sections as previously described [91,143]. Sections underwent antigen retrieval by placement in near-boiling citric acid (pH 6.0; Fisher Chemical, cat# A940-500). Endogenous peroxidase activity was quenched by incubation in 0.3% hydrogen peroxide (Sigma, cat# H-1009) in PBS. Non-specific binding was blocked with 3% normal donkey serum (Jackson ImmunoResearch, cat# 017-000-121) in 0.3% Triton X-100 in PBS. Sections were incubated overnight at room temperature with primary antibody against doublecortin (goat anti-DCX; Santa Cruz Biotechnology, cat# SC-8066; 1:500) or GFP (chicken anti-GFP; Aves Labs, cat# GFP-1020; 1:3000) diluted in 3% normal donkey serum with 0.3% Tween-20 in PBS. Following PBS washes, sections were incubated for 1 hour at room temperature with biotinylated secondary antibodies at 1:200 (donkey anti-goat IgG for DCX, Jackson ImmunoResearch, cat# 705-065-003; donkey anti-chicken IgG for GFP, Jackson ImmunoResearch, cat# 703-065-155; 1:200). After additional PBS washes, signal amplification was achieved using avidin-biotin complex (ABC; Vector Laboratories, cat# PK-6100) for HRP conjugation. The HRP signal was visualized using 3,3’-diaminobenzidine (DAB; Thermo Fisher Scientific, cat# 1856090) for DCX and Cy3-conjugated Tyramide Signal Amplification substrate (PerkinElmer, cat# FP1050) for GFP. Nuclei were counterstained with Fast Red (Vector Laboratories, cat# H-3403) for DCX IHC and DAPI (Roche, cat# 236276) for GFP IHC. Sections were dehydrated through a graded ethanol series, cleared in Citrasolv, and coverslipped using DPX mounting medium (Electron Microscopy Services, cat# 13512) with 24×60 mm coverglasses (VWR, cat# 48393).

### Stereological cell counts

DCX-immunoreactive (DCX+) cells were quantified by an observer blinded to experimental conditions using an Olympus BX-51 brightfield microscope at 400× magnification. DCX+ cells in the subgranular zone (SGZ) of the dentate gyrus granule cell layer (GCL) were counted in the right hemisphere of each section, with left hemisphere counted only if right hemisphere damage occurred. Stereological principles were applied as previously described [44,111]. Quantification was conducted along the entire anterior-posterior hippocampal axis (-0.82 to -4.33 mm from bregma).

Two cell classifications were recorded: (1) DCX+ immature neurons, defined as brown-stained soma in the SGZ with neurite and dendrite outgrowth containing at least one dendritic branching node, and (2) DCX+ progenitor cells, defined as brown-stained soma in the SGZ lacking neurite extension.

Total cell populations were calculated using the following stereological formula [145]. The raw cell counts for each bregma level are presented separately.

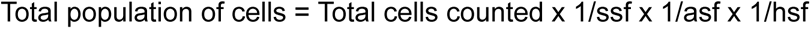

where ssf is the section sampling fraction (1/9 for one hemisphere analysis), asf is the area sampling fraction (1, as all cells were counted in sampled sections), and hsf is the height sampling fraction (1, given minimal edge artifact effects in counting soma <10 μm with ssf 1/18), as described previously [143–145]. Since only one hemisphere was counted, the total population for both hemispheres was calculated as:

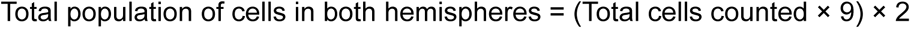

### Principal component analysis

Principal component analysis (PCA) was performed on six behavioral measures: spatial discrimination (low [dSLR] and high [sSLR]: mean d2 ratio), pattern separation (very high [xsSLR]: mean d2 ratio), goal tracking (mean reward chamber approaches, days 4-7), sign tracking (mean CS+ vs. CS- approach difference, days 4-7), and reversal learning (mean CS+ approaches after reversal, days 1-4). For Pavlovian conditioning measures (goal and sign tracking), performance was averaged across days 4-7 of acquisition when animals reached maximal performance. For reversal learning, performance was averaged across days 1-4 post-reversal to capture initial acquisition of the new contingency when cognitive flexibility demands were highest. All measures were z-score normalized across all subjects (grand mean = 0, SD = 1) prior to analysis. PCA was conducted using scikit-learn (version 1.3.0) in Python 3.10 [146]. Component loadings (eigenvectors × √eigenvalues) represent variable-component correlations; loadings >|0.40| were considered substantial. The first three principal components explained 57.6% of total variance (PC1: 20.8%, PC2: 19.7%, PC3: 17.2%). To quantify functional relationships between behavioral domains, vector angles from the three-dimensional loading space (PC1, PC2, PC3) were calculated using the arccosine of normalized dot products. Angles <60° indicated synergistic operations, angles ∼90° indicated functional independence, and angles >135° indicated competitive interactions.

### Computer scripts

Custom analysis scripts were developed in MATLAB and Python 3.10 (Google Colaboratory). A MATLAB script integrated Bonsai Ca²⁺ signal output with SLEAP pose estimation data to calculate zone-specific transient rates and amplitudes during fiber photometry recordings. Python scripts processed 15-min interval photobeam locomotion data and performed principal component analysis (scikit-learn 1.3.0) on multi-domain behavioral measures, including vector angle calculations and visualization. All scripts and documentation are publicly available at https://github.com/EischLab/NSRL22A with installation instructions and example workflows.

### Blinding, subject number, and data removal

All behavioral testing, tissue collection, and data analysis were conducted by investigators blinded to treatment conditions. Two mice were found dead during the study: one female (CDDO-EA/Sham) at 1 month post-IRR prior to data collection, and one male (CDDO-EA/Sham) at 2.5 months post-IRR after home-cage locomotion recording but before SLR testing. These subjects were excluded from all analyses. No additional mice were removed due to husbandry issues or veterinary recommendations. Task-specific subject numbers with exclusion criteria are detailed below and summarized in **S2 Table**. Treatment groups are ordered as Veh/Sham, Veh/33-GCR, CDDO-EA/Sham, and CDDO-EA/33-GCR throughout. Home-cage locomotion recording: One male CDDO-EA/Sham subject was not recorded due to technology failure. Final male subject numbers: n=12, 12, 10, and 12. Final female subject numbers: n=12, 12, 11, and 12. SLR: Subjects were excluded if they failed to meet predetermined criteria during the sample phase (>2 seconds/object, >10 seconds total exploration, equal percent exploration time per object) or test phase (>1 second/object, >5 seconds total exploration). Final male subject numbers: n=10, 10, 9, and 10. Final female subject numbers: n=12, 12, 9, and 11. Autoshaping: Subjects were excluded if they failed to reach acquisition criteria (≥25 trials in 2 out of 3 consecutive days during 11 days of testing). Final male subject numbers: n=10, 11, 10, and 10. Final female subject numbers: n=12, 12, 10, and 11.

### Statistical analysis

Data normality was assessed using D’Agostino-Pearson and Shapiro-Wilk tests with QQ plot visualization. Behavioral measures meeting normality assumptions were analyzed using ANOVA with drug (Veh, CDDO-EA) and irradiation (Sham, 33-GCR) as between-subjects factors. Two-way ANOVA was used for SLR discrimination indices (d2 ratio), SLR test locomotion, EPM measures (time in open arms, entries, exploration index, total distance), and DCX+ total cell counts. Three-way repeated measures ANOVA was used for SLR sample phase object exploration (drug × irradiation × object for objects 1, 2, 3), autoshaping measures (drug × irradiation × day for reward approaches, delta approaches, and percent accuracy during acquisition and reversal), and DCX+ cell distribution across bregma positions (drug × irradiation × bregma). Three-way repeated measures ANOVA was also used for body weight (drug × irradiation × time). Mixed-effects models were used when data were missing. Home cage locomotion (total, ambulatory, and fine beam breaks during both phase and interval testing) failed normality and was analyzed using Kruskal-Wallis tests with Dunn’s post-hoc correction. Fiber photometry data were analyzed using two-way ANOVA (load × irradiation for d2 ratio) or Kruskal-Wallis tests (for signal rate and amplitude measures). Significant ANOVA effects were followed by Tukey’s or Bonferroni post-hoc tests corrected for multiple comparisons. Statistical significance was set at α = 0.05. Data are presented as mean ± SEM. Analyses were performed using R Studio, and Python 3.10 (scikit-learn 1.3.0, NumPy 1.24.3, SciPy 1.11.1, statsmodels 0.14.0).

### Graphs and figures

Graphs were generated using GraphPad Prism 10 (GraphPad Software, San Diego, CA) and Python 3.10 (for PCA analysis). Brightfield photomicrographs were acquired using an Olympus digital camera with cellSens Standard software. Figures were assembled using Adobe Illustrator.

## Data availability statement

All data, including individual animal-level data, necessary to replicate the findings of this study are publicly available. Behavioral data (spontaneous location recognition, autoshaping, reversal learning, elevated plus maze), home cage locomotor activity data, body weight data, fiber photometry recordings, and stereological cell counts have been deposited in Zenodo: https://doi.org/10.5281/zenodo.19157801. Analysis scripts for fiber photometry, locomotor activity, and principal component analysis are available at https://github.com/EischLab/NSRL22A. Statistical output for all analyses is provided in **S1 Table**.

## Supporting Information

**S1 Fig. CDDO-EA reduces locomotor activity across the circadian cycle in both sexes.**

**(A–F)** Locomotor activity measured over 18 hours (4pm–10am) via photobeam breaks in 15-minute bins in males **(A–C)** and females **(D–F)**. Total activity **(A, D)**, ambulatory activity **(B, E)**, and fine activity **(C, F)**. Data are presented as mean ± SEM (n=10–12/group). Statistical analysis: 2-way ANOVA (IRR × Drug) on summed light phase (4pm–7pm, 7am–10am) and dark phase (7pm–7am) data. \**p*<0.05. Post-hoc: Veh/Sham vs CDDO-EA/Sham, ᵇ *p*<0.05, ᵇ’ *p*<0.01; Veh/Sham vs CDDO-EA/33-GCR, ᶜ *p*<0.05; Veh/33-GCR vs CDDO-EA/Sham, ᵈ *p*<0.05; # 0.05<*p*<0.08. Complete subject numbers and detailed statistical analyses are provided in **S1 Table.**

**S1 Table. Full statistical information.**

**S2 Table. Experimental dates and animal omissions.**

## Acknowledgments

We thank the team members at BNL/NSRL for their assistance with the 22A irradiation campaign, particularly Adam Rusek and Peter Guida. We are grateful to an anonymous donor for support of the Eisch Lab and this project. This work was supported by the following funding sources: S.Y. was supported by a 2019 NARSAD Young Investigator Grant from the Brain and Behavior Research Foundation, a 2020 Penn Undergraduate Research Foundation grant, NASA HERO grant 80NSSC21K0814, a 2022 Foerderer Fund for Excellence Award, two CHOP Junior Faculty Awards (2021, PI: Bhoj; 2023, PI: Van Batavia), and NIH awards MH076690 (PI: Tamminga), MH107945 (PI: Eisch), and MH129970 (PI: Eisch). F.K. was supported by the Translational Research Institute for Space Health (TRISH) through NASA cooperative agreement NNX16AO69A, a Penn Provost/CHOP Postdoctoral Fellowship for Academic Diversity, and a Perelman School of Medicine Department of Radiation Oncology Pilot Grant (PIs: Fan and Eisch). A.J.E. was supported by NIH awards MH129970, NS007413, DA007290, DA023555, DA016765, and MH107945; NASA awards NNX07AP84G, NNX12AB55G, and NNX15AE09G; and NIH NS126279 (PI: Ahrens-Nicklas). S.Y. and A.J.E. were also supported by NIH DK135871 (PI: Zderic), NIH NS088555 (PI: Stowe), and NIH MH117628 (PI: Lambert). J.W.S. was supported by NASA awards NNX16AE08G and NNX15AI21G. C.G. and I.S. were supported by PennPREP (R25 GM071745, PI: Jordan-Sciutto). H.H. was supported by the 2021 Penn Undergraduate Research Mentoring Program (PURM), the 2022 Summer Undergraduate Internship Program (SUIP), the 2023 Penn College Alumni Society Board of Managers, a Penn President’s Undergraduate Research Grant, and an augmentation award to NASA HERO grant 80NSSC21K0814 (PI: Yun). A.M. was supported by the Penn Career Services Summer Funding award during Summer 2022 and the Vagelos Molecular Life Sciences Program (2021-2025).

