## Supporting information for "Circuit-selective cognitive vulnerability to environmental stress: multi-domain assessment of space radiation in both sexes reveals countermeasure trade-offs"

S1 Figure O'Connor et al.

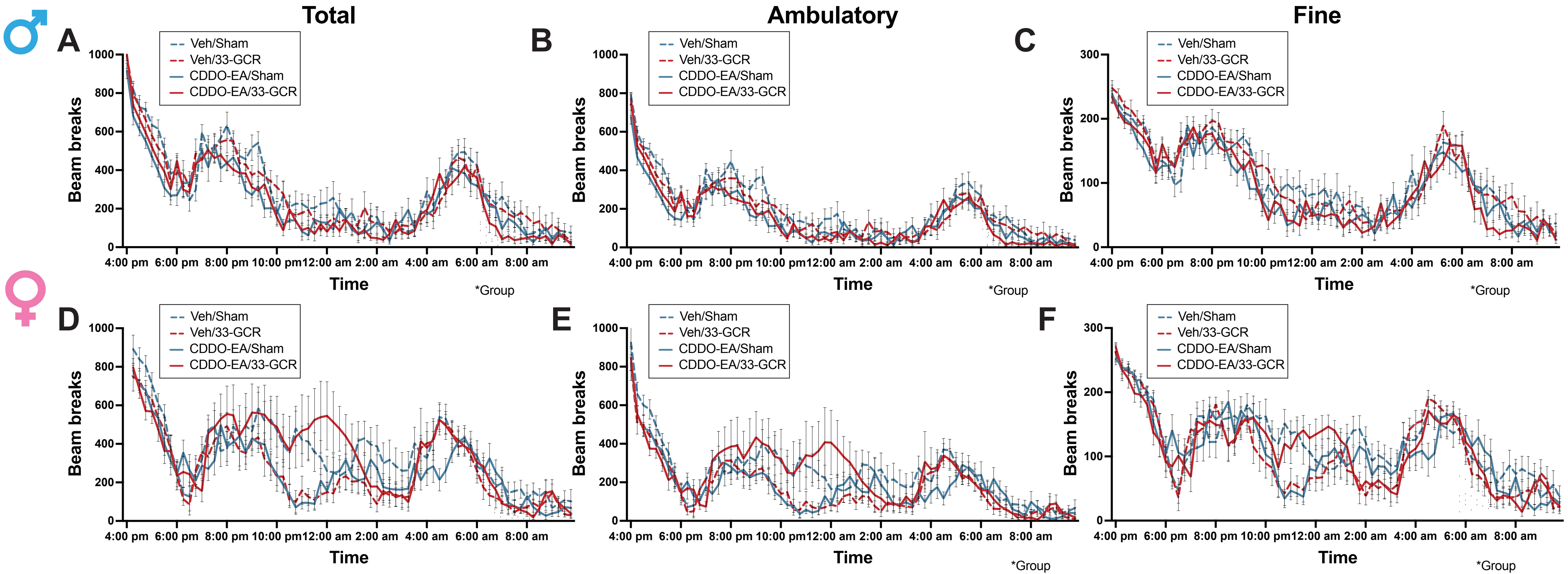

S1 Table. Statistical analysis

| Experiment | Measure | Test Phase | Load (Difficulty) | Figure Panel | Group | Sample Size: n (Event*) | | Mean (predicted or rank) or Median | | | | | | | Passed Normality test? | Test Statistic | Main Effect or Interaction (bold text: P<0.05) | Dist.-Value (Dfn, Dfd) | Main Effect P-Value (bold text: *p*<0.05) | % of Total Variation | Post-hoc Test Correction | Group Difference P-Value (bold text: P<0.05). Lower case italicized letter (e.g. *a*, *b*, *c*, etc) indicates specific group differences. Apostrophe indicates gradations of P values. Post-hoc values of 0.05<P<0.08 are only considered notable (and indicated below and on graph by #) when two additional conditions are met: if effect size is medium or large and if post-hoc 95% CI does not include zero |
|---|---|---|---|---|---|---|---|---|---|---|---|---|---|---|---|---|---|---|---|---|---|---|
| Male d-SLR Sample | % object exploration | -- | d | 2B | | | | *Object 1* | *Object 2* | *Object 3* | | | | | | 3-way RM ANOVA | | | | | Tukey's | |
| | | | | | Veh/Sham | 10 | | 29.944 | 32.957 | 37.1 | | | | | Yes | | **Object** | **F(2, 70)=9.3076** | ***p<0.001*** | | | |
| | | | | | Veh/33-GCR | 10 | | 24.333 | 25.299 | 50.367 | | | | | Yes | | Drug | F(1, 35)= 0.0008 | *p=0.978* | | | |
| | | | | | CDDO-EA/Sham | 9 | | 24.62888889 | 22.96888889 | 52.40111111 | | | | | Yes | | Radiation | F(1, 35)=0.0008 | *p=0.978* | | | |
| | | | | | CDDO-EA/33-GCR | 10 | | 24.554 | 26.952 | 48.495 | | | | | Yes | | Object x Drug | F(2, 70)=0.5637 | *p=0.572* | | | |
| | | | | | | | | | | | | | | | | | Object x Radiation | F(2, 70)= 0.2728 | *p=0.762* | | | |
| | | | | | | | | | | | | | | | | | Drug x Radiation | F(1, 35)=1.0589 | *p=0.311* | | | |
| | | | | | | | | | | | | | | | | | Object x Drug x Radiation | F(2, 70)=0.9425 | *p=0.395* | | | |
| Male d-SLR Test | d2 ratio | -- | d | 2C | Veh/Sham | 10 | | | | | | | | | Yes | 2-way ANOVA | Radiation | F(1, 35)=0.953 | *p=0.336* | | Tukey's | |
| | | | | | Veh/33-GCR | 10 | | | | | | | | | Yes | | Drug | F(1, 35)=0.055 | *p=0.816* | | | |
| | | | | | CDDO-EA/Sham | 9 | | | | | | | | | Yes | | Radiation x Drug | F(1, 35)=0.312 | *p=0.580* | | | |
| | | | | | CDDO-EA/33-GCR | 10 | | | | | | | | | Yes | | | | | | | |
| Male d-SLR Test | Locomotion | -- | d | 2D | Veh/Sham | 10 | | | | | | | | | Yes | 2-way ANOVA | Radiation | F(1, 35)=0.314 | *p=0.579* | | Tukey's | |
| | | | | | Veh/33-GCR | 10 | | | | | | | | | Yes | | Drug | F(1, 35)=0.0005 | *p=0.982* | | | |
| | | | | | CDDO-EA/Sham | 9 | | | | | | | | | Yes | | Radiation x Drug | F(1, 35)=0.314 | *p=0.581* | | | |
| | | | | | CDDO-EA/33-GCR | 10 | | | | | | | | | Yes | | | | | | | |
| Male s-SLR Sample | % object exploration | -- | s | 2F | | | | *Object 1* | *Object 2* | *Object 3* | | | | | | 3-way RM ANOVA | | | | | Tukey's | |
| | | | | | Veh/Sham | 10 | | 28.36 | 39.95 | 31.69 | | | | | Yes | | Object | F(2,70)=1.089 | *p=0.342* | | | |
| | | | | | Veh/33-GCR | 10 | | 36.87 | 30.98 | 32.15 | | | | | Yes | | Drug | F(1, 35)=0.08115 | *p=0.777* | | | |
| | | | | | CDDO-EA/Sham | 9 | | 35.61 | 29.18 | 35.21 | | | | | Yes | | Radiation | F(1, 35)=0.1268 | *p=0.724* | | | |
| | | | | | CDDO-EA/33-GCR | 10 | | 38.06 | 22.83 | 39.11 | | | | | Yes | | **Object x Drug** | **F(2, 70)=3.615** | ***p=0.032*** | | | |
| | | | | | | | | | | | | | | | | | Object x Radiation | F(2, 70)=2.508 | *p=0.089* | | | |
| | | | | | | | | | | | | | | | | | Drug x Radiation | F(1, 35)=2.683 | *p=0.110* | | | |
| | | | | | | | | | | | | | | | | | Object x Drug x Radiation | F(2, 70)=0.3709 | *p=0.692* | | | |
| Male s-SLR Test | d2 ratio | -- | s | 2G | Veh/Sham | 10 | | | | | | | | | Yes | 2-way ANOVA | Radiation | F(1, 35)=0.003 | *p=0.955* | | Tukey's | |
| | | | | | Veh/33-GCR | 10 | | | | | | | | | Yes | | Drug | F(1, 35)=0.483 | *p=0.492* | | | |
| | | | | | CDDO-EA/Sham | 9 | | | | | | | | | Yes | | Radiation x Drug | F(1, 35)=0.060 | *p=0.807* | | | |
| | | | | | CDDO-EA/33-GCR | 10 | | | | | | | | | Yes | | | | | | | |
| Male s-SLR Test | Locomotion | -- | s | 2H | Veh/Sham | 10 | | | | | | | | | Yes | 2-way ANOVA | Radiation | F(1, 35)=0.279 | *p=0.601* | | Tukey's | |
| | | | | | Veh/33-GCR | 10 | | | | | | | | | Yes | | **Drug** | **F(1, 35)= 4.242** | ***p=0.047*** | | | |
| | | | | | CDDO-EA/Sham | 9 | | | | | | | | | Yes | | Radiation x Drug | F(1, 35)=2.078 | *p=0.158* | | | |
| | | | | | CDDO-EA/33-GCR | 10 | | | | | | | | | Yes | | | | | | | |
| Male xs-SLR Sample | % object exploration | -- | xs | 2J | | | | *Object 1* | *Object 2* | *Object 3* | | | | | | 3-way RM ANOVA | | | | | Tukey's | |
| | | | | | Veh/Sham | 10 | | 32.56 | 33.5 | 33.94 | | | | | Yes | | Object | F(2,70)=0.454 | *p=0.637* | | | |
| | | | | | Veh/33-GCR | 10 | | 37.3 | 29.58 | 33.12 | | | | | Yes | | Drug | F(1, 35)= 3.236 | *p=0.081* | | | |
| | | | | | CDDO-EA/Sham | 9 | | 31.3 | 35.57 | 33.13 | | | | | Yes | | **Radiation** | **F(1, 35)= 6.85** | ***p=0.013*** | | | |
| | | | | | CDDO-EA/33-GCR | 10 | | 34.08 | 38.91 | 27.01 | | | | | Yes | | Object x Drug | F(2, 70)= 1.520 | *p=0.226* | | | |
| | | | | | | | | | | | | | | | | | Object x Radiation | F(2, 70)= 0.806 | *p=0.451* | | | |
| | | | | | | | | | | | | | | | | | Drug x Radiation | F(1, 35)= 1.130 | *p=0.295* | | | |
| | | | | | | | | | | | | | | | | | Object x Drug x Radiation | F(1, 70)= 0.6497 | *p=0.525* | | | |
| Male xs-SLR Test | d2 ratio | -- | xs | 2K | Veh/Sham | 10 | | | | | | | -0.1077 | | Yes | 2-way ANOVA | Radiation | F(1, 35)= 1.972 | *p=0.169* | | Tukey's | Veh/GCR vs. Veh/Sham**: a *p<0.05*** |
| | | | | | Veh/33-GCR | 10 | | | | | | | 0.1952 | | Yes | | **Drug** | **F(1, 35)= 4.567** | ***p=0.040*** | | | Veh/GCR vs. CDDO-EA/GCR**: e *p<0.05*** |
| | | | | | CDDO-EA/Sham | 9 | | | | | | | -0.08217 | | Yes | | **Radiation x Drug** | **F(1, 35)= 6.063** | ***p=0.019*** | | | |
| | | | | | CDDO-EA/33-GCR | 10 | | | | | | | -0.165 | | Yes | | | | | | | |
| Male xs-SLR Test | Locomotion | -- | xs | 2L | Veh/Sham | 10 | | | | | | | | | Yes | 2-way ANOVA | Radiation | F(1, 35)=0.008 | *p=0.931* | | Tukey's | |
| | | | | | Veh/33-GCR | 10 | | | | | | | | | Yes | | Drug | F(1, 35)=0.816 | *p=0.373* | | | |
| | | | | | CDDO-EA/Sham | 9 | | | | | | | | | Yes | | Radiation x Drug | F(1, 35)=0.880 | *p=0.355* | | | |
| | | | | | CDDO-EA/33-GCR | 10 | | | | | | | | | Yes | | | | | | | |
| Female d-SLR Sample | % object exploration | -- | d | 3B | | | | *Object 1* | *Object 2* | *Object 3* | | | | | | 3-way RM ANOVA | | | | | Tukey's | |
| | | | | | Veh/Sham | 12 | | 29.88 | 36.93 | 33.19 | | | | | Yes | | **Object** | **F(2, 80)=3.893** | ***p=0.0244*** | | | |
| | | | | | Veh/33-GCR | 12 | | 32.2 | 33.99 | 33.82 | | | | | Yes | | **Drug** | **F(1, 40)=4.390** | ***p=0.0425*** | | | |
| | | | | | CDDO-EA/Sham | 9 | | 31.8 | 37.65 | 30.55 | | | | | Yes | | Radiation | F(1, 40)=0.1031 | *p=0.750* | | | |
| | | | | | CDDO-EA/33-GCR | 11 | | 35.17 | 37.12 | 27.71 | | | | | Yes | | Object x Drug | F(2, 80)=1.900 | *p=0.156* | | | |
| | | | | | | | | | | | | | | | | | Object x Radiation | F(2, 80)=0.8132 | *p=0.447* | | | |
| | | | | | | | | | | | | | | | | | Drug x Radiation | F(1, 40)=0.6308 | *p=0.432* | | | |
| | | | | | | | | | | | | | | | | | Object x Drug x Radiation | F(2, 80)=0.3119 | *p=0.733* | | | |
| Female d-SLR Test | d2 ratio | -- | d | 3C | Veh/Sham | 12 | | | | | | | | | Yes | 2-way ANOVA | Radiation | F(1, 40)=0.088 | *p=0.768* | | Tukey's | |
| | | | | | Veh/33-GCR | 12 | | | | | | | | | Yes | | Drug | F(1, 40)=0.294 | *p=0.591* | | | |
| | | | | | CDDO-EA/Sham | 9 | | | | | | | | | Yes | | Radiation x Drug | F(1, 40)=1.433 | *p=0.238* | | | |
| | | | | | CDDO-EA/33-GCR | 11 | | | | | | | | | Yes | | | | | | | |
| Female d-SLR Test | Locomotion | -- | d | 3D | Veh/Sham | 12 | | | | | | | 2184.418 | | Yes | 2-way ANOVA | **Radiation** | **F(1, 40)= 12.711** | ***p<0.01*** | | Tukey's | Veh/GCR vs. Veh/Sham**: *a p<0.05*** |
| | | | | | Veh/33-GCR | 12 | | | | | | | 1756.65 | | Yes | | **Drug** | **F(1, 40)=4.821** | ***p=0.034*** | | | |
| | | | | | CDDO-EA/Sham | 9 | | | | | | | 1888.934 | | Yes | | Radiation x Drug | F(1, 40)=0.746 | *p=0.393* | | | |
| | | | | | CDDO-EA/33-GCR | 11 | | | | | | | 1628.008 | | No | | | | | | | |
| Female s-SLR Sample | % object exploration | -- | s | 3F | | | | *Object 1* | *Object 2* | *Object 3* | | | | | | 3-way RM ANOVA | | | | | Tukey's | |
| | | | | | Veh/Sham | 12 | | 34.84 | 32.83 | 32.33 | | | | | Yes | | Object | F(2,80)=0.5429 | *p=0.581* | | | |
| | | | | | Veh/33-GCR | 12 | | 33.76 | 30.56 | 35.68 | | | | | Yes | | Drug | F(1, 40)=0.01778 | *p=0.895* | | | |
| | | | | | CDDO-EA/Sham | 9 | | 36.89 | 28.31 | 34.8 | | | | | Yes | | Radiation | F(1, 40)=0.0058 | *p=0.939* | | | |
| | | | | | CDDO-EA/33-GCR | 11 | | 23.86 | 38.59 | 37.55 | | | | | Yes | | Object x Drug | F(2, 80)=0.6754 | *p=0.512* | | | |
| | | | | | | | | | | | | | | | | | Object x Radiation | F(2, 80)=2.188 | *p=0.119* | | | |
| | | | | | | | | | | | | | | | | | Drug x Radiation | F(1, 40)=0.0058 | *p=0.940* | | | |
| | | | | | | | | | | | | | | | | | Object x Drug x Radiation | F(2, 80)=2.191 | *p=0.119* | | | |
| Female s-SLR Test | d2 ratio | -- | s | 3G | Veh/Sham | 12 | | | | | | | | | Yes | 2-way ANOVA | Radiation | F(1, 40)=0.031 | *p=0.861* | | Tukey's | |
| | | | | | Veh/33-GCR | 12 | | | | | | | | | Yes | | Drug | F(1, 40)=0.276 | *p= 0.602* | | | |
| | | | | | CDDO-EA/Sham | 9 | | | | | | | | | Yes | | Radiation x Drug | F(1, 40)=2.613 | *p=0.114* | | | |
| | | | | | CDDO-EA/33-GCR | 11 | | | | | | | | | Yes | | | | | | | |
| Female s-SLR Test | Locomotion | -- | s | 3H | Veh/Sham | 12 | | | | | | | 1668.442 | | Yes | 2-way ANOVA | Radiation | F(1, 40)=3.872 | *p=0.056* | | Tukey's | CDDO/Sham vs. Veh/Sham: ***b p<0.05*** |
| | | | | | Veh/33-GCR | 12 | | | | | | | 1424.134 | | Yes | | **Drug** | **F(1, 40)=9.612** | ***p=0.004*** | | | |
| | | | | | CDDO-EA/Sham | 9 | | | | | | | 1325.551 | | Yes | | Radiation x Drug | F(1, 40)=0.705 | *p=0.406* | | | |
| | | | | | CDDO-EA/33-GCR | 11 | | | | | | | 1227.38 | | No | | | | | | | |
| Female xs-SLR Sample | % object exploration | -- | xs | 3J | | | | *Object 1* | *Object 2* | *Object 3* | | | | | | 3-way RM ANOVA | | | | | Tukey's | |
| | | | | | Veh/Sham | 12 | | 33.13 | 28.92 | 37.95 | | | | | Yes | | Object | F(2, 80)=1.547 | *p=0.219* | | | |
| | | | | | Veh/33-GCR | 12 | | 31.54 | 32.89 | 35.58 | | | | | Yes | | Drug | F(1, 40)=1.143 | *p=0.292* | | | |
| | | | | | CDDO-EA/Sham | 9 | | 33.82 | 28.88 | 37.31 | | | | | Yes | | Radiation | F(1, 40)=2.121 | *p=0.153* | | | |
| | | | | | CDDO-EA/33-GCR | 11 | | 32.48 | 35.38 | 32.14 | | | | | Yes | | Object x Drug | F(2, 80)=0.2584 | *p=0.773* | | | |
| | | | | | | | | | | | | | | | | | Object x Radiation | F(2, 80)=1.788 | *p=0.174* | | | |
| | | | | | | | | | | | | | | | | | Drug x Radiation | F(1, 40)=3.453 | *p=0.071* | | | |
| | | | | | | | | | | | | | | | | | Object x Drug x Radiation | F(2, 80)=0.1457 | *p=0.865* | | | |
| Female xs-SLR Test | d2 ratio | -- | xs | 3K | Veh/Sham | 12 | | | | | | | | | Yes | 2-way ANOVA | Radiation | F(1, 40)=0.971 | *p=0.330* | | Tukey's | |
| | | | | | Veh/33-GCR | 12 | | | | | | | | | Yes | | Drug | F(1, 40)=0.134 | *p=0.716* | | | |
| | | | | | CDDO-EA/Sham | 9 | | | | | | | | | Yes | | Radiation x Drug | F(1, 40)=0.042 | *p=0.839* | | | |
| | | | | | CDDO-EA/33-GCR | 11 | | | | | | | | | Yes | | | | | | | |
| Female xs-SLR Test | Locomotion | -- | xs | 3L | Veh/Sham | 12 | | | | | | | | | Yes | 2-way ANOVA | **Radiation** | **F(1, 40)=5.138** | ***p=0.029*** | | Tukey's | |
| | | | | | Veh/33-GCR | 12 | | | | | | | | | Yes | | Drug | F(1, 40)=2.179 | *p=0.148* | | | |
| | | | | | CDDO-EA/Sham | 9 | | | | | | | | | Yes | | Radiation x Drug | F(1, 40)=1.233 | *p=0.273* | | | |
| | | | | | CDDO-EA/33-GCR | 11 | | | | | | | | | No | | | | | | | |
| Male Animal [illegible] | Weight [illegible] | -- | -- | 4A | | | | **0 Mon post-IRR** | **1 Mon post-IRR** | **2 Mon post-IRR** | **3 Mon post-IRR** | **4 Mon post-IRR** | **6 Mon post-IRR** | **7 Mon post-IRR** | | 3-way Mixed effect RM [illegible] | | | | | Bonferroni | |
| | | | | | Veh/Sham | 12 | | 29.62 | 30.93 | 36.58 | 37.61 | 34.03 | 43.16 | 44.15 | Yes | | Radiation | F(1,43)=0.3024 | *p=0.585* | | | |
| | | | | | Veh/33-GCR | 12 | | 30.08 | 30.39 | 35.49 | 36.36 | 33.4 | 41.66 | 43.28 | Yes | | Drug | F(1,43)=2.8160 | *p=0.101* | | | |
| | | | | | CDDO-EA/Sham | 11-10 | | 29.31 | 30.9 | 36.32 | 37.42 | 34.2 | 43.3 | 46.09 | Yes | | **Time** | **F(6,258)=353.618** | ***p<0.001*** | | | |

| | | | | | | | | | | | | | | | | | | | | | | | | |
|---|---|---|---|---|---|---|---|---|---|---|---|---|---|---|---|---|---|---|---|---|---|---|---|---|
| Weights | (Long-term) | -- | -- | 4A | CDDO-EA/33-GCR | 12 | 30.67 | 32.13 | 38.6 | 39.83 | 35.68 | 45.88 | 46.85 | | | | | Yes | ANOVA | Radiation x Drug | F(1,43)=2.0611 | p=0.158 | Bonferroni | |
| | | | | | | | | | | | | | | | | | | | | Radiation x Time | F(6,258)=0.2278 | p=0.967 | | |
| | | | | | | | | | | | | | | | | | | | | Drug x Time | F(6,258)=1.9330 | p=0.076 | | |
| | | | | | | | | | | | | | | | | | | | | Radiation x Drug x Time | F(6,258)=0.9517 | p=0.459 | | |
| | | | | | | | 0 Mon post-IRR | 1 Mon post-IRR | 2 Mon post-IRR | 3 Mon post-IRR | 4 Mon post-IRR | 6 Mon post-IRR | 7 Mon post-IRR | | | | | | | | | | | |
| Female Animal Weights | Weight (Long-term) | -- | -- | 4B | Veh/Sham | 12 | 22.88 | 22.39 | 24.79 | 28.6 | 31.92 | 30.97 | 33.16 | | | | | Yes | | **Radiation** | **F(1,43)=6.28** | **p=0.016** | | Veh/Sham vs Veh/GCR: **a p<0.05** Months 2, 3, 4, **a' p<0.01** Months 6, 7 |
| | | | | | Veh/33-GCR | 12 | 23.26 | 23.19 | 27.1 | 32.33 | 36.21 | 36.52 | 39.23 | | | | | Yes | | Drug | F(1,43)=0.47 | p=0.495 | Bonferroni | Veh/Sham vs CDDO-EA/Sham: **b' p<0.01** Months 0, 1 |
| | | | | | CDDO-EA/Sham | 11 | 24.41 | 24.33 | 26.12 | 29.42 | 30.95 | 31 | 32.34 | | | | | Yes | 3-way RM ANOVA | **Time** | **F(6,258)=250.06** | **p<0.001** | | |
| | | | | | CDDO-EA/33-GCR | 12 | 23.35 | 23.11 | 25.33 | 29.53 | 34.06 | 34.23 | 36.09 | | | | | Yes | | Radiation x Drug | F(1,43)=1.76 | p=0.192 | | |
| | | | | | | | | | | | | | | | | | | | | **Radiation x Time** | **F(6,258)=12.13** | **p<0.001** | | |
| | | | | | | | | | | | | | | | | | | | | **Drug x Time** | **F(6,258)=3.25** | **p=0.004** | | |
| | | | | | | | | | | | | | | | | | | | | Radiation x Drug x Time | F(6,258)=0.47 | p=0.829 | | |
| | | | | | | | | | | | | Mean beam breaks | | | | | | | | | | | | |
| Male Phase Locomotion - Total | Beam Breaks | -- | -- | 4C | Veh/Sham | 12 | | | | | | 22191.42 | | | | | | No | | | | | | Veh/Sham vs. CDDO/Sham: **b p=0.0446** |
| | | | | | Veh/33-GCR | 12 | | | | | | 20844.58 | | | | | | No | Kruskal-Wallis | Group | df=3 | **p=0.01** | Dunn's test | Veh/GCR vs. CDDO-EA/Sham: **d p=0.0483** |
| | | | | | CDDO-EA/Sham | 10 | | | | | | 17333.8 | | | | | | No | | | | | | Veh/Sham vs. CDDO-EA/GCR: **c# p=0.0560** |
| | | | | | CDDO-EA/33-GCR | 12 | | | | | | 17189.42 | | | | | | No | | | | | | Veh/GCR vs. CDDO-EA/GCR: **e# p=0.0606** |
| | | | | | | | | | | | | Mean beam breaks | | | | | | | | | | | | |
| Male Phase Locomotion - Ambulatory | Beam Breaks | -- | -- | 4D | Veh/Sham | 12 | | | | | | 14406.667 | | | | | | No | | | | | | Veh/Sham vs. CDDO/Sham: **b p=0.0242** |
| | | | | | Veh/33-GCR | 12 | | | | | | 13009.25 | | | | | | No | Kruskal-Wallis | Group | df=3 | **p=0.02** | Dunn's test | Veh/Sham vs. CDDO-EA/GCR: **c p=0.0404** |
| | | | | | CDDO-EA/Sham | 10 | | | | | | 10625.2 | | | | | | No | | | | | | |
| | | | | | CDDO-EA/33-GCR | 12 | | | | | | 10515.083 | | | | | | No | | | | | | |
| | | | | | | | | | | | | Mean beam breaks | | | | | | | | | | | | |
| Male Phase Locomotion - Fine | Beam Breaks | -- | -- | 4E | Veh/Sham | 12 | | | | | | 7784.75 | | | | | | No | | | | | | |
| | | | | | Veh/33-GCR | 12 | | | | | | 7835.33 | | | | | | No | Kruskal-Wallis | Group | df=3 | p=0.11 | | |
| | | | | | CDDO-EA/Sham | 10 | | | | | | 6708.6 | | | | | | No | | | | | | |
| | | | | | CDDO-EA/33-GCR | 12 | | | | | | 6674.33 | | | | | | No | | | | | | |
| | | | | | | | | | | | | Mean beam breaks | | | | | | | | | | | | |
| Female Phase Locomotion - Total | Beam Breaks | -- | -- | 4F | Veh/Sham | 12 | | | | | | 26594.33 | | | | | | No | | | | | | Veh/Sham vs. Veh/GCR: **a p=0.0154** |
| | | | | | Veh/33-GCR | 12 | | | | | | 19679.67 | | | | | | No | Kruskal-Wallis | Group | df = 3 | **p=0.012** | Dunn's test | Veh/Sham vs. CDDO/Sham: **b p=0.0112** |
| | | | | | CDDO-EA/Sham | 11 | | | | | | 19999.64 | | | | | | No | | | | | | |
| | | | | | CDDO-EA/33-GCR | 12 | | | | | | 25288.42 | | | | | | No | | | | | | |
| | | | | | | | | | | | | Mean beam breaks | | | | | | | | | | | | |
| Female Phase Locomotion - Ambulatory | Beam Breaks | -- | -- | 4G | Veh/Sham | 12 | | | | | | 17690.58 | | | | | | No | | | | | | Veh/Sham vs. Veh/GCR: **a p=0.0327** |
| | | | | | Veh/33-GCR | 12 | | | | | | 12189.25 | | | | | | No | Kruskal-Wallis | Group | df=3 | **p=0.019** | Dunn's test | Veh/Sham vs. CDDO/Sham: **b p=0.0127** |
| | | | | | CDDO-EA/Sham | 11 | | | | | | 12113.18 | | | | | | No | | | | | | |
| | | | | | CDDO-EA/33-GCR | 12 | | | | | | 17236.75 | | | | | | No | | | | | | |
| | | | | | | | | | | | | Mean beam breaks | | | | | | | | | | | | |
| Female Phase Locomotion - Fine | Beam Breaks | -- | -- | 4H | Veh/Sham | 12 | | | | | | 8903.75 | | | | | | No | | | | | | Veh/Sham vs. Veh/GCR: **a' p<0.01** |
| | | | | | Veh/33-GCR | 12 | | | | | | 7490.417 | | | | | | No | Kruskal-Wallis | Group | df=3 | **p=0.02** | Dunn's test | Veh/Sham vs. CDDO/Sham: **b# p=0.06** |
| | | | | | CDDO-EA/Sham | 11 | | | | | | 7886.455 | | | | | | No | | | | | | |
| | | | | | CDDO-EA/33-GCR | 12 | | | | | | 8051.667 | | | | | | No | | | | | | |
| | | | | | | | *Day 1* | *Day 2* | *Day 3* | *Day 4* | *Day 5* | *Day 6* | *Day 7* | *Day 8* | *Day 9* | *Day 10* | *Day 11* | | | | | | | |
| Male Autoshaping | Reward approaches (CS+ trials) | Acquisition | -- | 5B | Veh/Sham | 10 | 4 | 2.2 | 2.9 | 4.5 | 7.1 | 2.4 | 3.4 | 5.2 | 3.9 | 2.6 | 2.1 | Yes | | **Time** | **F(10,370)=10.45** | **p<0.0001** | | Veh/GCR: **p=0.0344** Day 4 vs Day 11 |
| | | | | | Veh/33-GCR | 11 | 3.182 | 2.636 | 2.182 | 9.455 | 9.455 | 7.545 | 5.364 | 5 | 5.273 | 3.545 | 1 | Yes | | Radiation | F(1, 37)= 0.4703 | p=0.4971 | | Veh/GCR: **p=0.0344** Day 5 vs Day 11 |
| | | | | | CDDO-EA/Sham | 10 | 2.6 | 4.6 | 4 | 9 | 8.2 | 3.9 | 4.8 | 3.8 | 1.9 | 2.9 | 2.3 | Yes | 3-way RM ANOVA | Drug | F(1, 37)= 1.336 | p=0.2551 | Tukey's | CDDO/SHAM: **p=0.0387** Day 4 vs Day 11 |
| | | | | | CDDO-EA/33-GCR | 10 | 2.2 | 2.8 | 2.9 | 10 | 12.7 | 7.8 | 9.2 | 6.8 | 4.6 | 3.4 | 3.9 | Yes | | Time x Drug | F(10,370)= 1.7339 | p=0.07159 | | CDDO/SHAM: **p=0.0212** Day 4 vs Day 9 |
| | | | | | | | | | | | | | | | | | | | | Time x Radiation | F(10,370)= 0.8985 | p=0.53468 | | |
| | | | | | | | | | | | | | | | | | | | | Drug x Radiation | F(1, 37)= 0.0197 | p=0.8891 | | |
| | | | | | | | | | | | | | | | | | | | | Time x Drug x Radiation | F(10,370)= 0.6270 | p=0.79081 | | |
| | | | | | | | *Day 1* | *Day 2* | *Day 3* | *Day 4* | *Day 5* | *Day 6* | *Day 7* | *Day 8* | *Day 9* | *Day 10* | *Day 11* | | | | | | | |
| Male Autoshaping | Δ Approaches (CS+ vs. CS-) | Acquisition | -- | 5C | Veh/Sham | 10 | -0.7 | -0.1 | -0.6 | -1.7 | 3 | 1.3 | 0.6 | 1.4 | 0.9 | -1 | 1.3 | Yes | | **Time** | **F(10,370)= 5.942** | **p<0.0001** | | |
| | | | | | Veh/33-GCR | 11 | -0.7273 | 0.3636 | -0.5455 | 1 | 3.909 | 3.909 | 2.727 | 3.273 | 4.636 | 3.909 | 3.364 | Yes | | Drug | F(1, 37)= 1.810 | p=0.1867 | | |
| | | | | | CDDO-EA/Sham | 10 | -0.2 | 1.2 | 1.1 | 2.9 | 5.2 | 5.4 | 2.8 | 2.6 | 2.4 | 3.1 | 3.1 | Yes | | Radiation | F(1, 37)= 0.7282 | p=0.3989 | Tukey's | |
| | | | | | CDDO-EA/33-GCR | 10 | -0.3 | 1 | 0.7 | 0.2 | 2.4 | 0.3 | 3.7 | 4.1 | 2.7 | 3.6 | 2.9 | Yes | 3-way RM ANOVA | Time x Drug | F(10,370)= 0.4081 | p=0.9426 | | |
| | | | | | | | | | | | | | | | | | | | | Time x Radiation | F(10,370= 1.299 | p=0.2293 | | |
| | | | | | | | | | | | | | | | | | | | | Drug x Radiation | F(1, 37)= 3.737 | p=0.0609 | | |
| | | | | | | | | | | | | | | | | | | | | Time x Drug x Radiation | F(10, 370= 1.234 | p=0.2674 | | |
| | | | | | | | *Day 1* | *Day 2* | *Day 3* | *Day 4* | *Day 5* | *Day 6* | *Day 7* | *Day 8* | *Day 9* | *Day 10* | *Day 11* | | | | | | | |
| Male Autoshaping | % Accuracy to CS+ approach | Acquisition | -- | 5D | Veh/Sham | 10 | 30.69 | 46.65 | 45.68 | 37.18 | 42 | 41.99 | 44.74 | 47.88 | 41.73 | 34 | 49.33 | Yes | | **Time** | **F(10,370)= 3.261** | **p=0.0005** | | |
| | | | | | Veh/33-GCR | 11 | 35.84 | 36.24 | 41.17 | 43.83 | 44.3 | 47.56 | 48.14 | 52.29 | 54.23 | 53.85 | 66.05 | Yes | | Drug | F(1, 37)= 0.01333 | p=0.9087 | | |
| | | | | | CDDO-EA/Sham | 10 | 28.62 | 33.85 | 44.76 | 54.88 | 52 | 46.07 | 48.19 | 44.11 | 52.42 | 59.38 | 60.92 | Yes | | Radiation | F(1, 37)= 0.0077 | p=0.9305 | Tukey's | |
| | | | | | CDDO-EA/33-GCR | 10 | 39.62 | 42.13 | 41.72 | 34.52 | 48.04 | 36.07 | 48.5 | 38.09 | 45.91 | 51.56 | 48.01 | Yes | 3-way RM ANOVA | Time x Drug | F(10,370)= 0.7706 | p=0.6572 | | |
| | | | | | | | | | | | | | | | | | | | | Time x Radiation | F(10,370)= 0.4470 | p=0.9224 | | |
| | | | | | | | | | | | | | | | | | | | | Drug x Radiation | F(1, 37)= 0.8658 | p=0.3581 | | |
| | | | | | | | | | | | | | | | | | | | | Time x Drug x Radiation | F(10,370)= 1.441 | p=0.160 | | |
| | | | | | | | *Day 1* | *Day 2* | *Day 3* | *Day 4* | *Day 5* | *Day 6* | *Day 7* | *Day 8* | *Day 9* | *Day 10* | *Day 11* | | | | | | | CDDO/Sham vs. Veh/Sham: **b p=0.0164** Day 5 |
| Female Autoshaping | Reward approaches (CS+ trials) | Acquisition | -- | 5E | Veh/Sham | 12 | 1.25 | 2.25 | 2.667 | 4.417 | 3.833 | 2.333 | 1.5 | 2.75 | 1.25 | 1.917 | 1.5 | Yes | | **Time** | **F(10, 410)= 7.0647** | **p<0.0001** | | Veh/Sham vs. Veh/GCR: **a# p=0.0558** Day 5 |
| | | | | | Veh/33-GCR | 12 | 0.6667 | 1.333 | 2 | 4.167 | 6.833 | 4.667 | 4.5 | 5.333 | 3.333 | 0.9167 | 1 | Yes | | **Drug** | **F(1, 41)= 7.3475** | **p= 0.009762** | | CDDO/Sham vs. Veh/Sham: **b p=0.0344** Day 6 |
| | | | | | CDDO-EA/Sham | 10 | 2.4 | 2.8 | 2.8 | 3.9 | 8 | 6.8 | 2.7 | 5 | 3.9 | 2.9 | 3.7 | Yes | | Radiation | F(1, 41)= 2.5751 | p=0.116233 | Tukey's | CDDO/GCR vs. Veh/GCR: **e p=0.0446** Day 6 |
| | | | | | CDDO-EA/33-GCR | 11 | 1.727 | 1.818 | 6.909 | 6 | 7.909 | 13.09 | 8.273 | 4.727 | 7.818 | 3.364 | 3.636 | Yes | 3-way RM ANOVA | Time x Drug | F(10 , 410)= 1.6730 | p= 0.084764 | | CDDO/Sham vs. CDDO/GCR: **f p=0.0492** Day 7 |
| | | | | | | | | | | | | | | | | | | | | **Time x Radiation** | **F(10, 410)= 2.0311** | **p=0.029098** | | Veh/Sham vs. Veh/GCR: **a# p=0.0623** Day 7 |
| | | | | | | | | | | | | | | | | | | | | Drug x Radiation | F(1, 41)= 0.3783 | p=0.541936 | | Veh/Sham vs. Veh/GCR: **a p=0.0249** Day 9 |
| | | | | | | | | | | | | | | | | | | | | Time x Drug x Radiation | F(10, 410)= 0.9068 | p= 0.526739 | | CDDO/GCR vs. Veh/GCR: **e p=0.0466** Day 10 |
| | | | | | | | *Day 1* | *Day 2* | *Day 3* | *Day 4* | *Day 5* | *Day 6* | *Day 7* | *Day 8* | *Day 9* | *Day 10* | *Day 11* | | | | | | | |
| Female Autoshaping | Δ Approaches (CS+ vs. CS-) | Acquisition | -- | 5F | Veh/Sham | 12 | -0.1667 | 0.25 | -2.083 | -0. 1667 | 0 | 1.917 | 3.583 | 1.917 | 1.333 | 3.083 | 2.5 | Yes | | **Time** | **F(10,410)= 4.7645** | **p<0.0001** | | |
| | | | | | Veh/33-GCR | 12 | 0.3333 | -0.5 | -1.083 | -2.667 | -0.25 | 0.5833 | 1 | 1.167 | 1.167 | 1.917 | 3.583 | Yes | | Drug | F(1, 41)= 0.1414 | p=0.7088 | | |
| | | | | | CDDO-EA/Sham | 10 | 1 | -1.9 | 1.2 | 0.6 | 1.9 | 1.9 | 1.1 | 1.6 | 2.8 | 3.3 | 2.2 | Yes | | Radiation | F(1, 41)= 0.9454 | p=0.3366 | Tukey's | |
| | | | | | CDDO-EA/33-GCR | 11 | 0.8182 | -0.4545 | -0.1818 | -0.6364 | 1.273 | 0.9091 | 1.273 | 1.909 | 1.091 | 1.273 | 0.2727 | Yes | 3-way RM ANOVA | Time x Drug | F(10,410)=1.4105 | p=0.1729 | | |
| | | | | | | | | | | | | | | | | | | | | Time x Radiation | F(10,410)=0.4741 | p=0.9066 | | |
| | | | | | | | | | | | | | | | | | | | | Drug x Radiation | F(1, 41)= 0.0064 | p=0.9367 | | |
| | | | | | | | | | | | | | | | | | | | | Time x Drug x Radiation | F(10,410)=0.7709 | p=0.6570 | | |
| | | | | | | | *Day 1* | *Day 2* | *Day 3* | *Day 4* | *Day 5* | *Day 6* | *Day 7* | *Day 8* | *Day 9* | *Day 10* | *Day 11* | | | | | | | |
| Female Autoshaping | % Accuracy to CS+ approach | Acquisition | -- | 5G | Veh/Sham | 12 | 42.18 | 37.74 | 50.94 | 38.92 | 40.5 | 48.43 | 54.4 | 42.16 | 46.71 | 56.58 | 49.75 | Yes | | **Time** | **F(10,410)= 2.4388** | **p=0.007817** | | |
| | | | | | Veh/33-GCR | 12 | 47.7 | 42.64 | 42.67 | 39.43 | 42.39 | 37.59 | 38.14 | 46.71 | 46.16 | 56.53 | 61.54 | Yes | | Drug | F(1, 41)= 1.5502 | p=0.220181 | | |
| | | | | | CDDO-EA/Sham | 10 | 46.83 | 31.91 | 49.83 | 42.18 | 38.41 | 33.13 | 32.46 | 39.55 | 38.95 | 44.64 | 53.07 | Yes | | Radiation | F(1, 41)= 0.0206 | p=0.886528 | Tukey's | |
| | | | | | CDDO-EA/33-GCR | 11 | 50.52 | 37.82 | 51.89 | 36.46 | 40.13 | 34.18 | 44.02 | 44.07 | 31.78 | 37.1 | 36.65 | Yes | 3-way RM ANOVA | Time x Drug | F(6.800, 278.8)= 1.1896 | p=0.295874 | | |
| | | | | | | | | | | | | | | | | | | | | Time x Radiation | F(6.800, 278.8)= 0.4370 | p=0.928075 | | |
| | | | | | | | | | | | | | | | | | | | | Drug x Radiation | F(1, 41)= 0 | p=0.995829 | | |
| | | | | | | | | | | | | | | | | | | | | Time x Drug x Radiation | F(6.800, 278.8)= 1.4199 | p=0.168733 | | |
| | | | | | | | *Day 1* | *Day 2* | *Day 3* | *Day 4* | *Day 5* | | | | | | | | | | | | | |
| Male Autoshaping | Approaches to lit CS+ | Reversal | -- | 6B | Veh/Sham | 10 | 5.3 | 7.4 | 8.4 | 6.9 | 7 | | | | | | | Yes | | **Time** | **F(4, 148)= 3.642** | **p=0.00733** | | |
| | | | | | Veh/33-GCR | 11 | 6 | 5.909 | 6.727 | 5.455 | 7 | | | | | | | Yes | | Drug | F(1, 37)= 3.732 | p=0.0611 | | |
| | | | | | CDDO-EA/Sham | 10 | 5 | 4.7 | 5.5 | 6.3 | 6.3 | | | | | | | Yes | | Radiation | F(1, 37)= 1.149 | p=0.2906 | Tukey's | |
| | | | | | CDDO-EA/33-GCR | 10 | 3.3 | 5.4 | 4.3 | 5.5 | 5 | | | | | | | Yes | 3-way RM ANOVA | Time x Drug | F(4, 148)= 2.006 | p=0.0966 | | |
| | | | | | | | | | | | | | | | | | | | | Time x Radiation | F(4, 148)= 0.5453 | p=0.7027 | | |
| | | | | | | | | | | | | | | | | | | | | Drug x Radiation | F(1, 37)= 0.0026 | p=0.9596 | | |
| | | | | | | | | | | | | | | | | | | | | Time x Drug x Radiation | F(4, 148)= 2.239 | p=0.0676 | | |
| | | | | | | | *Day 1* | *Day 2* | *Day 3* | *Day 4* | *Day 5* | | | | | | | | | | | | | |
| Male Autoshaping | % Accuracy to CS+ approach | Reversal | -- | 6C | Veh/Sham | 10 | 36.41 | 49.33 | 56.83 | 46 | 49 | | | | | | | Yes | | **Time** | **F(4, 148)= 4.364** | **p=0.002299** | | Veh/33-GCR vs. CDDO/33-GCR: **e' p=0.00846** Day 1 |
| | | | | | Veh/33-GCR | 11 | 40.43 | 39.52 | 46.31 | 39.82 | 50.5 | | | | | | | Yes | | **Drug** | **F(1, 37)= 4.803** | **p=0.0348** | | Veh/Sham vs. CDDO/Sham: **b p<0.05** Days 2,3 |
| | | | | | CDDO-EA/Sham | 10 | 33.33 | 31.33 | 36.67 | 42.14 | 42.14 | | | | | | | Yes | | Radiation | F(1, 37)= 0.8976 | p=0.3496 | Tukey's | |
| | | | | | CDDO-EA/33-GCR | 10 | 22.33 | 36 | 28.67 | 38.12 | 34.3 | | | | | | | Yes | 3-way RM ANOVA | Time x Drug | F(4, 148)= 2.011 | p=0.09589 | | |
| | | | | | | | | | | | | | | | | | | | | Time x Radiation | F(4, 148)= 0.4512 | p=0.7714 | | |
| | | | | | | | | | | | | | | | | | | | | Drug x Radiation | F(1, 37)= 0.011 | p=0.9171 | | |
| | | | | | | | | | | | | | | | | | | | | Time x Drug x Radiation | F(4, 148)= 2.034 | p=0.0926 | | |
| | | | | | | | *Day 1* | *Day 2* | *Day 3* | *Day 4* | *Day 5* | | | | | | | | | | | | | |
| | | | | | Veh/Sham | 12 | 5.75 | 6.833 | 7.417 | 7.5 | 9.75 | | | | | | | | | **Time** | **F(4, 164)= 12.61** | **p<0.0001** | | Veh/Sham vs. Veh/33-GCR: **a p=0.0341** Day 6 |

| | | | | | | | | | | | | | | | | | | | | | | | | | | |
|---|---|---|---|---|---|---|---|---|---|---|---|---|---|---|---|---|---|---|---|---|---|---|---|---|---|---|
| Female Autoshaping | Approaches to lit CS+ | Reversal | -- | 6D | Veh/33-GCR | 12 | | 5 | 5.583 | 5.833 | 6.417 | 7.25 | | | | | | | | | 3-way RM ANOVA | Drug<br>Radiation<br>Time x Drug<br>**Time x Radiation**<br>Drug x Radiation<br>Time x Drug x Radiation | F(1, 41)= 0.7074<br>F(1, 41)= 0.9073<br>F(4, 164)= 0.7324<br>**F(4, 164)= 2.645**<br>F(1, 41)= 0.9307<br>F(4, 164)= 1.425 | *p=0.4052*<br>*p=0.3464*<br>*p=0.5711*<br>***p=0.0354***<br>*p=0.3404*<br>*p=0.228* | Tukey's | CDDO/Sham vs. CDDO/33-GCR: ***f p=0.0232*** Day 6 |
| | | | | | CDDO-EA/Sham | 10 | | 4.8 | 4.6 | 5.5 | 7.1 | 8.5 | | | | | | | | | | | | | | |
| | | | | | CDDO-EA/33-GCR | 11 | | 6.273 | 6.455 | 5.182 | 5.182 | 7.455 | | | | | | | | | | | | | | |
| Female Autoshaping | % Accuracy to CS+ approach | Reversal | -- | 6E | | | | ***Day 1*** | ***Day 2*** | ***Day 3*** | ***Day 4*** | ***Day 5*** | | | | | | | | | 3-way RM ANOVA | **Time**<br>Drug<br>Radiation<br>Time x Drug<br>**Time x Radiation**<br>Drug x Radiation<br>Time x Drug x Radiation | **F(4, 164)= 12.41**<br>F(1, 41)= 0.5290<br>F(1, 41)= 1.355<br>F(4, 164)= 0.6330<br>**F(4, 164)= 2.599**<br>F(1, 41)= 0.6011<br>F(4, 164)=1.763 | ***p<0.0001***<br>*p=0.4712*<br>*p=0.2511*<br>*p=0.640*<br>***p=0.03815***<br>*p=0.4426*<br>*p=0.139* | Tukey's | Veh/Sham vs. Veh/33-GCR: ***a p=0.0277*** Day 6<br>CDDO/Sham vs. CDDO/33-GCR: ***f p=0.0462*** Day 4 |
| | | | | | Veh/Sham | 12 | | 38.33 | 47.5 | 49.44 | 50 | 65 | | | | | | | | | | | | | | |
| | | | | | Veh/33-GCR | 12 | | 33.61 | 37.92 | 38.89 | 42.78 | 49.7 | | | | | | | | | | | | | | |
| | | | | | CDDO-EA/Sham | 10 | | 32.41 | 33.42 | 37.83 | 51.9 | 58 | | | | | | | | | | | | | | |
| | | | | | CDDO-EA/33-GCR | 11 | | 42.1 | 43.03 | 34.55 | 34.7 | 49.7 | | | | | | | | | | | | | | |
| Male EPM | Time spent in open arms (s) | -- | -- | 7A | | | | **Mean time** | | | | | | | | | | | | | 2-way ANOVA | Drug<br>Radiation<br>Drug x Radiation | F(1, 43)= 1.387<br>F(1, 43)= 0.163<br>F(1, 43)= 2.417 | *p=0.245*<br>*p=0.688*<br>*p=0.127* | Tukey's | |
| | | | | | Veh/Sham | 12 | | 35.13 | | | | | | | | | | | | Yes | | | | | | |
| | | | | | Veh/33-GCR | 12 | | 45.87 | | | | | | | | | | | | Yes | | | | | | |
| | | | | | CDDO-EA/Sham | 11 | | 37.2 | | | | | | | | | | | | Yes | | | | | | |
| | | | | | CDDO-EA/33-GCR | 12 | | 30.89 | | | | | | | | | | | | Yes | | | | | | |
| Male EPM | Number of entries into open arms | -- | -- | 7B | | | | **Mean number of entries** | | | | | | | | | | | | | 2-way ANOVA | Drug<br>Radiation<br>**Drug x Radiation** | F(1, 43)= 0.345<br>F(1, 43)= 0.005<br>**F(1, 43)= 4.629** | *p=0.560*<br>*p=0.944*<br>***p=0.037*** | Tukey's | |
| | | | | | Veh/Sham | 12 | | 5.5 | | | | | | | | | | | | Yes | | | | | | |
| | | | | | Veh/33-GCR | 12 | | 7.75 | | | | | | | | | | | | Yes | | | | | | |
| | | | | | CDDO-EA/Sham | 11 | | 8.273 | | | | | | | | | | | | Yes | | | | | | |
| | | | | | CDDO-EA/33-GCR | 12 | | 6.167 | | | | | | | | | | | | Yes | | | | | | |
| Male EPM | Ratio time spent in open:closed arms ("Exploration Index") | -- | -- | 7C | | | | **Mean Ratio** | | | | | | | | | | | | | 2-way ANOVA | Drug<br>Radiation<br>Drug x Radiation | F(1, 43)= 1.097<br>F(1, 43)= 0.441<br>F(1, 43)= 2.772 | *p=0.301*<br>*p=0.510*<br>*p=0.103* | Tukey's | |
| | | | | | Veh/Sham | 12 | | 0.1662 | | | | | | | | | | | | Yes | | | | | | |
| | | | | | Veh/33-GCR | 12 | | 0.2584 | | | | | | | | | | | | Yes | | | | | | |
| | | | | | CDDO-EA/Sham | 11 | | 0.1907 | | | | | | | | | | | | Yes | | | | | | |
| | | | | | CDDO-EA/33-GCR | 12 | | 0.151 | | | | | | | | | | | | Yes | | | | | | |
| Male EPM | Total distance traveled | -- | -- | 7D | | | | **Mean distance** | | | | | | | | | | | | | 2-way ANOVA | Drug<br>Radiation<br>Drug x Radiation | F(1, 43)= 0.005<br>F(1, 43)=0.015<br>F(1, 43)= 0.116 | *p=0.945*<br>*p=0.904*<br>*p=0.735* | Tukey's | |
| | | | | | Veh/Sham | 12 | | 1011 | | | | | | | | | | | | Yes | | | | | | |
| | | | | | Veh/33-GCR | 12 | | 1008 | | | | | | | | | | | | Yes | | | | | | |
| | | | | | CDDO-EA/Sham | 11 | | 1005 | | | | | | | | | | | | Yes | | | | | | |
| | | | | | CDDO-EA/33-GCR | 12 | | 1029 | | | | | | | | | | | | Yes | | | | | | |
| Female EPM | Time spent in open arms (s) | -- | -- | 7E | | | | **Mean time** | | | | | | | | | | | | | 2-way ANOVA | Drug<br>Radiation<br>**Drug x Radiation** | F(1, 43)= 0.1555<br>F(1, 43)= 1.774<br>**F(1, 43)= 7.048** | *p=0.695*<br>*p=0.190*<br>***p=0.0111*** | Tukey's | CDDO-EA/Sham vs. CDDO-EA/GCR: ***f p=0.0379*** |
| | | | | | Veh/Sham | 12 | | 52.91 | | | | | | | | | | | | Yes | | | | | | |
| | | | | | Veh/33-GCR | 12 | | 61.91 | | | | | | | | | | | | Yes | | | | | | |
| | | | | | CDDO-EA/Sham | 11 | | 73.65 | | | | | | | | | | | | Yes | | | | | | |
| | | | | | CDDO-EA/33-GCR | 12 | | 46.54 | | | | | | | | | | | | Yes | | | | | | |
| Female EPM | Number of entries into open arms | -- | -- | 7F | | | | **Mean number of entries** | | | | | | | | | | | | | 2-way ANOVA | Drug<br>Radiation<br>**Drug x Radiation** | F(1, 43)= 3.104<br>F(1, 43)= 1.739<br>**F(1, 43)= 10.02** | *p=0.085*<br>*p=0.194*<br>***p=0.003*** | Tukey's | Veh:Sham vs. CDDO-EA:Sham: ***b' p=0.0068***<br>CDDO-EA:Sham vs. CDDO-EA:33-GCR: ***f p=0.0157*** |
| | | | | | Veh/Sham | 12 | | 9.167 | | | | | | | | | | | | Yes | | | | | | |
| | | | | | Veh/33-GCR | 12 | | 11.25 | | | | | | | | | | | | Yes | | | | | | |
| | | | | | CDDO-EA/Sham | 11 | | 14.73 | | | | | | | | | | | | Yes | | | | | | |
| | | | | | CDDO-EA/33-GCR | 12 | | 9.667 | | | | | | | | | | | | Yes | | | | | | |
| Female EPM | Ratio time spent in open:closed arms ("Exploration Index") | -- | -- | 7G | | | | **Mean ratio** | | | | | | | | | | | | | 2-way ANOVA | Drug<br>Radiation<br>**Drug x Radiation** | F(1, 43)= 0.042<br>F(1, 43)= 0.764<br>**F(1, 43)= 5.555** | *p=0.839*<br>*p=0.387*<br>***p=0.023*** | Tukey's | |
| | | | | | Veh/Sham | 12 | | 0.3067 | | | | | | | | | | | | Yes | | | | | | |
| | | | | | Veh/33-GCR | 12 | | 0.3936 | | | | | | | | | | | | Yes | | | | | | |
| | | | | | CDDO-EA/Sham | 11 | | 0.4568 | | | | | | | | | | | | Yes | | | | | | |
| | | | | | CDDO-EA/33-GCR | 12 | | 0.2674 | | | | | | | | | | | | Yes | | | | | | |
| Female EPM | Total distance traveled | -- | -- | 7H | | | | **Mean distance** | | | | | | | | | | | | | 2-way ANOVA | Drug<br>**Radiation**<br>**Drug x Radiation** | F(1, 43)= 0.742<br>**F(1, 43)= 10.36**<br>**F(1, 43)= 5.212** | *p=0.3940*<br>***p=0.002***<br>***p=0.027*** | Tukey's | Veh:33-GCR vs. CDDO-EA:Sham: ***c p=0.0323***<br>CDDO-EA:Sham vs. CDDO-EA:33-GCR: ***f' p=0.0021*** |
| | | | | | Veh/Sham | 12 | | 1333 | | | | | | | | | | | | Yes | | | | | | |
| | | | | | Veh/33-GCR | 12 | | 1275 | | | | | | | | | | | | Yes | | | | | | |
| | | | | | CDDO-EA/Sham | 11 | | 1527 | | | | | | | | | | | | Yes | | | | | | |
| | | | | | CDDO-EA/33-GCR | 12 | | 1187 | | | | | | | | | | | | Yes | | | | | | |
| PCA | PC1 score | -- | -- | 8D | | M | F | **Male** | **Female** | | | | | | | | | | | | 2-way ANOVA | Sex<br>**Treatment**<br>Sex x Treatment | F(1, 74)=2.294<br>**F(3, 74)=3.005**<br>F(3, 74)=0.581 | *p=0.134*<br>***p=0.035***<br>*p=0.629* | Tukey's | Veh:33-GCR vs. CDDO-EA:33-GCR ***p=0.022** |
| | | | | | Veh/Sham | 9 | 12 | -0.390361323 | -0.2995025795 | | | | | | | | | | | Yes | | | | | | |
| | | | | | Veh/33-GCR | 9 | 12 | 0.04779781591 | -0.6394011499 | | | | | | | | | | | Yes | | | | | | |
| | | | | | CDDO-EA/Sham | 9 | 10 | 0.5290672458 | -0.1122118777 | | | | | | | | | | | Yes | | | | | | |
| | | | | | CDDO-EA/33-GCR | 10 | 11 | 0.6373606878 | 0.3942566367 | | | | | | | | | | | No | | | | | | |
| PCA | PC2 score | -- | -- | 8E | Veh/Sham | 9 | 12 | -0.3802076372 | -0.1063386657 | | | | | | | | | | | Yes | 2-way ANOVA | Sex<br>Treatment<br>Sex x Treatment | F(1, 74)=0.191<br>F(3, 74)=1.397<br>F(3, 74)=0.316 | *p=0.663*<br>*p=0.250*<br>*p=0.813* | Tukey's | |
| | | | | | Veh/33-GCR | 9 | 12 | 0.3349109752 | 0.1297192855 | | | | | | | | | | | Yes | | | | | | |
| | | | | | CDDO-EA/Sham | 9 | 10 | 0.3600741083 | 0.2345554561 | | | | | | | | | | | Yes | | | | | | |
| | | | | | CDDO-EA/33-GCR | 10 | 11 | -0.06681617758 | -0.4355415673 | | | | | | | | | | | No | | | | | | |
| PCA | PC3 score | -- | -- | 8F | Veh/Sham | 9 | 12 | -0.3305734888 | -0.5999957053 | | | | | | | | | | | Yes | 2-way ANOVA | Sex<br>**Treatment**<br>Sex x Treatment | F(1, 74)=0.016<br>F(3, 74)=3.990<br>F(3, 74)=0.322 | *p=0.898*<br>***p=0.010***<br>*p=0.808* | Tukey's | |
| | | | | | Veh/33-GCR | 9 | 12 | -0.1008596522 | 0.1206707392 | | | | | | | | | | | Yes | | | | | | |
| | | | | | CDDO-EA/Sham | 9 | 10 | -0.1920146258 | -0.03512035985 | | | | | | | | | | | Yes | | | | | | |
| | | | | | CDDO-EA/33-GCR | 10 | 11 | 0.6644760938 | 0.4608520142 | | | | | | | | | | | No | | | | | | |
| Male FP | d2 ratio | -- | d<br>xs | 9C | | | | D-SLR | xs-SLR | | | | | | | | | | | | 2-way ANOVA | load x Radiation<br>load<br>Radiation | F (1, 5) = 1.437<br>F (1, 5) = 0.7941<br>F (1, 5) = 0.06145 | *p=0.2843*<br>*p=0.4137*<br>*p=0.8141* | | |
| | | | | | Veh/Sham | 3 | | 0.237357897 | -0.1513636432 | | | | | | | | | | | | | | | | | |
| | | | | | Veh/33-GCR | 4 | | 0.1572280179 | 0.2144738396 | | | | | | | | | | | | | | | | | |
| Male FP | Signal Rate-Sample | -- | d | 9E | | | | ***Object 1*** | ***Object 2*** | ***Object 3*** | | | | | | | | | | No | Kruskal-Wallis | | df=6 | *p=0.7974* | Dunn's test | |
| | | | | | Veh/Sham | 3 (124*) | | 3.455 | 3.209 | 3.409 | | | | | | | | | | No | | | | | | |
| | | | | | Veh/33-GCR | 4 (97*) | | 3.093 | 3.061 | 3.464 | | | | | | | | | | | | | | | | |
| Male FP | Signal Amplitude-Sample | -- | d | 9F | | | | ***Object 1*** | ***Object 2*** | ***Object 3*** | | | | | | | | | | No | Kruskal-Wallis | | df=6 | ***p<0.0001*** | | Veh/Sham_Obj 1 vs Veh/Sham_Obj 2 ***p<0.0001***<br>Veh/Sham_Obj 1 vs Veh/Sham_Obj 3 ***p<0.0001***<br>Veh/Sham_Obj 1 vs Veh/GCR_Obj 2 ***p=0.0036***<br>Veh/Sham_Obj 1 vs Veh/GCR_Obj 3 ***p=0.0001***<br>Veh/Sham_Obj 2 vs Veh/GCR_Obj 1 ***p<0.0001***<br>Veh/Sham_Obj 3 vs Veh/GCR_Obj 1 ***p<0.0001***<br>Veh/Sham_Obj 3 vs Veh/GCR_Obj 3 ***p=0.0398***<br>Veh/GCR_Obj 1 vs Veh/GCR_Obj 2 ***p<0.0001***<br>Veh/GCR_Obj 1 vs Veh/GCR_Obj 3 ***p<0.0001*** |
| | | | | | Veh/Sham | 3 (124*) | | 0.01777595479 | -0.00503273650 | -0.00979253521 | | | | | | | | | | No | | | | | | |
| | | | | | Veh/33-GCR | 4 (97*) | | 0.04183 | 0.008583 | 0.002671 | | | | | | | | | | | | | | | | |
| Male FP | Signal Rate-Sample | -- | xs | 9G | | | | ***Object 1*** | ***Object 2*** | ***Object 3*** | | | | | | | | | | No | Kruskal-Wallis | | df=6 | ***p=0.0002*** | | Veh/Sham_Obj 3 vs. Veh/GCR_Obj 2 ***p=0.0005***<br>Veh/Sham_Obj 3 vs. Veh/GCR_Obj 3 ***p=0.0187***<br>Veh/GCR_Obj 1 vs. Veh/GCR_Obj 2 ***p=0.0392*** |
| | | | | | Veh/Sham | 3 (118*) | | 3.29 | 3.164 | 3.714 | | | | | | | | | | No | | | | | | |
| | | | | | Veh/33-GCR | 4(105*) | | 3.35 | 2.416 | 2.653 | | | | | | | | | | | | | | | | |
| Male FP | Signal Amplitude-Sample | -- | xs | 9H | | | | ***Object 1*** | ***Object 2*** | ***Object 3*** | | | | | | | | | | Yes | 2-way ANOVA | Object x Radiation<br>**Object**<br>**Radiation** | F (2, 267) = 0.5401<br>F (2, 267) = 6.681<br>F (1, 267) = 61.07 | *p=0.5834*<br>***p=0.0015***<br>***p<0.0001*** | | ***Veh/Sham vs. Veh/33-GCR p<0.0001 Obj 1,2,3***<br>Veh/GCR_Obj 1 vs. Veh/GCR_Obj 2 ***p=0.0034***<br>Veh/GCR_Obj 1 vs. Veh/GCR_Obj 3 ***p=0.0135*** |
| | | | | | Veh/Sham | 3 (118*) | | 0.02124 | 0.007508 | 0.0069 | | | | | | | | | | Yes | | | | | | |
| | | | | | Veh/33-GCR | 4(105*) | | 0.06576 | 0.03995 | 0.04251 | | | | | | | | | | | | | | | | |
| Male FP | Signal Rate-Test | -- | d | 9I | | | | ***Familiar*** | ***Novel*** | | | | | | | | | | | | Kruskal-Wallis | | df=4 | ***p=0.0011*** | | Veh/Sham_F vs. Veh/GCR_Novel ***p=0.0022***<br>Veh/Sham_N vs. Veh/GCR_Novel ***p=0.0169*** |
| | | | | | Veh/Sham | 3 (53*) | | 4.19 | 3.653 | | | | | | | | | | | | | | | | | |
| | | | | | Veh/33-GCR | 4(23*) | | 2.689 | 2.19 | | | | | | | | | | | | | | | | | |
| Male FP | Signal Amplitude-Test | -- | d | 9J | | | | ***Familiar*** | ***Novel*** | | | | | | | | | | | | Kruskal-Wallis | | df=4 | ***p<0.0001*** | | Veh/Sham_F vs. Veh/GCR_Familiar ***p<0.0001***<br>Veh/Sham_F vs. Veh/GCR_Novel ***p=0.0013***<br>Veh/Sham_N vs. Veh/GCR_Familiar ***p<0.0001*** |
| | | | | | Veh/Sham | 3 (53*) | | 0.04757 | 0.04306 | | | | | | | | | | | | | | | | | |
| | | | | | Veh/33-GCR | 4(23*) | | -0.001335 | 0.01522 | | | | | | | | | | | | | | | | | |
| Male FP | Signal Rate-Test | -- | xs | 9K | | | | ***Familiar*** | ***Novel*** | | | | | | | | | | | | Kruskal-Wallis | | df=4 | *p=0.5420* | | |
| | | | | | Veh/Sham | 3 (34*) | | 3.738 | 2.947 | | | | | | | | | | | | | | | | | |
| | | | | | Veh/33-GCR | 4(57*) | | 3.266 | 3.186 | | | | | | | | | | | | | | | | | |
| Male FP | Signal Amplitude-Test | -- | xs | 9L | | | | ***Familiar*** | ***Novel*** | | | | | | | | | | | | Kruskal-Wallis | | df=4 | ***p=0.0002*** | | Veh/Sham_Novel vs. Veh/GCR_Familiar ***p=0.006***<br>Veh/GCR_Familiar vs. Veh/GCR_Novel ***p=0.0002*** |
| | | | | | Veh/Sham | 3 (34*) | | 0.03284 | 0.04062 | | | | | | | | | | | | | | | | | |
| | | | | | Veh/33-GCR | 4(57*) | | 0.003559 | 0.02044 | | | | | | | | | | | | | | | | | |
| Male DCX | total DCX+ immature neurons | -- | -- | 10B | Veh/Sham | 11 | | 1057 | | | | | | | | | | | | Yes | 2-way ANOVA | Drug<br>Radiation<br>Drug x Radiation | F(1, 37)= 0.406<br>F(1, 37)= 0.755<br>F(1, 37)= 3.052 | *p=0.528*<br>*p=0.39*<br>*p=0.089* | Tukey's | |
| | | | | | Veh/33-GCR | 10 | | 849.6 | | | | | | | | | | | | Yes | | | | | | |
| | | | | | CDDO-EA/Sham | 9 | | 868 | | | | | | | | | | | | Yes | | | | | | |
| | | | | | CDDO-EA/33-GCR | 11 | | 937.6 | | | | | | | | | | | | Yes | | | | | | |
| | | | | | | | | **Bregma -0.82** | **Bregma -1.18** | **Bregma -1.54** | **Bregma -1.9** | **Bregma -2.26** | **Bregma -2.62** | **Bregma -2.98** | **Bregma -3.34** | **Bregma -3.7** | **Bregma -4.06** | **Bregma -4.42** | **Bregma -4.78** | | | | | | | |
| | | | | | Veh/Sham | 11 | | 0 | 0.1818181818 | 6.636363636 | 11.45454545 | 8.545454545 | 7.454545455 | 7 | 6.272727273 | 6.363636364 | 3.909090909 | 0.9090909091 | 0 | Yes | | **Bregma** | **F(11,407)= 47.764** | ***p<0.0001*** | | |

| | | | | | | | | | | | | | | | | | | | | | | | | | | |
|---|---|---|---|---|---|---|---|---|---|---|---|---|---|---|---|---|---|---|---|---|---|---|---|---|---|---|
| | | | | | Veh/33-GCR | 10 | | 0 | 1.3 | 5.3 | 5.9 | 5.8 | 6.9 | 6.3 | 6.4 | 4.6 | 2.6 | 0.8 | 0 | Yes | | Drug | F(1, 37)= 0.2584 | *p=0.6142* | | |
| | | | | | CDDO-EA/Sham | 9 | | 0 | 0.6666666667 | 5 | 8.555555556 | 6.555555556 | 5.888888889 | 8.333333333 | 5.888888889 | 5.222222222 | 2.111111111 | 0 | 0 | Yes | | Radiation | F(1, 37)= 1.114 | *p=0.2980* | | |
| **Male DCX** | **immature neurons x Bregma** | -- | -- | **10C** | CDDO-EA/33-GCR | 11 | | 0 | 1.181818182 | 8.363636364 | 8.727272727 | 7.818181818 | 5.727272727 | 7.363636364 | 8 | 4.454545455 | 0.4545454545 | 0 | 0 | Yes | 3-way RM ANOVA | Bregma x Drug | F(11,407)= 0.888 | *p=0.552* | Tukey's | |
| | | | | | | | | | | | | | | | | | | | | | | Bregma x Radiation | F(11, 407)= 1.372 | *p=0.183* | | |
| | | | | | | | | | | | | | | | | | | | | | | Drug x Radiation | F(1, 37)= 3.871 | *p=0.0570* | | |
| | | | | | | | | | | | | | | | | | | | | | | Bregma x Drug x Radiation | F(11,407)= 1.336 | *p=0.202* | | |
| | | | | | Veh/Sham | 12 | | 1253 | | | | | | | | | | | | Yes | | | | | | |
| **Female DCX** | **total DCX+ immature neurons** | -- | -- | **10D** | Veh/33-GCR | 11 | | 1065 | | | | | | | | | | | | Yes | 2-way ANOVA | Drug | F(1, 37)= 0.550 | *p=0.463* | Tukey's | |
| | | | | | CDDO-EA/Sham | 10 | | 1400 | | | | | | | | | | | | No | | Radiation | F(1, 37)= 4.045 | *p=0.052* | | |
| | | | | | CDDO-EA/33-GCR | 8 | | 1098 | | | | | | | | | | | | Yes | | Drug x Radiation | F(1, 37)= 0.224 | *p=0.639* | | |
| | | | | | | | | **Bregma -0.82** | **Bregma -1.18** | **Bregma -1.54** | **Bregma -1.9** | **Bregma -2.26** | **Bregma -2.62** | **Bregma -2.98** | **Bregma -3.34** | **Bregma -3.7** | **Bregma -4.06** | **Bregma -4.42** | **Bregma -4.78** | | | | | | | |
| | | | | | Veh/Sham | 12 | | 0 | 0.4166666667 | 3.916666667 | 9.583333333 | 9.916666667 | 9.833333333 | 8.916666667 | 9.75 | 8.083333333 | 5.916666667 | 3.25 | 0 | Yes | | **Bregma** | **F(11, 407)= 52.286** | ***p<0.0001*** | | |
| | | | | | Veh/33-GCR | 11 | | 0 | 0.5454545455 | 4.727272727 | 8.363636364 | 11.72727273 | 8.545454545 | 6.727272727 | 8 | 7 | 3.545454545 | 0 | 0 | Yes | | Drug | F(1, 37)= 0.425 | *p=0.518* | | CDDO/Sham vs. CDDO/GCR: ***f p = 0.0353*** Bregma -1.90 |
| | | | | | CDDO-EA/Sham | 10 | | 0 | 0.1 | 6.6 | 12.7 | 11.3 | 10.2 | 11.2 | 10.6 | 8 | 5.7 | 1.4 | 0 | Yes | | **Radiation** | **F(1, 37)= 4.460** | ***p=0.042*** | | |
| **Female DCX** | **immature neurons x Bregma** | -- | -- | **10E** | CDDO-EA/33-GCR | 8 | | 0 | 0.125 | 3.875 | 7.875 | 11.25 | 9.625 | 8.375 | 7.875 | 6.875 | 3.625 | 1 | 0.5 | Yes | 3-way RM ANOVA | Bregma x Drug | F(11,407)= 0.366 | *p=0.968* | Tukey's | |
| | | | | | | | | | | | | | | | | | | | | | | Bregma x Radiation | F(11, 407= 1.435 | *p=0.154* | | |
| | | | | | | | | | | | | | | | | | | | | | | Drug x Radiation | F(1, 37)= 0.322 | *p=0.574* | | |
| | | | | | | | | | | | | | | | | | | | | | | Bregma x Drug x Radiation | F(11, 407)= 0.676 | *p=0.762* | | |
| | | | | | Veh/Sham | 11 | | 3096 | | | | | | | | | | | | Yes | | | | | | |
| **Male DCX** | **total DCX+ progenitor cells** | -- | -- | **10F** | Veh/33-GCR | 10 | | 2707 | | | | | | | | | | | | Yes | 2-way ANOVA | Drug | F(1, 37)= 0.4362 | *p=0.513* | Tukey's | |
| | | | | | CDDO-EA/Sham | 9 | | 2946 | | | | | | | | | | | | Yes | | Radiation | F(1, 37)= 3.368 | *p=0.075* | | |
| | | | | | CDDO-EA/33-GCR | 11 | | 2589 | | | | | | | | | | | | Yes | | Drug x Radiation | F(1, 37)= 0.006 | *p=0.939* | | |
| | | | | | | | | **Bregma -0.82** | **Bregma -1.18** | **Bregma -1.54** | **Bregma -1.9** | **Bregma -2.26** | **Bregma -2.62** | **Bregma -2.98** | **Bregma -3.34** | **Bregma -3.7** | **Bregma -4.06** | **Bregma -4.42** | **Bregma -4.78** | | | | | | | |
| | | | | | Veh/Sham | 11 | | 0 | 1.818181818 | 14 | 28.09090909 | 26.45454545 | 21.27272727 | 21 | 27.90909091 | 22.18181818 | 7 | 2.272727273 | 0 | Yes | | **Bregma** | **F(11, 407)= 76.43** | ***p<0.0001*** | | |
| | | | | | Veh/33-GCR | 10 | | 0 | 2.5 | 13.1 | 20.3 | 19.2 | 18.6 | 18.1 | 22.7 | 20.6 | 7.3 | 3 | 0 | Yes | | Drug | F(1, 37)= 0.201 | *p=0.656* | | |
| | | | | | CDDO-EA/Sham | 9 | | 0 | 1 | 18.88888889 | 26.88888889 | 21.44444444 | 18.77777778 | 21.55555556 | 25 | 19.33333333 | 10.77777778 | 0 | 0 | Yes | 3-way RM ANOVA | **Radiation** | **F(1, 37)= 4.415** | ***p=0.0420*** | | Veh/Sham vs. Veh/GCR: ***a# p=0.0764*** Bregma -1.90 |
| **Male DCX** | **progenitor cells x Bregma** | -- | -- | **10G** | CDDO-EA/33-GCR | 11 | | 0 | 3.636363636 | 18.81818182 | 25.54545455 | 18.36363636 | 15.81818182 | 21.45454545 | 21.54545455 | 15.72727273 | 2.636363636 | 0.2727272727 | 0 | Yes | | Bregma x Drug | F(11,470)= 1.289 | *p=0.228* | Tukey's | CDDO/Sham vs. CDDO/GCR: ***f# p=0.0694*** Bregma -4.06 |
| | | | | | | | | | | | | | | | | | | | | | | Bregma x Radiation | F(11,407)= 0.978 | *p=0.466* | | |
| | | | | | | | | | | | | | | | | | | | | | | Drug x Radiation | F(1, 37)= 0.093 | *p=0.762* | | |
| | | | | | | | | | | | | | | | | | | | | | | Bregma x Drug x Radiation | F(11,407)= 0.628 | *p=0.805* | | |
| | | | | | Veh/Sham | 12 | | 3101 | | | | | | | | | | | | Yes | | | | | | |
| **Female DCX** | **total DCX+ progenitor cells** | -- | -- | **10H** | Veh/33-GCR | 11 | | 2739 | | | | | | | | | | | | Yes | | Drug | F(1, 37)= 1.412 | *p=0.242* | Tukey's | |
| | | | | | CDDO-EA/Sham | 10 | | 3530 | | | | | | | | | | | | No | 2-way ANOVA | **Radiation** | **F(1, 37)= 6.948** | ***p=0.012**** | | |
| | | | | | CDDO-EA/33-GCR | 8 | | 2801 | | | | | | | | | | | | Yes | | Drug x Radiation | F(1, 37)= 0.789 | *p=0.380* | | |
| | | | | | | | | **Bregma -0.82** | **Bregma -1.18** | **Bregma -1.54** | **Bregma -1.9** | **Bregma -2.26** | **Bregma -2.62** | **Bregma -2.98** | **Bregma -3.34** | **Bregma -3.7** | **Bregma -4.06** | **Bregma -4.42** | **Bregma -4.78** | | | | | | | |
| | | | | | Veh/Sham | 12 | | 0 | 0.8333333333 | 13.58333333 | 25 | 24.58333333 | 22 | 22.75 | 24.41666667 | 21.58333333 | 14 | 3.5 | 0 | Yes | | **Bregma** | **F(11,407)= 54.06** | ***p<0.0001*** | | |
| | | | | | Veh/33-GCR | 11 | | 0 | 0.6363636364 | 11.90909091 | 21.36363636 | 23.63636364 | 17.27272727 | 21.09090909 | 21.54545455 | 24.18181818 | 10.36363636 | 0 | 0 | Yes | | Drug | F(1, 37)= 1.345 | *p=0.254* | | CDDO/Sham vs. CDDO/GCR: ***f p = 0.0456*** Bregma -1.90 |
| | | | | | CDDO-EA/Sham | 10 | | 0 | 0.8 | 15.1 | 29.1 | 25.3 | 23.7 | 20.8 | 31.6 | 30.2 | 16.4 | 3.1 | 0 | Yes | | **Radiation** | **F(1, 37)= 7.212** | ***p=0.011*** | | |
| **Female DCX** | **progenitor cells x Bregma** | -- | -- | **10I** | CDDO-EA/33-GCR | 8 | | 0 | 0.125 | 12 | 18 | 22.875 | 21.75 | 20.875 | 23.75 | 21 | 10.5 | 3.875 | 0 | Yes | 3-way RM ANOVA | Bregma x Drug | F(11,407)= 0.339 | *p=0.977* | Tukey's | |
| | | | | | | | | | | | | | | | | | | | | | | Bregma x Radiation | F(11,407)= 0.658 | *p=0.778* | | |
| | | | | | | | | | | | | | | | | | | | | | | Drug x Radiation | F(1, 37)= 0.846 | *p=0.364* | | |
| | | | | | | | | | | | | | | | | | | | | | | Bregma x Drug x Radiation | F(11,407)= 0.607 | *p=0.824* | | |
| | | | | | Veh/Sham | 12 | | | | | | | | | | | | | | Yes | | **Drug** | **F(1, 42)= 8.261** | ***p=0.006*** | | |
| **Male Interval Locomotion - Total Light** | **Beam Breaks** | -- | -- | **Supp 1A** | Veh/33-GCR | 12 | | | | | | | | | | | | | | Yes | 2-way ANOVA | Radiation | F(1, 42)= 0.330 | *p=0.568* | | |
| | | | | | CDDO-EA/Sham | 10 | | | | | | | | | | | | | | Yes | | Drug x Radiation | F(1, 42)= 0.006 | *p=0.940* | | |
| | | | | | CDDO-EA/33-GCR | 12 | | | | | | | | | | | | | | Yes | | | | | | |
| | | | | | Veh/Sham | 12 | | | | | | | | | | | | | | Yes | | **Drug** | **F(1, 42)= 5.043** | ***p=0.030*** | | |
| **Male Interval Locomotion - Total Dark** | **Beam Breaks** | -- | -- | **Supp 1B** | Veh/33-GCR | 12 | | | | | | | | | | | | | | No | 2-way ANOVA | Radiation | F(1, 42)= 0.239 | *p=0.627* | | |
| | | | | | CDDO-EA/Sham | 10 | | | | | | | | | | | | | | Yes | | Drug x Radiation | F(1, 42)= 0.192 | *p=0.663* | | |
| | | | | | CDDO-EA/33-GCR | 12 | | | | | | | | | | | | | | No | | | | | | |
| | | | | | Veh/Sham | 12 | | | | | | | | | | | | | | Yes | | **Drug** | **F(1, 42)= 11.750** | ***p=0.001*** | | |
| **Male Interval Locomotion - Ambulatory Light** | **Beam Breaks** | -- | -- | **Supp 1C** | Veh/33-GCR | 12 | | | | | | | | | | | | | | Yes | 2-way ANOVA | Radiation | F(1, 42)= 0.365 | *p=0.549* | | |
| | | | | | CDDO-EA/Sham | 10 | | | | | | | | | | | | | | Yes | | Drug x Radiation | F(1, 42)= 0.436 | *p=0.513* | | |
| | | | | | CDDO-EA/33-GCR | 12 | | | | | | | | | | | | | | Yes | | | | | | |
| | | | | | Veh/Sham | 12 | | | | | | | | | | | | | | Yes | | **Drug** | **F(1, 42)= 5.544** | ***p=0.023*** | | |
| **Male Interval Locomotion - Ambulatory Dark** | **Beam Breaks** | -- | -- | **Supp 1D** | Veh/33-GCR | 12 | | | | | | | | | | | | | | Yes | 2-way ANOVA | Radiation | F(1, 42)= 0.556 | *p=0.460* | | |
| | | | | | CDDO-EA/Sham | 10 | | | | | | | | | | | | | | No | | Drug x Radiation | F(1, 42)= 0.329 | *p=0.569* | | |
| | | | | | CDDO-EA/33-GCR | 12 | | | | | | | | | | | | | | Yes | | | | | | |
| | | | | | Veh/Sham | 12 | | | | | | | | | | | | | | Yes | | Drug | F(1, 42)= 5.471 | *p= 0.024* | | |
| **Male Interval Locomotion - Fine Light** | **Beam Breaks** | -- | -- | **Supp 1E** | Veh/33-GCR | 12 | | | | | | | | | | | | | | Yes | 2-way ANOVA | Radiation | F(1, 42)= 0.0000988 | *p=0.992* | | |
| | | | | | CDDO-EA/Sham | 10 | | | | | | | | | | | | | | Yes | | Drug x Radiation | F(1, 42)= 0.394 | *p=0.534* | | |
| | | | | | CDDO-EA/33-GCR | 12 | | | | | | | | | | | | | | Yes | | | | | | |
| | | | | | Veh/Sham | 12 | | | | | | | | | | | | | | No | | Drug | F(1, 42)= 3.148 | *p=0.083* | | |
| **Male Interval Locomotion - Fine Dark** | **Beam Breaks** | -- | -- | **Supp 1F** | Veh/33-GCR | 12 | | | | | | | | | | | | | | Yes | 2-way ANOVA | Radiation | F(1, 42)= 0.000614 | *p=0.980* | | |
| | | | | | CDDO-EA/Sham | 10 | | | | | | | | | | | | | | Yes | | Drug x Radiation | F(1, 42)= 0.029000 | *p=0.867* | | |
| | | | | | CDDO-EA/33-GCR | 12 | | | | | | | | | | | | | | Yes | | | | | | |
| | | | | | Veh/Sham | 12 | | | | | | | | | | | | | | Yes | | Drug | F(1, 43)= 1.338 | *p=0.254* | | |
| **Female Interval Locomotion - Total Light** | **Beam Breaks** | -- | -- | **Supp 1G** | Veh/33-GCR | 12 | | | | | | | | | | | | | | Yes | 2-way ANOVA | Radiation | F(1, 43)= 2.865 | *p=0.098* | | |
| | | | | | CDDO-EA/Sham | 11 | | | | | | | | | | | | | | No | | Drug x Radiation | F(1, 43)= 1.210 | *p=0.277* | | |
| | | | | | CDDO-EA/33-GCR | 12 | | | | | | | | | | | | | | Yes | | | | | | |
| | | | | | Veh/Sham | 12 | | | | | | | | | | | | | | Yes | | Drug | F(1, 43)= 0.007 | *p=0.936* | | |
| **Female Interval Locomotion - Total Dark** | **Beam Breaks** | -- | -- | **Supp 1H** | Veh/33-GCR | 12 | | | | | | | | | | | | | | Yes | 2-way ANOVA | Radiation | F(1, 43)= 0.007 | *p=0.935* | | |
| | | | | | CDDO-EA/Sham | 11 | | | | | | | | | | | | | | Yes | | **Drug x Radiation** | **F(1, 43)= 5.561** | ***p=0.023*** | | |
| | | | | | CDDO-EA/33-GCR | 12 | | | | | | | | | | | | | | No | | | | | | |
| | | | | | Veh/Sham | 12 | | | | | | | | | | | | | | Yes | | Drug | F(1, 43)= 1.204 | *p=0.279* | | |
| **Female Interval Locomotion - Ambulatory Light** | **Beam Breaks** | -- | -- | **Supp 1I** | Veh/33-GCR | 12 | | | | | | | | | | | | | | Yes | 2-way ANOVA | Radiation | F(1, 43)= 2.481 | *p=0.123* | | |
| | | | | | CDDO-EA/Sham | 11 | | | | | | | | | | | | | | Yes | | Drug x Radiation | F(1, 43)= 1.507 | *p=0.226* | | |
| | | | | | CDDO-EA/33-GCR | 12 | | | | | | | | | | | | | | Yes | | | | | | |
| | | | | | Veh/Sham | 12 | | | | | | | | | | | | | | Yes | | Drug | F(1, 43)= 0.013 | *p=0.910* | | |
| **Female Interval Locomotion - Ambulatory Dark** | **Beam Breaks** | -- | -- | **Supp 1J** | Veh/33-GCR | 12 | | | | | | | | | | | | | | Yes | 2-way ANOVA | Radiation | F(1, 43)= 0.064 | *p=0.802* | | |
| | | | | | CDDO-EA/Sham | 11 | | | | | | | | | | | | | | Yes | | **Drug x Radiation** | **F(1, 43)= 5.007** | ***p=0.030*** | | |
| | | | | | CDDO-EA/33-GCR | 12 | | | | | | | | | | | | | | No | | | | | | |
| | | | | | Veh/Sham | 12 | | | | | | | | | | | | | | Yes | | Drug | F(1, 43)= 1.214 | *p=0.277* | | |
| **Female Interval Locomotion - Fine Light** | **Beam Breaks** | -- | -- | **Supp 1K** | Veh/33-GCR | 12 | | | | | | | | | | | | | | Yes | 2-way ANOVA | Radiation | F(1, 43)= 2.888 | *p=0.096* | | |
| | | | | | CDDO-EA/Sham | 11 | | | | | | | | | | | | | | No | | Drug x Radiation | F(1, 43)= 0.275 | *p=0.603* | | |
| | | | | | CDDO-EA/33-GCR | 12 | | | | | | | | | | | | | | Yes | | | | | | |
| | | | | | Veh/Sham | 12 | | | | | | | | | | | | | | Yes | | Drug | F(1, 43)= 0.024 | *p=0.877* | | |
| **Female Interval Locomotion - Fine Dark** | **Beam Breaks** | -- | -- | **Supp 1L** | Veh/33-GCR | 12 | | | | | | | | | | | | | | Yes | 2-way ANOVA | Radiation | F(1, 43)= 1.193 | *p=0.281* | | |
| | | | | | CDDO-EA/Sham | 11 | | | | | | | | | | | | | | Yes | | **Drug x Radiation** | **F(1, 43)= 4.859** | ***p=0.033*** | | |
| | | | | | CDDO-EA/33-GCR | 12 | | | | | | | | | | | | | | Yes | | | | | | |

S2. Table. Sample omission

| Experiment | Figure | Sex | Group | Number of animals omitted from data analysis | Animal IDs omitted | Reason for omission | n/group | n/sex | n total | Criteria for omission |
|---|---|---|---|---|---|---|---|---|---|---|
| Overall | 1 | Male | Veh/Sham | 0 | NA | NA | 12 | 47 | 94 | Live vs. died |
| | | | Veh/33-GCR | 0 | NA | NA | 12 | | | |
| | | | CDDO/Sham | 1 | 9 | Died after locomotion - but excluded from all stats | 11 | | | |
| | | | CDDO/33-GCR | 0 | NA | NA | 12 | | | |
| | 1 | Female | Veh/Sham | 0 | NA | NA | 12 | 47 | | |
| | | | Veh/33-GCR | 0 | NA | NA | 12 | | | |
| | | | CDDO/Sham | 1 | 28 | Died before locomotion | 11 | | | |
| | | | CDDO/33-GCR | 0 | NA | NA | 12 | | | |
| Spontaneous Location Recognition (SLR) | 2 | Male | Veh/Sham | 2 | 91, 92 | Did not reach either test or sample criteria for inclusion | 10 | 39 | 83 | **Sample** criteria: animals spent >2 sec/obj, >10 sec total exploration, and did not display unequal exploration (<3%/obj)<br><br>**Test** Criteria: animals spent >1sec/obj, >5 sec total exploration |
| | | | Veh/33-GCR | 2 | 28, 41 | | 10 | | | |
| | | | CDDO/Sham | 2 | 47, 76, 9 (died) | | 9 | | | |
| | | | CDDO/33-GCR | 2 | 49, 50 | | 10 | | | |
| | 3 | Female | Veh/Sham | 0 | NA | NA | 12 | 44 | | |
| | | | Veh/33-GCR | 0 | NA | NA | 12 | | | |
| | | | CDDO/Sham | 2 | 25, 26, 28 (died) | Did not reach either test or sample criteria for inclusion | 9 | | | |
| | | | CDDO/33-GCR | 1 | 29 | | 11 | | | |
| Locomotion | 4 | Male | Veh/Sham | 0 | NA | NA | 12 | 46 | 93 | Recording integrity |
| | | | Veh/33-GCR | 0 | NA | NA | 12 | | | |
| | | | CDDO/Sham | 1 | 10, 9 (died) | Recording malfunction | 10 | | | |
| | | | CDDO/33-GCR | 0 | NA | NA | 12 | | | |
| | 4 | Female | Veh/Sham | 0 | NA | NA | 12 | 47 | | |
| | | | Veh/33-GCR | 0 | NA | NA | 12 | | | |
| | | | CDDO/Sham | 0 | 28 (died) | NA | 11 | | | |
| | | | CDDO/33-GCR | 0 | NA | NA | 12 | | | |
| Autoshaping | 5 | Male | Veh/Sham | 2 | 91, 94 | Did not reach acquisition criteria for inclusion | 10 | 41 | 86 | Each animal has to reach at least **25** trials in **2 out of 3** consecutive days (out of 11 total days) of acquisition training. |
| | | | Veh/33-GCR | 1 | 25 | | 11 | | | |
| | | | CDDO/Sham | 1 | 76, 9 (died) | | 10 | | | |
| | | | CDDO/33-GCR | 2 | 16, 50 | | 10 | | | |
| | 6 | Female | Veh/Sham | 0 | NA | | 12 | 45 | | |
| | | | Veh/33-GCR | 0 | NA | | 12 | | | |
| | | | CDDO/Sham | 1 | 12, 28 (died) | Did not reach acquisition criteria for inclusion | 10 | | | |
| | | | CDDO/33-GCR | 1 | 31 | | 11 | | | |
| Elevated Plus Maze (EPM) | 7 | Male | Veh/Sham | 0 | NA | NA | 12 | 47 | 94 | Total distance traveled (cm) outlier test |
| | | | Veh/33-GCR | 0 | NA | NA | 12 | | | |
| | | | CDDO/Sham | 0 | 9 (died) | NA | 11 | | | |
| | | | CDDO/33-GCR | 0 | NA | NA | 12 | | | |
| | 7 | Female | Veh/Sham | 0 | NA | NA | 12 | 47 | | |
| | | | Veh/33-GCR | 0 | NA | NA | 12 | | | |
| | | | CDDO/Sham | 0 | 28 (died) | NA | 11 | | | |
| | | | CDDO/33-GCR | 0 | NA | NA | 12 | | | |
| Double-Cortin X (DCX) | 10 | Male | Veh/Sham | 1 | 3 | Not able to count DCX cells | 11 | 43 | 84 | Damage to L **and** R dentate gyrus (DG) in at least one countable brain section, or outlier on statistics |
| | | | Veh/33-GCR | 2 | 5, 8 | | 10 | | | |
| | | | CDDO/Sham | 2 | 75, 12, 9 (died) | | 9 | | | |
| | | | CDDO/33-GCR | 1 | 81 | | 11 | | | |
| | 10 | Female | Veh/Sham | 0 | NA | NA | 12 | 41 | | |
| | | | Veh/33-GCR | 1 | 22 | Not able to count DCX cells | 11 | | | |
| | | | CDDO/Sham | 1 | 11, 28 (died) | | 10 | | | |
| | | | CDDO/33-GCR | 4 | 16, 29, 46, 48 | | 8 | | | |
